# Stochastic Character Mapping of Continuous Traits on Phylogenies

**DOI:** 10.1101/2024.08.12.607655

**Authors:** B. S. Martin, M. G. Weber

**Affiliations:** Department of Plant Biology, Ecology, Evolution, and Behavior Program, Michigan State University, East Lansing, MI 48824, USA; Department of Plant Biology, University of Georgia, Athens, GA 30602, USA; Department of Ecology and Evolutionary Biology, University of Michigan, Ann Arbor, MI 48109, USA

**Keywords:** eucalypts, macroevolution, comparative methods, stochastic character mapping, ancestral state reconstruction, continuous trait evolution, Brownian Motion, rate heterogeneity

## Abstract

Fossilized organisms only represent a small fraction of Earth’s evolutionary history, motivating “ancestral state reconstruction” techniques for inferring unobserved phenotypes of evolving lineages based on measurements of their relatives. Stochastic character mapping has emerged as a particularly powerful approach in this regard, allowing researchers to sample histories of discrete variables on phylogenies and better account for the inherent uncertainty of reconstructed ancestral states. Here, we generalize this procedure to work with continuous variables by developing an efficient algorithm for sampling evolutionary histories under Brownian Motion, implemented in a new R package *contsimmap*. To demonstrate potential applications of these “*continuous* stochastic character maps”, we develop a pipeline for inferring relationships between rates of continuous trait evolution and continuously-varying factors (e.g., body size, generation time)—a difficult statistical problem for which relatively few methods are available. After verifying this novel pipeline’s performance on simulated data, we use it to show that smaller eucalypt trees tend to exhibit higher rates of flower and leaf trait evolution overall, aligning with well-established predictions based on life history theory as well as empirical patterns in other systems. Ultimately, continuous stochastic character mapping is a valuable new tool for analyzing macroevolutionary data, enabling rigorous yet flexible investigation of complex evolutionary dynamics involving continuous traits and/or continuous variables hypothesized to affect evolutionary processes.

---

Understanding phenotypic evolution over deep time is central to many fundamental questions in evolutionary biology. Accounting for the evolutionary history of particular phenotypes is necessary to comprehensively address questions ranging from “what trait-trait or trait-environment correlations are present across the tree of life?” to “which factors constrain or promote trait and lineage diversity?”. Unfortunately, directly observing phenotypic evolution over macroevolutionary timescales is often impossible due to the incompleteness of the fossil record. Accordingly, biologists have developed numerous “indirect” approaches to instead infer these largely unobserved evolutionary processes from comparative phenotypic data. Perhaps the oldest and most explicit approach to this task is “ancestral state reconstruction” (ASR)—that is, estimating the unobserved phenotypes of evolving lineages based on the observed phenotypes of their (typically living) relatives (Dobzhansky and Sturtevant, 1938; Sanger et al., 1955; Pauling et al., 1963; Witmer, 1995; Schultz et al., 1996; Sumrall and Brochu, 2008).

With the development of rigorous statistical methods for performing ASR under various phylogenetic comparative models of trait evolution, ASR has become ubiquitous in macroevolutionary research (Schluter et al., 1997; Groussin et al., 2016; Joy et al., 2016). Stochastic character mapping in particular has become an extremely popular ASR technique, allowing researchers to randomly sample evolutionary histories of characters along phylogenies according to their probability under a given model of trait evolution (Nielsen, 2002; Huelsenbeck et al., 2003; Bollback, 2006). This approach is generally more useful than other ASR techniques because, while the most likely estimates of ancestral phenotypes under a given trait evolution model are often inaccurate (particularly among older lineages without accompanying fossil data; e.g., Slater et al., 2012; Joy et al., 2016), they are typically so uncertain as to be consistent with a wide variety of reasonable evolutionary trajectories. By sampling hundreds or thousands of stochastic character maps, researchers can flexibly conduct macroevolutionary analyses over distributions of possible evolutionary histories and better account for the inherent uncertainty of reconstructed ancestral states in their inferences. Accordingly, stochastic character mapping has been used across numerous studies for a variety of purposes, from determining the timing/frequency of evolutionary events (e.g., Baliga and Law, 2016; Tornabene et al., 2016; Freyman and Höhna, 2019; Hughes et al., 2021; Landis et al., 2021; Siqueira et al., 2023) to investigating correlations between past life history/environmental factors and evolutionary outcomes (e.g., de Alencar et al., 2017; Borstein et al., 2019; Burns and Bloom, 2020; Fabre et al., 2020; Rincon-Sandoval et al., 2020; Drury et al., 2021; Nations et al., 2021; Friedman and Muñoz, 2023).

While stochastic character mapping has revolutionized statistical approaches to macroevolutionary analysis, all currently available implementations are limited to use on discrete variables, with no analogue for continuous variables. This conspicuous methodological gap constrains approaches for analyzing macroevolutionary data, as many methods for testing whether evolutionary dynamics vary according to some “explanatory factor” (e.g., habitat, reproductive strategy) explicitly rely on fixed estimates of the factor’s evolutionary history—often provided in the form of stochastic character maps (e.g., Revell, 2013a; Clavel et al., 2015; Beaulieu and O’Meara, 2023). Consequently, researchers investigating the interplay between evolutionary dynamics and continuously-varying factors are generally forced to arbitrarily divide continuous variation into discrete categories and/or develop custom statistical pipelines tailored to their specific research goals (Cooper and Purvis, 2009; Hansen et al., 2008; Welch and Waxman, 2008; FitzJohn, 2010; Lartillot and Poujol, 2011; Felsenstein, 2012; Baker et al., 2015; Weir and Lawson, 2015; Clavel and Morlon, 2017; Harvey and Rabosky, 2017; Cooney and Thomas, 2021; Hansen et al., 2022; Uyeda et al., 2021; Boyko et al., 2023a; Tribble et al., 2023). Given that many biologically important variables are continuous in nature (e.g., body size, generation time, climatic niche; Friedman et al., 2019; Gingerich, 2001; Tribble et al., 2023), the lack of stochastic character mapping methods for continuous variables currently presents a limiting hurdle for comparative research.

Here, we introduce an efficient and flexible method for stochastic character mapping of continuous variables under Brownian Motion models of evolution. These “*continuous* stochastic character maps” may be used to better incorporate uncertainty in analyses that aim to characterize the evolutionary dynamics of continuous traits, elucidate the timing/phylogenetic locations of major transitions in continuously-measured phenotypes, and/or explore how continuously-varying factors affect evolutionary dynamics, among other common goals in macroevolutionary research. We first present an algorithm for generating continuous stochastic character maps that works by sampling values of continuous variables across phylogenies while conditioning on any observed data at the phylogeny’s nodes/tips. Second, we showcase potential uses of continuous stochastic character mapping by developing a general pipeline for inferring relationships between continuous trait evolution dynamics and continuously-varying factors based on mapped factor histories—focusing on factor-associated evolutionary rate variation in particular. Lastly, after verifying this pipeline’s performance on simulated data, we use it to show that lineages of smaller eucalypt trees tend to exhibit higher rates of leaf and flower trait evolution overall, demonstrating the empirical utility of both our pipeline as well as continuous stochastic character mapping in general.

## Materials and Methods

Continuous stochastic character mapping is a flexible framework for sampling evolutionary histories of continuous variables along phylogenies under Brownian Motion models. We implement this framework in an R package called *contsimmap*, which supports stochastic character mapping of one or more potentially correlated variables while also accommodating multiple measurements per tip/node with associated intraspecific variation/measurement error (Felsenstein, 2008; Hansen and Bartoszek, 2012; Landis and Schraiber, 2017), missing measurements, and even among-lineage variation in evolutionary trends (Hansen and Martins, 1996) and/or rates. The R package additionally provides tools for transforming, summarizing, and visualizing continuous stochastic character maps (Fig. 1), as well as fitting Brownian Motion models with parameters dependent on mapped continuous variables.

**Figure 1.**
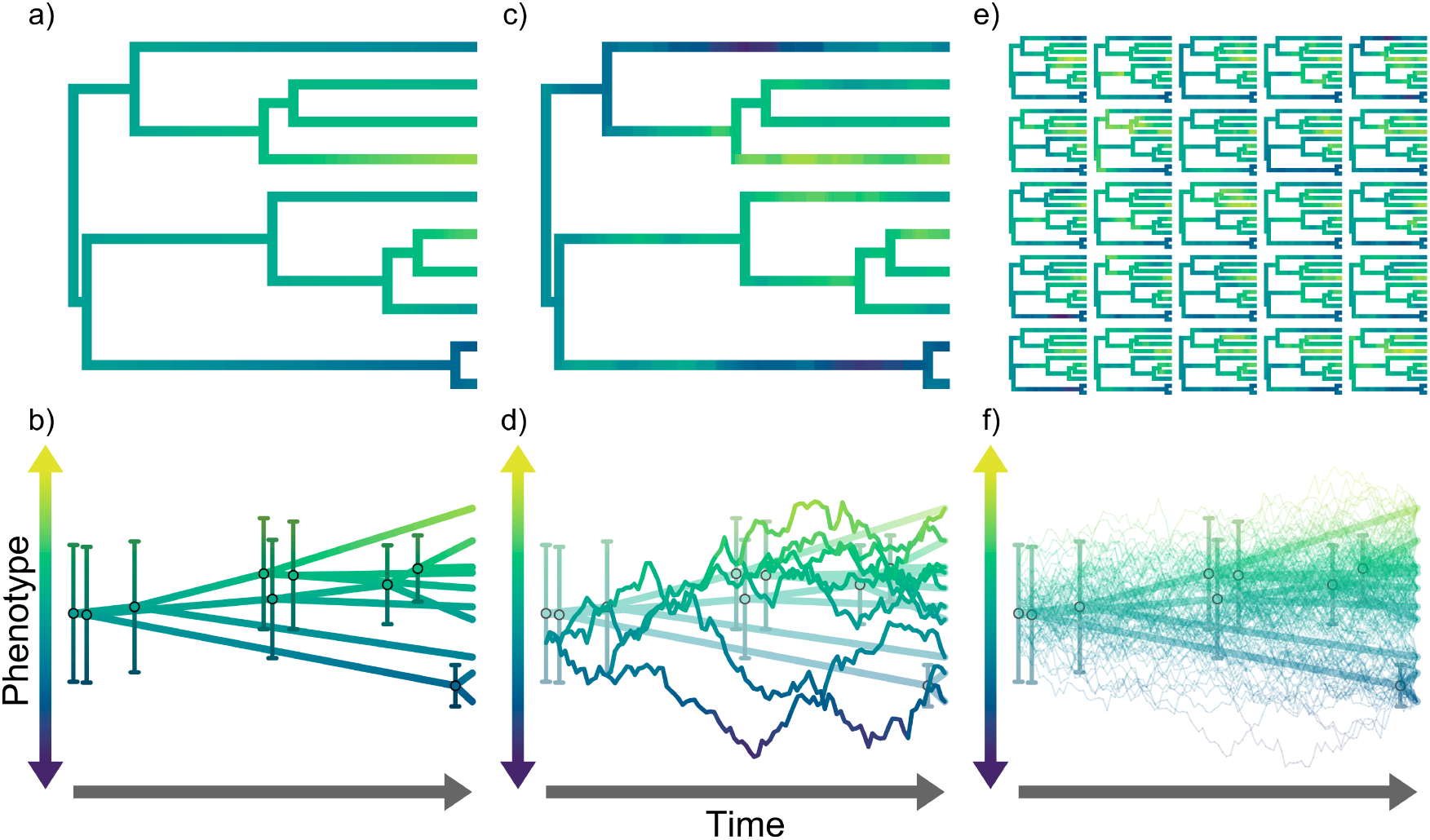
Phylogram and phenogram-based (Revell, 2013b) visualizations of continuous stochastic character maps, colored according to reconstructed phenotypic values. Panels a-b: conventional ancestral state reconstructions of a continuously-varying phenotype under a Brownian Motion model. The phylogram in panel a is colored according to mean phenotypic estimates, which are also depicted by the phenogram in panel b along with error bars representing 95% confidence intervals on the estimates at each node. Panels c-d: a single continuous stochastic character map generated using the same data and Brownian Motion model, again represented as both a colored phylogram and phenogram in panels c and d, respectively. Panels e-f: a sample of 25 continuous stochastic character maps representing the overall distribution of generated maps also depicted as phylograms and phenograms in panels e and f, respectively. Panels c-f also include conventional ancestral state reconstructions for reference.

### Overview of Continuous Stochastic Character Mapping

In this section, we provide an overview of our algorithm for generating continuous stochastic character maps. For simplicity, the explanations in this section generally assume we are sampling the evolutionary history of a single continuous variable (hereafter “phenotype”) under a homogeneous Brownian Motion model based on exact measurements at the tips of a phylogeny. A more in-depth and technical description of the algorithm—including how it is modified to work with multiple correlated variables, among-lineage variation in evolutionary trends/rates, and “messier” comparative data involving multiple/missing measurements at both tips and internal nodes—can be found in the Online Appendix (available under Supplementary Material on bioRxiv).

Overall, our algorithm works by sampling phenotypic values at fine-grained time points (including all nodes) across a phylogeny while conditioning on observed measurements at tips. This procedure is analogous to conventional discrete stochastic character mapping, but with two key distinctions. First, discrete stochastic character mapping generally samples evolutionary histories under continuous time Markov chain models, whereas our method uses Brownian Motion. Second, while discrete stochastic character mapping works by directly mapping evolutionary transitions among different states onto a phylogeny, continuous stochastic character mapping instead samples phenotypic values at a finite number of time points evenly distributed across a phylogeny. The overall density of time points is controlled by a user-specified “resolution” parameter corresponding to the approximate number of points along the longest root-to-tip path in the phylogeny. Critically, sampling phenotypic values along edges allows researchers to account for uncertainty in phenotypic evolutionary trajectories between nodes, which increases with edge length. Such uncertainty is completely obscured by conventional methods for ASR of continuous traits, which generally report estimated phenotypes at nodes only.

Our algorithm ensures sampled phenotypic values follow the joint distribution of unobserved phenotypic states across all time points under an assumed Brownian Motion model (i.e., it explicitly accounts for expected correlations among unobserved phenotypes of closely-related lineages). For computational efficiency, we avoid sampling phenotypic values from all time points at once by recursively computing and sampling from distributions of phenotypic values for one point at a time. To do this, our algorithm employs a “backward then forwards” traversal approach, similarly to other ancestral state reconstruction algorithms (e.g., Yang, 2006; Hiscott et al., 2016; Goolsby, 2017; Hassler et al., 2022). This procedure consists of first calculating distributions of possible phenotypic values at each node during a backwards traversal across the phylogeny from its tips to the root, followed by a forwards (i.e., root to tips) traversal that progressively updates and samples from the distributions at both nodes and time points along edges. We visually illustrate this process in Fig. 2 and describe each of these traversals in more detail below:

**Figure 2.**
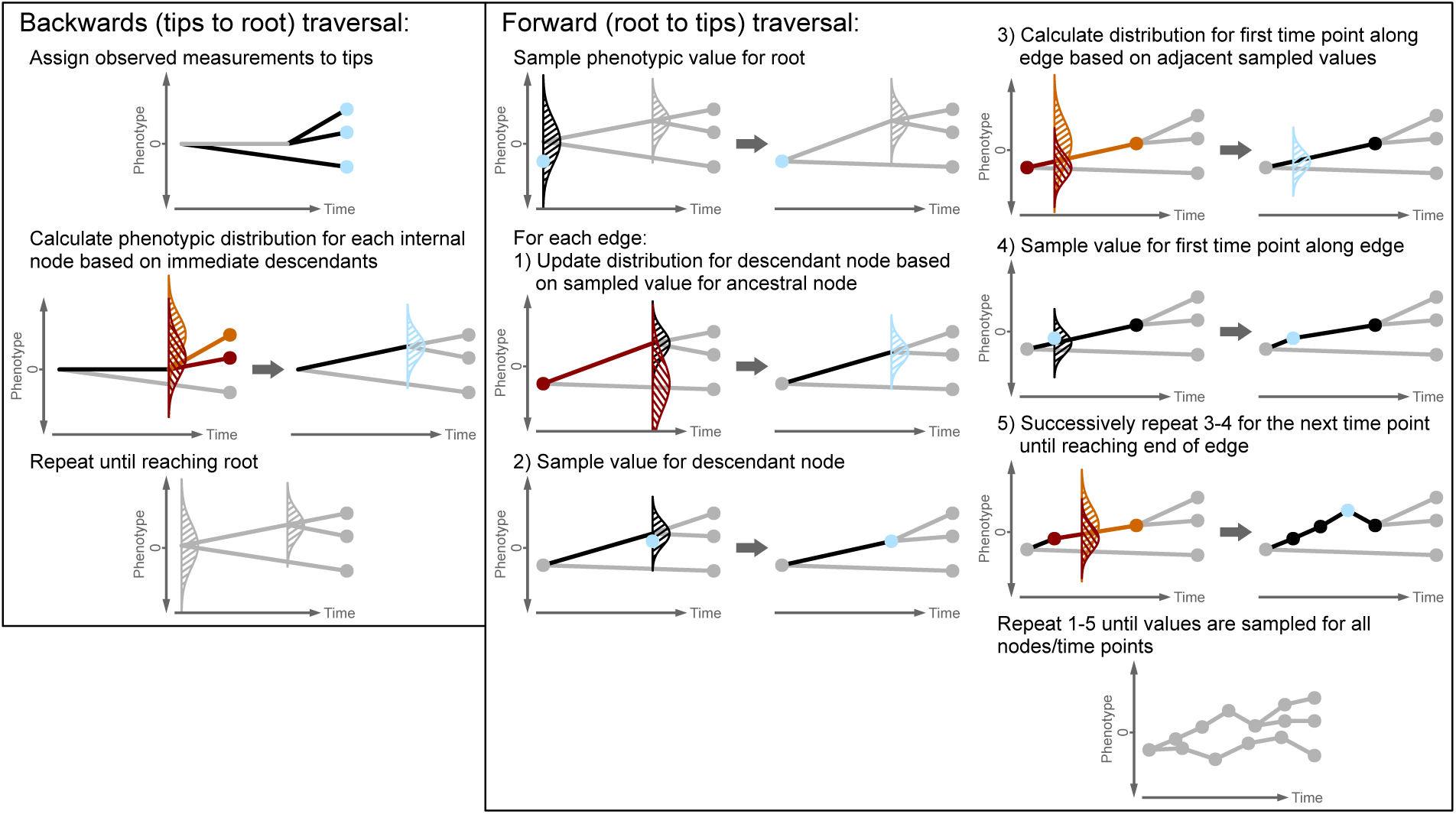
Visual phenogram-based (Revell, 2013b) overview of the major steps in our continuous stochastic character mapping algorithm. Line-shaded, vertically-oriented normal distributions represent distributions of possible phenotypic values at particular points on the phylogeny, while dots indicate exact phenotypic values (either measured or sampled). The algorithm generally focuses on one edge in a given step, which—along with any associated distributions/dots—are drawn in black, though light blue is used to highlight particular distributions/dots on the focal edge that are updated in a given step. Medium-dark red/orange are also used to indicate distributions briefly used in intermediate calculations—these distributions are conditioned on particular nodes/edges drawn in matching colors. Note that, for purely aesthetic purposes, phenotypic distributions at internal nodes are initialized at 0 while values at time points along edges are set to their maximum likelihood expectation before being explicitly sampled (i.e., a straight line connecting adjacent nodes). Line-shaded normal distributions and dots are not drawn at these points to convey that proper phenotypic distributions are yet to be calculated/sampled from.

#### Backwards traversal

This first traversal calculates phenotypic distributions at all nodes in the phylogeny based only on the measurements associated with each node’s descendants. After assigning observed measurements to their corresponding tips, the algorithm visits each internal node in the phylogeny, making sure to only visit a given node after all of its direct descendants have been visited. By leveraging convenient properties of Brownian Motion and normal distributions (see Online Appendix for specific details), the phenotypic distribution for each node is computed recursively from the distributions associated with the node’s immediate descendants. After reaching and calculating the phenotypic distribution at the root of the phylogeny, the algorithm moves on to the subsequent forwards traversal.

#### Forwards traversal

The second traversal has two main functions: updating phenotypic distributions at each node based on measurements associated with each node’s *non*-descendants and sampling exact phenotypic values for all nodes/time points across the phylogeny. The algorithm begins by sampling a phenotypic value for the root from its associated distribution (i.e., the one calculated during the backwards traversal). Next, the algorithm visits each node, first updating the node’s phenotypic distribution based on the value sampled for its direct ancestor. Then, the algorithm samples a value for the given node from this updated distribution. Before moving on to another node, the algorithm samples phenotypic values at equally-spaced time points along the given node’s preceding edge. The phenotypic distributions at these time points are derived by conditioning the assumed Brownian Motion process to start and end at the sampled phenotypic values for each point’s immediate ancestor and the node descending from the given edge (i.e., a “Brownian bridge”; Karatzas and Shreve, 2004; see also Horvilleur and Lartillot, 2014; Lartillot et al., 2016; Quintero and Landis, 2020; Jhwueng, 2021; Quintero, 2025).

Upon completing the forward traversal, the algorithm generates a single continuous stochastic character map. From here, the process can be repeated to sample additional maps in proportion to their probability under the assumed Brownian Motion model.

### Validation Study

To verify that our continuous stochastic character mapping algorithm correctly generates samples from the joint distribution of unobserved phenotypic states under an assumed Brownian Motion model, we conducted a validation study comparing the results of our algorithm to their theoretical expectations. To derive the expected distribution of phenotypic values, we first consider the joint distribution of both observed measurements and phenotypic values at all time points across the phylogeny. Because these values are multivariate normally distributed under Brownian Motion models, we can use well-established formulae to compute the corresponding conditional distribution of phenotypic values given exact values for all observed measurements (Petersen and Pedersen, 2012; Jhwueng, 2021; see Online Appendix for details). We used one-sample Kolmogorov-Smirnov tests (Marsaglia et al., 2003) to determine whether the continuous stochastic character maps generated by our algorithm follow these expected distributions (using Mahalanobis distances and *χ*^2^ distributions as univariate summaries of these multivariate samples/distributions; see Online Appendix).

To comprehensively validate our algorithm, we generated continuous stochastic character maps and calculated expected distributions under a variety of simulated scenarios. In total, we defined 360 distinct simulation conditions that encompassed trait datasets simulated on both ultrametric and non-ultrametric phylogenies ranging from 10-100 tips with or without evolutionary trends, intraspecific variation/measurement error, multiple potentially correlated phenotypes, multiple observations per tip, and missing data (See Online Appendix for specific details). For half of these conditions, we generated continuous stochastic character maps and calculated expected distributions under the same Brownian Motion models used to simulate data. For the other half, we instead generated maps/calculated expected distributions under Brownian Motion models with random parameters (i.e., micmicking parameter misestimation/model misspecificiation). We simulated 10 phylogenies and associated trait datasets per condition for a total of 3,600 simulations, generating 100 continuous stochastic character maps (ranging in resolution from 20–500) per simulated dataset.

### Pipeline using Continuous Stochastic Character Maps to Model Factor-Dependent Trait Evolution

Stochastic character maps are particularly useful for “sequential inference” pipelines, whereby the impact of an explanatory factor on an evolutionary process is inferred by first generating stochastic character maps of factor histories, then fitting factor-dependent evolutionary models based on these maps. For discrete stochastic character maps, evolutionary models can be made factor-dependent by simply allowing the model’s parameters—such as evolutionary rates or trends in the case of Brownian Motion models—to vary among lineages in different discrete states (e.g., lineages in mountain versus lowland habitats, annual versus perennial plant lineages; see Revell, 2013a for further description of this approach). For continuous stochastic character maps, with an infinite continuum of states, such models must instead use “parameter functions” that convert factor values to parameter values (e.g., the value of a parameter could increase/decrease according to a linear or exponential relationship with the factor; e.g., FitzJohn, 2010).

We implemented a set of tools in our R package for sequential inference of factor-dependent Brownian Motion models based on continuous stochastic character maps of factor histories (Fig. 3). Specifically, our implementation constructs a likelihood function for a trait dataset given a set of continuous stochastic character maps (step 1 in Fig. 3) along with parameter functions that convert these mapped values to the evolutionary rate, trend, and/or intraspecific/measurement error variance parameters of a Brownian Motion model (step 2 in Fig. 3). The parameter functions are themselves controlled by free parameters determining the overall magnitude of rates, trends, etc. and/or how they vary with respect to factor values, analogously to intercepts and slopes in linear regression. Thus, fitting parameter functions to data involves inferring the values of these free parameters via maximum likelihood or Bayesian approaches. Given estimates for free parameters, a likelihood function constructed using our R package automatically transforms the given set of factor histories into sets of parameter histories—assuming the factor/parameter values at each time point are constant during their preceding time intervals—and uses the pruning algorithm outlined by Hassler et al., 2022 to calculate likelihoods conditional on each continuous stochastic character map (step 3 in Fig. 3). Ultimately, the likelihood function returns a single overall likelihood by averaging all the conditional likelihoods (in practice, we sometimes found it beneficial to account for “outlier” conditional likelihoods before taking these averages; see Online Appendix). This can be viewed as a Monte Carlo approach to estimating the “true” likelihood that would be obtained by integrating over all possible factor histories rather than a finite sample of them (Mayrose and Otto, 2011).

**Figure 3.**
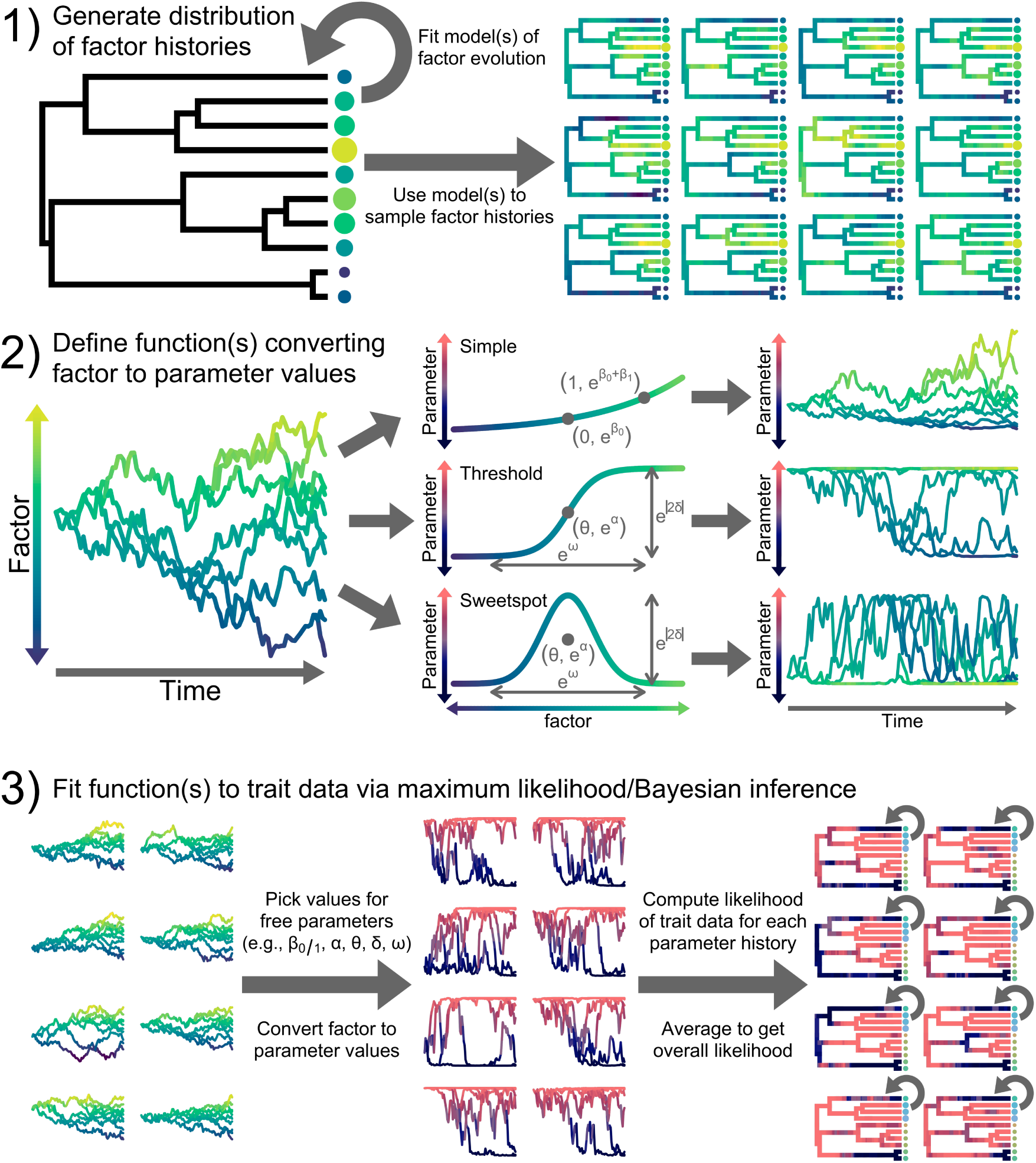
Visual overview of our continuous stochastic character map-based pipeline for inferring relationships between continuously-varying factors and the parameters of a Brownian Motion model of trait evolution. The pipeline consists of three major steps: 1) fitting Brownian Motion models to factor data and using continuous stochastic character mapping to generate a distribution of possible factor histories, 2) defining parameter functions that convert mapped factor values to parameter values, and 3) fitting parameter functions to trait data by using maximum likelihood/Bayesian approaches to infer values of free parameters defining parameter functions. The lighter purple-yellow and darker blue-red color gradients indicate different factor and parameter values, respectively. Differently sized/colored dots arrayed along the tips of phylogenies indicate observed factor and trait data in steps 1 and 3, respectively. Note in step 2 that we illustrate the three main parameter functions (along with their associated free parameters) used to model factor-rate relationships in the current work.

Our R package currently provides tools for using the C++ library NLOPT (Johnson, 2021), interfaced through the R package *nloptr* (Ypma et al., 2022), to infer free parameter values that maximize a given likelihood function. While our implementation allows users to select any optimization algorithm available in the NLOPT library and customize optimizer settings as they see fit, we generally recommend the (currently default) SBPLX (Rowan, 1990) algorithm due to its flexibility and robustness under many conditions.

### Pipeline Simulation Study

To assess the performance of our pipeline for inferring relationships between continuous trait evolution dynamics and continuously-varying factors, we tested whether our approach could be used to reliably detect and quantify factor-dependent rates of continuous trait evolution from simulated data. We simulated the evolution of a single continuous trait, *Y*, with rates depending on a simulated continuous factor—either an observed factor *X_o_* or unobserved factor *X_h_*—and applied our pipeline to the simulated data to infer relationships between rates of *Y* evolution and *X_o_*. In an empirical context, *X_h_* represents a “hidden” factor that affected rates of trait evolution and may thus mislead hypothesis testing for factor-dependent rates (May and Moore, 2020; see also Beaulieu and O’Meara, 2016; Boyko and Beaulieu, 2023; Boyko et al., 2023b; Tribble et al., 2023). After outlining our simulation procedure below, we outline our analysis procedure, which includes a pragmatic technique for constructing additional null models that help account for hidden factors and thereby mitigate their potential effects on hypothesis testing.

We allowed rates of *Y* evolution, *σ*^2^, to vary with factor values (denoted *X* below) according to one of four parameter functions: 1) “simple” functions whereby rates exponentially increase/decrease, 2) “threshold” functions whereby rates shift between some minimum and maximum value, 3) “sweetspot” functions whereby rates peak or dip around a particular factor value, and 4) “constant” functions whereby rates do not vary at all. We define simple functions as:

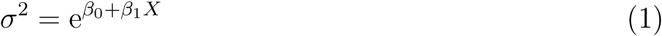

Where the free parameters *β*_0_ and *β*_1_ represent the intercept and slope of the factor-rate relationship on a logarithmic scale (see step 2 in Fig. 3 for a visual representation). Simple functions are useful in scenarios where rates are hypothesized to increase or decrease in association with a factor like temperature (Clavel and Morlon, 2017; Slater et al., 2017) or generation time (Gingerich, 2001). We use an exponential—as opposed to linear—relationship to ensure estimated rates are always positive while also better reflecting empirical findings that evolutionary rates tend to vary multiplicatively rather than additively (Limpert et al., 2001; Gingerich, 2009).

Threshold functions are useful in cases where rates are hypothesized to follow a logistic relationship with some factor. A benefit of threshold functions is that, unlike simple functions, they are bounded between minimum and maximum rates. Under simple functions, rates of trait evolution can become arbitrarily close to 0 and grow without bound. In reality, various factors like mutation rates, integration among traits, and/or physical limits to trait variation will constrain how slow or fast rates can be. We define threshold functions as:

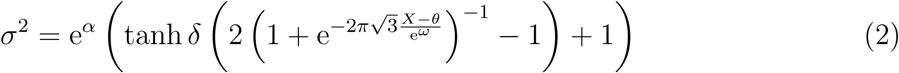

Here, *α* alters the overall scale of rates, specifically corresponding to the natural log of the rate halfway between the minimum and maximum possible rates or “mid rate”, while *θ* determines the factor value or “location” at which rates shift (i.e., the value of *X* at which *σ*^2^ = e*^α^*). We denote *δ*, which controls the direction and magnitude of the rate shift, the “rate deviation”. The fold-difference between minimum and maximum rates is given by e^2|^*^δ|^*, and positive and negative values of *δ* yield upward and downward shifts with increasing factor values, respectively. Lastly, *ω* adjusts the “width” of the shift, with rates roughly reaching their minimum/maximum values at factor values of *θ* ± e*^ω^/*2 (see step 2 in Fig. 3 for a visual representation of these free parameters).

Sweetspot functions can be used when rates are hypothesized to peak or dip at intermediate factor values (Cooper and Purvis, 2009; FitzJohn et al., 2009; Feldman et al., 2016; Amado et al., 2021). For example, Cooper and Purvis (2009) showed that rates of cranial shape evolution are slowest among Paltyrrhine monkeys and Palangeriforme possums of intermediate size, increasing among both smaller and especially larger-bodied lineages. Under sweetspot functions, factor-rate relationships follow positively-valued Gaussian curves of arbitrary height and orientation:

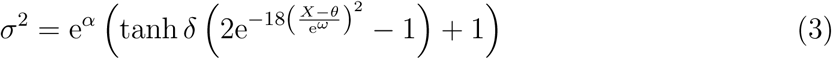

Where *α*, *θ*, *δ*, and *ω* largely have the same effects and interpretations as they do for the threshold function (see step 2 in Fig. 3 for a visual representation). Under the sweetspot function, however, rates peak to their maximum at *θ* and reach their minimum at factor values around *θ* ± e*^ω^/*2 if *δ* is positive. Conversely, if *δ* is negative, rates will instead dip to their minimum at *θ* and reach their maximum around *θ* ± e*^ω^/*2.

Lastly, we define constant functions, whereby rates do not vary, as *σ*^2^ = e*^α^* for consistency with the *α* parameter under threshold and sweetspot functions.

For each simulation, we used *phytools* (Revell, 2012) to simulate an ultrametric, pure-birth phylogeny of height 1 with either 50, 100, or 200 tips. To simulate the continuous factors *X_o_* and *X_h_*, we generated two densely-sampled continuous factor histories (i.e., resolution of 500) evolving under Brownian Motion processes with root values of 0, no trends, and constant rates of 4. For the trait *Y*, we simulated an additional Brownian Motion process with root value 0 and no trends, but with rates varying according one of 12 possible factor-rate relationships which differed in type (i.e., constant, simple, threshold, or sweetspot), whether they depended on the observed factor *X_o_* or hidden factor *X_h_*, the overall magnitude of rate variation or “strength”, and how gradually rates change with respect to factor values or “width” (see Table S1 for the specific relationships/free parameter values used). To assess the robustness of our approach to imperfect correspondences between factors and rates, we repeated these simulations while adding random variation or noise around factor-rate relationships in a manner inspired by both uncorrelated relaxed clock models and rate fluctuations under Lévy processes (e.g., Drummond et al., 2006; Landis et al., 2013; Lartillot et al., 2016; see Online Appendix for details on noise simulation). Ultimately, we defined 72 distinct simulation conditions and repeated our simulation procedure 120 times per condition, yielding a grand total of 8,640 simulated phylogenies and associated factor/trait datasets.

For the analysis procedure, we retained the phylogeny and tip factor/trait values of *X_o_* and *Y* from each simulation. We analyzed the factor/trait datasets by first fitting Brownian Motion models to the *X_o_* data assuming no intraspecific variation/measurement error or evolutionary trends and using the fitted model to generate 100 continuous stochastic character maps of *X_o_* with resolutions of 100 (e.g., step 1 of Fig. 3). We then fit four Brownian Motion models to the *Y* data conditioned on these maps (again assuming no intraspecific variation/measurement error or evolutionary trends)—three non-null models assuming rates depend on mapped *X_o_* values through either a simple, threshold, or sweetspot function, plus a null model assuming a constant rate (e.g., steps 2 and 3 in Fig. 3). To better account for rate variation due to hidden factors, we fit three additional null models to the *Y* data assuming rates depend on a mapped “dummy factor” *D*—which is simulated under the same Brownian Motion model fitted to the *X_o_* data—through either a simple, threshold, or sweetspot relationship. Because *D* exhibits the same evolutionary dynamics as *X_o_* but with random tip values, support for *D*-dependent over *X_o_*-dependent models provide evidence that rates of *Y* evolution vary, but in a way not necessarily related to the observed factor *X_o_* (see also Tribble et al., 2023, which employs a similar approach in testing for relationships between discrete trait evolution and continuous factors). See the Online Appendix for details on the particular conditional likelihood averaging/optimizer settings used to fit models to simulated data.

We analyzed the results of our simulation study through both model selection and parameter inference-based approaches. For model selection, we used sums of small sample size corrected Akaike Information Criteria (AICc) weights across all *X_o_*-dependent models to gauge whether simulated datasets exhibit “significant” support for a relationship between rates of *Y* evolution and the observed factor *X_o_*. For parameter inference, we compared model-averaged predictions of *X_o_*-rate relationships based on AICc weights to the true relationships used to simulate data (see Online Appendix for further details on both our model selection and model-averaging procedures).

### Empirical Example

We applied our pipeline for inferring factor-dependent rates of continuous trait evolution to test whether rates of phenotypic evolution are associated with size among eucalypts, a diverse clade of ∼800 predominantly Australian woody plant species. Body size is often associated with many aspects of life history, as larger organisms tend to live in smaller populations with slower generational turnover (Calder, 2001; Niklas and Enquist, 2001; White et al., 2007; Sibly and Brown, 2007; Adler et al., 2014; Salguero-Gómez et al., 2016; Bakewell et al., 2020). This pattern has led to the hypothesis that evolutionary rates should generally be slower for larger organisms compared to their smaller relatives. However, how rates of phenotypic evolution scale with body size remains unclear, particularly in plants (Cooper and Purvis, 2009; Baker et al., 2015; Chira et al., 2018; Friedman et al., 2019; Zimova et al., 2023; see also Lanfear et al., 2013; Boucher et al., 2017; Vasconcelos et al., 2022). Eucalypts are a powerful system for investigating relationships between body size and rates of phenotypic evolution in plants, as their growth forms range from shrubby “mallees” only reaching a few meters in height (e.g., *Corymbia setosa*, *Eucalyptus erectifolia*, *E. macrocarpa*, *E. odontocarpa*, *E. vernicosa*) to gargantuan trees that grow nearly 100 meters tall (e.g., *E. obliqua*, *E. regnans*, *E. viminalis*; Brooker et al., 2015; Nicolle, 2024).

Leveraging a recently-published dataset consisting of 101 nuclear exonic loci (Crisp et al., 2024), we inferred a time-calibrated phylogeny consisting of 388 ingroup tips belonging to 318 unique eucalypt species representing most major subclades of the group (see Online Appendix for details on phylogenetic inference). We then gathered species-level data on maximum reported measurements for 14 continuous traits: height (a proxy for body size which acts as a factor in our analyses), 8 flower traits (peduncle length, inflorescence/infructescence pedicel length, bud length/width, fruit length/width, seed length), and 5 leaf traits (adult petiole length, juvenile/adult leaf length/width). We obtained these data using a regular expression-based approach to extract continuous trait measurements from all species descriptions in the online Australian eucalypt encyclopedia EUCLID (Brooker et al., 2015). Species descriptions yielding missing and/or outlier measurements were flagged and manually checked/corrected as necessary (see Online Appendix for details on trait data processing). We supplemented the EUCLID data with trait measurements for *E. deglupta*—the only species in our phylogeny not native to Australia—from several regional floras (Pool, 2001; Parnell and Chantaranotahi, 2002; Barrie, 2009) and obtained trait data from published descriptions for two recently named eucalypt species in our phylogeny, *E. connexa* and *E. plumula* (Nicolle and French, 2021). Our final dataset consisted of 311 eucalypt species with 0% missing data for height, 0–3.5% for floral traits, and 5.5–9% for leaf traits. All trait values were log-transformed for our comparative analyses. Prior to applying our pipeline, we used Pagel’s lambda test via the R package *geiger* (Pennell et al., 2014) to assess the level of phylogenetic signal for each trait.

We generated 400 continuous stochastic character maps of eucalypt height (with a resolution of 100) under an evolving rates or “evorates” model whereby rates of height evolution are allowed to gradually change over time and across lineages (Martin et al., 2023; see appendix for further details). We then generated 400 corresponding dummy factor maps by simulating trait evolution with root values and rates identical to those of the height maps. Using these continuous stochastic character maps, we fit seven Brownian Motion models to each individual trait—six models assuming rates depend either on height or the dummy factor through a simple, threshold, or sweetspot relationship, plus a constant rates model (see Online Appendix for details on optimizer settings). Despite using maximum reported measurements for our trait data, we still inferred measurement error parameters in all models for two reasons. First, most measurements were not exact but rounded to the nearest mm or cm (or even 5 m increments in the case of height measurements for tall eucalypt species). Second, incorporating measurement error into our models should render rate variation inferences more conservative in the face of general “tip fog” (e.g., assigning trait data to the wrong tip due to taxonomic conflicts; Beaulieu and O’Meara, 2025) and/or deviations from Brownian Motion-like evolution (Landis and Schraiber, 2017).

To explore the effect of uncertain free parameter estimation on our empirical inferences, we used the R package *dentist* (Boyko and O’Meara, 2024) to sample free parameter values around our maximum likelihood estimates. For each height-dependent and constant rate model, we ran Markov chain Monte Carlo under dentist’s default settings for 10,000 iterations. We used the resulting samples to construct 95% confidence intervals on predicted size-rate relationships under each height-dependent model by taking the minimum/maximum predicted rates for each height value across all sampled parameter values within their respective 95% confidence regions. We then used AICc weights to model-average the 95% confidence interval bounds for each trait.

Finally, we explored the distribution of marginal rate estimates across the phylogeny under our fitted models (“rate mapping”). To do so, we first calculated the estimated rates at each time point of the phylogeny for each height/dummy factor map under the maximum likelihood estimates for a given model. We then used the conditional likelihoods associated with each continuous stochastic character map under the model to take a weighted average of the estimated rates at each time point, yielding a single overall rate map for a given model. Lastly, we used AICc weights to model-average the resulting rate maps across all models for a given trait and generate overall rate maps for each trait. In addition to rates maps for individual traits, we also summarized patterns of rate variation across multiple traits by rescaling the rate maps to have a mean rate of 1—thereby accounting for variation in average evolutionary rates across different traits—and averaging these “relative rates” at each time point. To quantitatively describe the size-rate associations implied by these rate maps, we used the *phylolm* R package (Ho and Ané, 2014) to estimate the relationship between mapped rates and maximum heights at the tips of the phylogeny.

## Results

### Validation Study

Overall, our validation study confirms our continuous stochastic character mapping algorithm correctly samples phenotypic values across a phylogeny according to their expected distribution under a given Brownian Motion model. Kolmgorov-Smirnov (KS) tests comparing observed and expected distributions of Mahalanobis distances for all simulations yielded more-or-less uniformly distributed p-values with only ∼5.4% of p-values falling below 0.05, as expected under the null hypothesis (i.e., the observed samples follow their expected distributions; Fig. 4). This result held across different simulation condtions, with the observed proportions of p-values less than 0.05 not significantly differing from 5% across all conditions (*p* ≈ 0.3 according to a Monte Carlo *χ*^2^ test).

**Figure 4.**
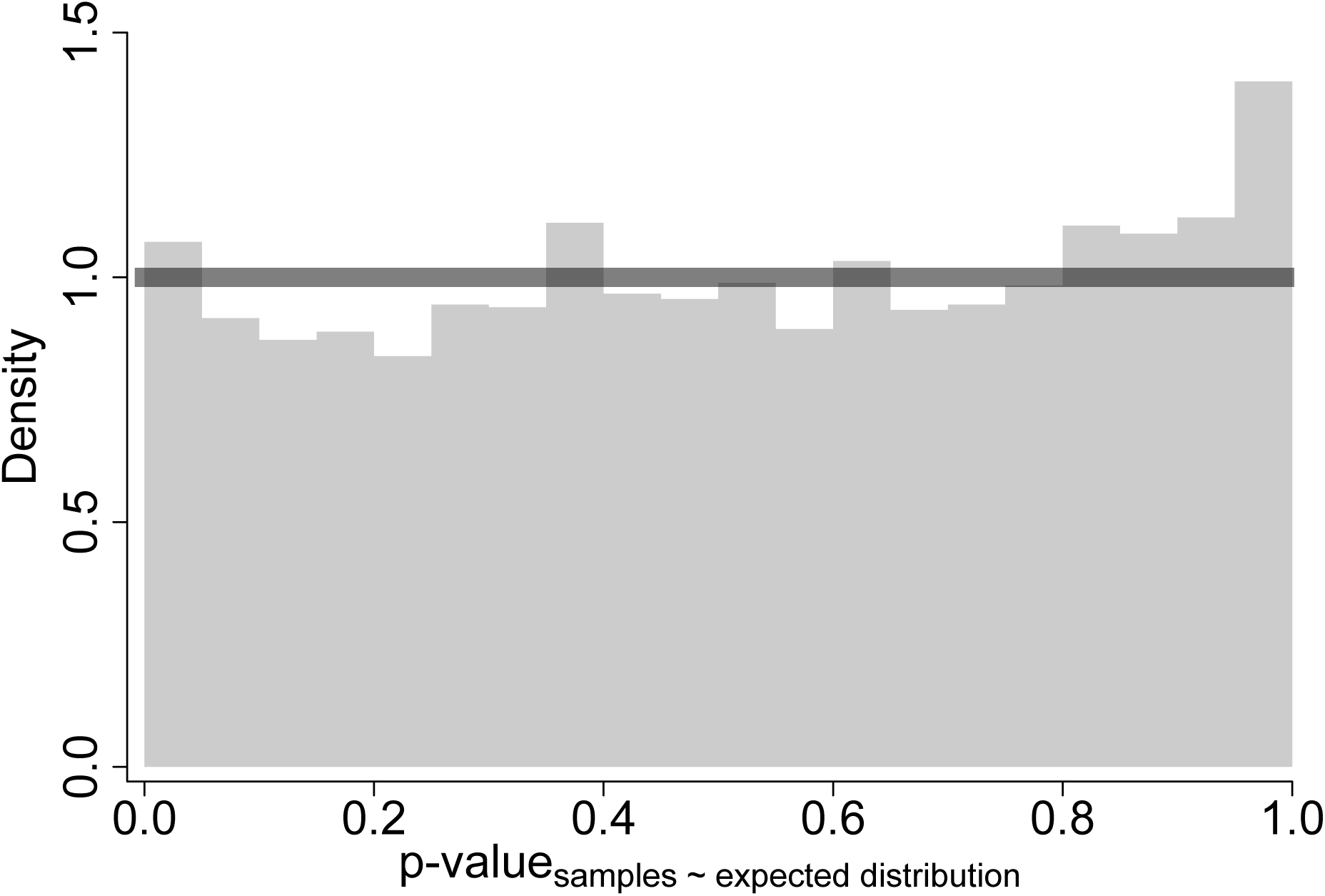
Distribution of p-values for Kolmgorov-Smirnov (KS) tests comparing samples generated by our continuous stochastic character mapping algorithm to their expected multivariate normal distribution under a given Brownian Motion model (using observed Mahalanobis distances and comparing to *χ*^2^ distributions). Assuming the null hypothesis is true—that is, observed samples do follow their expected distributions—this distribution should be uniform. The thick horizontal line indicates the expected density of a uniform distribution for reference.

Closer examination of variation in KS test statistics/p-values based on particular features of different simulation conditions (e.g., number of phenotypes, amount of missing data, etc.) further confirmed a lack of significant differences, with the notable exception that p-values tended to be higher when the same parameters were used to both simulate data and generate continuous stochastic character maps/compute expected distributions. Indeed, an additional KS test suggests the overall distribution of p-values under these conditions deviated from exact uniformity due to a slightly higher-than-expected frequency of p-values around 1 (*p <* 5 × 10*^−^*^7^; also apparent in overall distribution of p-values in Fig. 4). In contrast, the distribution of p-values for conditions under which maps/expected distributions were based on randomly sampled parameters appeared exactly uniform (*p* ≈ 0.75). Ultimately, we suspect using identical parameters to simulate data and compute expected distributions resembles scenarios whereby expected distributions are derived from observed data, which violates a key assumption of the one-sample KS test and results in high, overly conservative p-values (Capasso et al., 2008). We therefore conclude that our continuous stochastic character mapping algorithm works correctly in general and does not noticeably break down under any of the conditions tested here.

### Pipeline Simulation Study

To illustrate potential uses of continuous stochastic character maps, we performed a simulation study to investigate whether continuous stochastic character maps can be used to infer relationships between rates of continuous trait evolution and continuously-varying factors, in analogy with popular approaches for inferring relationships between rates and discretely-varying factors (e.g., Revell, 2013a). Generally, we found model selection using our new pipeline most effective when using AICc weights to summarize the relative goodness-of-fit over multiple models, rather than directly comparing AICc among individual models (see Online Appendix for further details on our model selection procedure). Here, we considered a simulated dataset to exhibit “significant” overall evidence for a factor-rate relationship if the sum of AICc weights for all factor-dependent models (excluding those based on simulated dummy factors) exceeded a cutoff. Averaging across all simulation conditions, a cutoff of 0.9 yields error (i.e., false positive) and power (i.e., true positive) rates of ∼5 and 65%, respectively (Tables 1-2; Fig. 5), while relaxing this cutoff to 0.8–0.7 increases these rates to about 7–11% and 71–76%, respectively (Tables S3-S6).

**Figure 5.**
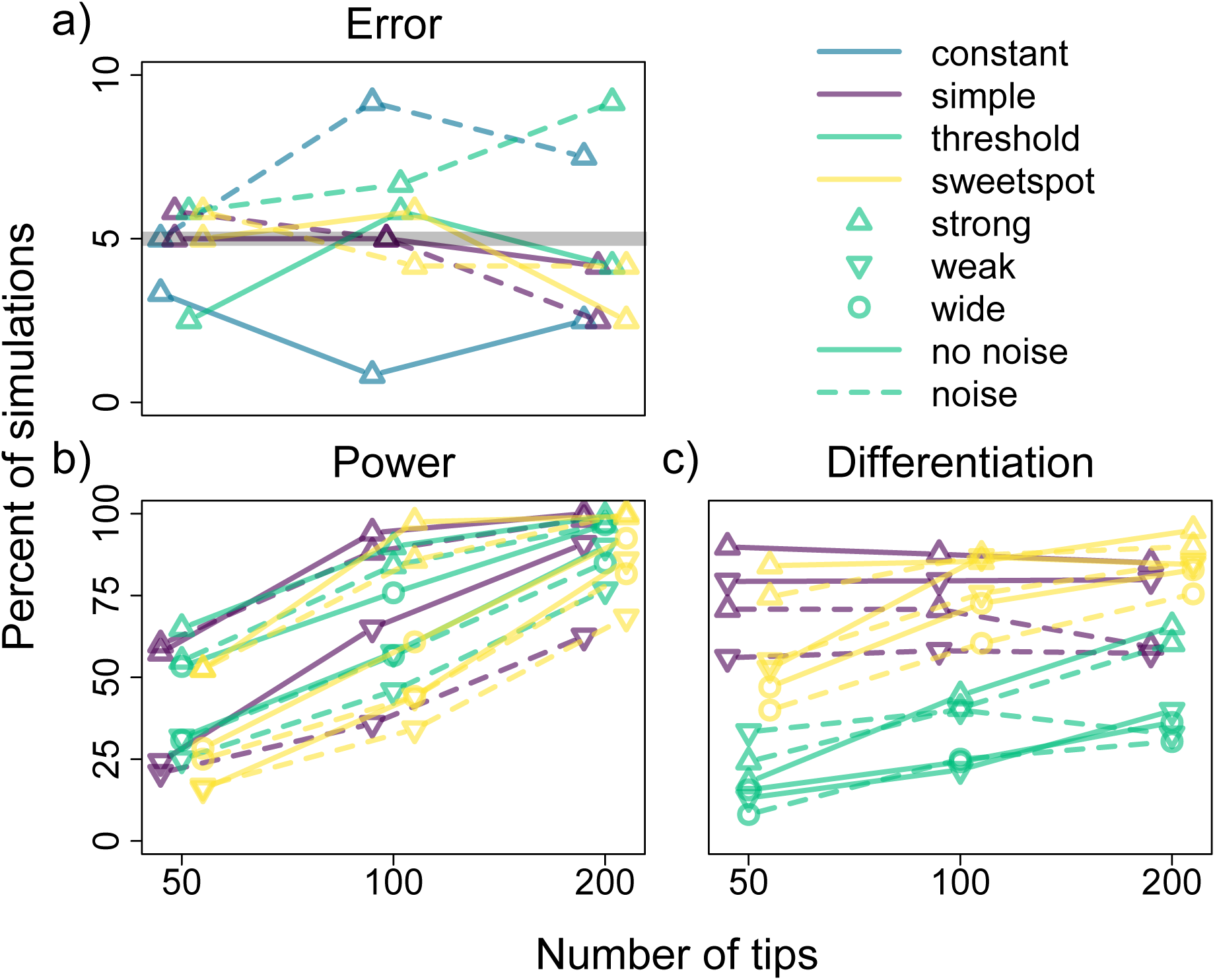
Error, power, and differentiation rates of our continuous stochastic character map-based pipeline for inferring relationships between rates of trait evolution and continuously-varying factors based on a summed AICc weight cutoff of 0.9 (see Figs. S1-S2 for results with lower cutoffs of 0.8 and 0.7). Different colors correspond to simulations with different factor-rate relationships (i.e., constant, exponential/simple, logistic/threshold, and Gaussian/sweetspot), different symbols to simulations with differing relationship strength and—in the case of threshold and sweetspot models—width, and dashed versus solid lines to simulations with versus without random variation in rates (noise) around simulated factor-rate relationships. Panel a: percent of simulations (120 replicates per condition) with either constant or hidden factor-dependent rates for which the best-fitting model is an observed factor-dependent model (i.e., error rates), with a thick gray line indicating a rate of 5%. Panel b: percent of simulations (120 replicates per condition) with observed factor-dependent rates for which the best-fitting model is also an observed factor-dependent model (i.e., power rates). Panel c: percent of observed-factor dependent simulations (varying number of replicates per condtion) exhibiting significant evidence for a factor-rate relationship for which the best fitting model assumed the same kind of factor-rate relationship used to simulate the data (i.e., differentiation rates).

**Table 1.** Proportions of times (based on 120 replicates) our continuous stochastic character map-based method for inferring relationships between rates of trait evolution and continuously-varying factors selected a model assuming a given kind of factor-rate relationship (i.e., null, exponential/simple, logistic/threshold, or Gaussian/sweetspot; corresponding to different rows) as the best-fitting one (based on a summed AICc weight cutoff of 0.9; see Tables S3-S6 for results with lower cutoffs of 0.8 and 0.7). Proportions for simulations with and without random variation around simulated factor-rate relationships (noise) are given within and without parentheses, respectively. Here, we report results for simulations with constant or hidden factor-dependent rates only and consider all models assuming either constant or dummy factor-dependent rates to be null models (see Table S2 for results with each null model separated out).

|  | constant | hidden factor-dependent |  |  |
| --- | --- | --- | --- | --- |
|  |  | simple | threshold | sweetspot |
|  | 50 tips |  |  |  |
| null | 0.97 (0.95) | 0.95 (0.94) | 0.98 (0.94) | 0.95 (0.94) |
| simple | 0.03 (0.00) | 0.03 (0.02) | 0.00 (0.02) | 0.02 (0.01) |
| threshold | 0.01 (0.03) | 0.00 (0.02) | 0.01 (0.03) | 0.02 (0.02) |
| sweetspot | 0.00 (0.02) | 0.02 (0.03) | 0.02 (0.02) | 0.02 (0.03) |
|  | 100 tips |  |  |  |
| null | 0.99 (0.91) | 0.95 (0.95) | 0.94 (0.93) | 0.94 (0.96) |
| simple | 0.00 (0.01) | 0.03 (0.03) | 0.00 (0.00) | 0.02 (0.00) |
| threshold | 0.01 (0.03) | 0.01 (0.00) | 0.03 (0.03) | 0.01 (0.02) |
| sweetspot | 0.00 (0.06) | 0.01 (0.03) | 0.03 (0.04) | 0.03 (0.03) |
|  | 200 tips |  |  |  |
| null | 0.98 (0.92) | 0.96 (0.98) | 0.96 (0.91) | 0.98 (0.96) |
| simple | 0.00 (0.00) | 0.00 (0.00) | 0.00 (0.00) | 0.00 (0.00) |
| threshold | 0.01 (0.01) | 0.02 (0.01) | 0.01 (0.02) | 0.00 (0.02) |
| sweetspot | 0.02 (0.07) | 0.03 (0.02) | 0.03 (0.07) | 0.03 (0.03) |

**Table 2.**
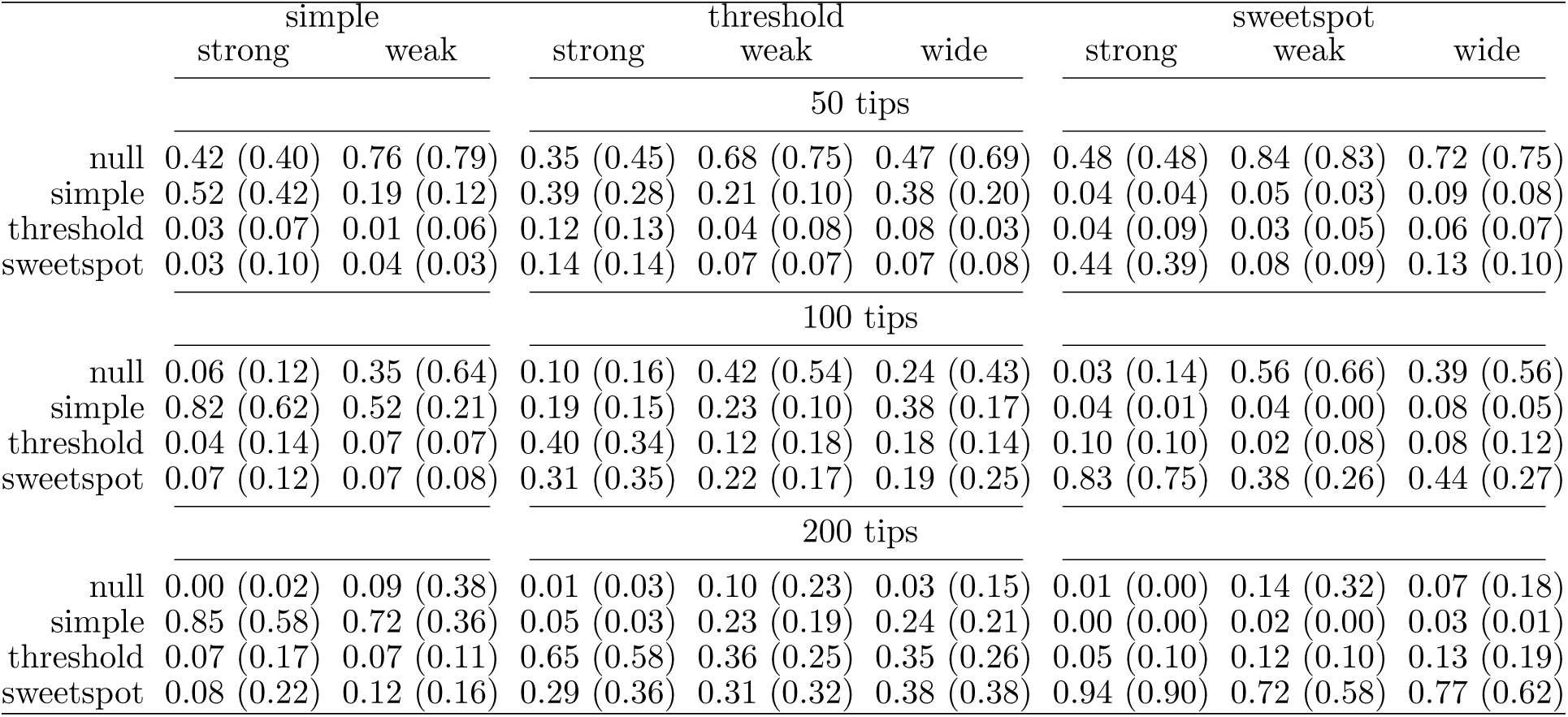
Proportions of times (based on 120 replicates) our continuous stochastic character map-based method for inferring relationships between rates of trait evolution and continuously-varying factors selected a model assuming a given kind of factor-rate relationship (i.e., null, exponential/simple, logistic/threshold, or Gaussian/sweetspot; corresponding to different rows) as the best-fitting one (based on a summed AICc weight cutoff of 0.9; see Tables S3-S6 for results with lower cutoffs of 0.8 and 0.7). Proportions for simulations with and without random variation around factor-rate relationships (noise) are given within and without parentheses, respectively. Here, we report results for simulations with factor-dependent rates only and consider all models assuming either constant or dummy factor-dependent rates to be null models (see Table S2 for results with each null model separated out).

Our pipeline exhibited consistent error rates across most simulation conditions (Fig. 5a), though error tended to decline as sample size increased (i.e., greater numbers of tips), especially when using AICc weight cutoffs lower than 0.9 (Figs. S1-S2a). Additionally, error rates were elevated under conditions with noisy yet otherwise relatively constant evolutionary rates (e.g., constant and threshold factor-rate relationships)—presumably reflecting dummy factor-dependent models being more effective at modeling rate variation due to hidden factors than noise. Perhaps unsurprisingly, simulation conditions with perfectly constant rates exhibited relatively low error rates.

Power (Fig. 5b) varied drastically across simulation conditions, increasing with both sample size and the magnitude of factor-associated rate variation. Among simulations with strong, narrow factor-rate relationships (i.e., ∼20-fold rate variation over factor values between −2 and 2), power rates ranged from just above 50% for 50-tip simulations to over 90% for 200-tip simulations based on an AICc weight cutoff of 0.9. In contrast, under conditions with wider (i.e., ∼20-fold variation from −4 to 4) or weaker (i.e., ∼5-fold variation from −2 to 2) relationships, power dropped to around 25–50% for 50–100-tip simulations and 60–90% for 200-tip simulations. Relaxing the AICc weight cutoff to lower values increased power rather evenly across all simulation conditions (Figs. S1-S2b). Noise around simulated factor-rate relationships tended to reduce power, though this trend was generally weaker at both small *and* large sample sizes. This pattern likely reflects our pipeline’s limited ability to detect rate noise at smaller sample sizes, while signals of factor-rate associations were strong enough to “overcome” any effects of noise at sufficiently large sample sizes. Comparing among different types of simulated factor-rate relationships, our pipeline typically exhibited the most power to detect factor-rate associations under simple (i.e., exponential) relationships, followed by threshold (i.e., logistic) and sweetspot (i.e., Gaussian) relationships. However, noisy and weak simple relationships were associated with exceptionally low power at large sample sizes. Additionally, the power to detect factor-rate associations under sweetspot relationships grew rapidly with sample size.

We also investigated our pipeline’s ability to distinguish among different types of factor-rate relationships by comparing AICc among factor-dependent models (i.e., “differentiation rates”; Fig. 5c; note that this metric only considered simulated datasets already exhibiting significant overall support for a factor-rate relationship based on summed AICc weights). While our approach was effective at correctly inferring simple and sweetspot relationships, it struggled to detect threshold relationships. Instead, threshold relationships were commonly mistaken for simple relationships at small sample sizes and for sweetspot relationships at large sample sizes. Our pipeline also exhibited a slight bias towards inferring more complex factor-rate relationships as sample size increased; while the ability to correctly infer threshold and sweetspot relationships improved with sample size, differentiation rates for simple relationships declined slightly. Noise also caused simple relationships to be mistaken for threshold/sweetspot relationships more frequently. Notably, while power rates were typically lowest for simulations with weak factor-rate relationships, differentiation rates were lowest for those with wide relationships. Thus, while our pipeline’s power was mainly affected by the overall magnitude of factor-associated rate variation, differentiation appeared more strongly influenced by how abruptly rates change with respect to factor values.

Beyond AICc-based model selection, we explored how accurately our pipeline can infer the particular shape of factor-rate relationships based on model-averaged predictions (Fig. 6). Overall, predicted factor-rate relationships largely recapitulated simulated ones—particularly as sample size increased. However, predicted rates also exhibited shrinkage whereby low and high rates were over- and underestimated, respectively (Fig. S3). This shrinkage tended to flatten predicted factor-rate relationships, especially under conditions with limited power (e.g., small sample sizes, weak/wide relationships). However, even predictions for 200-tip simulations could exhibit minor shrinkage around factor values associated with abrupt rate shifts as well as extreme factor values in the tails of their simulated distribution (i.e., generally factor values less than −2 or greater than 2 under our simulation conditions). In contrast to predicted rates for extreme factor values, predictions at “typical” factor values towards the center of their distribution (i.e., factors values between −2 and 2) appeared fairly accurate if imprecise (Figs. S4-S5). While predicted rates at typical factor values often varied widely across simulations with identical factor-rate relationships (∼3–30-fold in general), they only differed from simulated rates by a median 15–50%. Noise around simulated factor-rate relationships decreased the accuracy and precision of predicted rates, though this effect was small relative to that of sample size.

**Figure 6.**
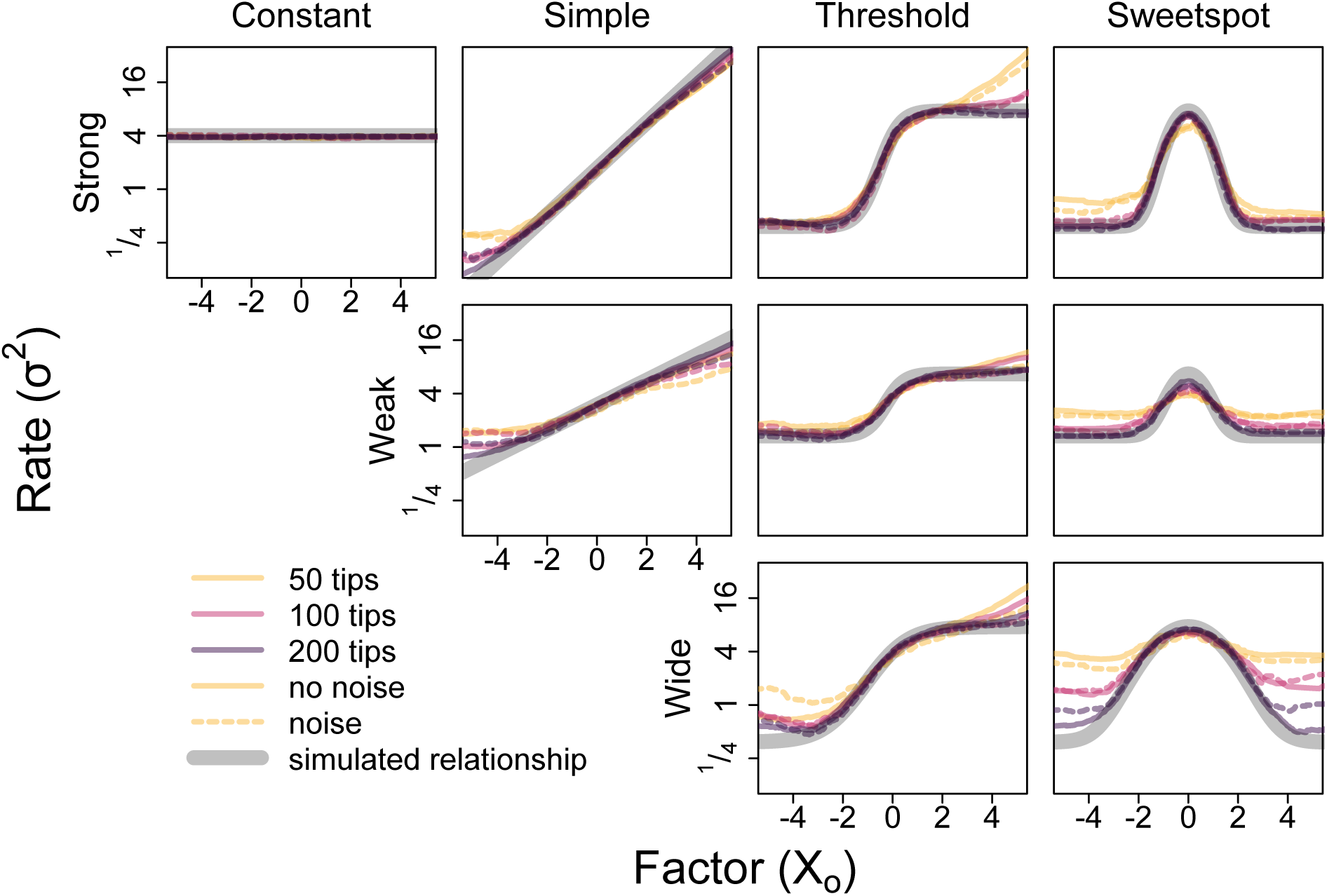
Median model-averaged factor-rate relationships inferred using our continuous stochastic character map-based pipeline for inferring relationships between rates of trait evolution and continuously-varying factors under all simulation conditions with either constant or observed factor-dependent rates. Different colors correspond to different phylogeny sizes (i.e., number of tips) and dashed versus solid lines to simulation conditions with versus without random variation in rates (noise) around simulated factor-rate relationships. Simulated factor-rate relationships are represented by thick gray lines for reference.

### Empirical Example

In addition to the simulation study, we applied our continuous stochastic character map-based pipeline for inferring factor-dependent rates of continuous trait evolution to test whether rates of flower and leaf trait evolution scale with body size among eucalypts.

All traits investigated here exhibited variable yet consistently significant phylogenetic signal, with weak to moderate signal among leaf traits (*λ* ∼0.3–0.7), moderate to strong signal among flower traits (∼0.6–0.95), and relatively strong signal for height, our proxy for body size (∼0.85). Across all traits, dummy factor-dependent models tended to account for the majority of AICc weight, implying that rates of flower and leaf trait evolution, while variable, are mostly unrelated to size. Only bud width/length (summed AICc weight *>* 0.85) and adult leaf width (summed AICc weight *>* 0.7) exhibit notable evidence for associations between size and evolutionary rates (Fig. 7). Bud length and width both show substantial support for sweetspot size-rate relationships in particular, though the specific shape of this relationship differs between the two traits. Model-averaged predictions suggest a ∼3-fold peak in rates of bud length evolution among lineages around 5 m tall yet a 10-fold dip in rates of bud width evolution among lineages around 15 m tall (Fig. 8). Nonetheless, these inferences both suggest that rates of bud size evolution decrease with size between roughly 5 and 15 m in height. In contrast to results for bud size, rates of leaf width evolution exhibit a largely simple size-rate relationship, with rates decreasing by ∼5% per 10% increase in height.

**Figure 7.**
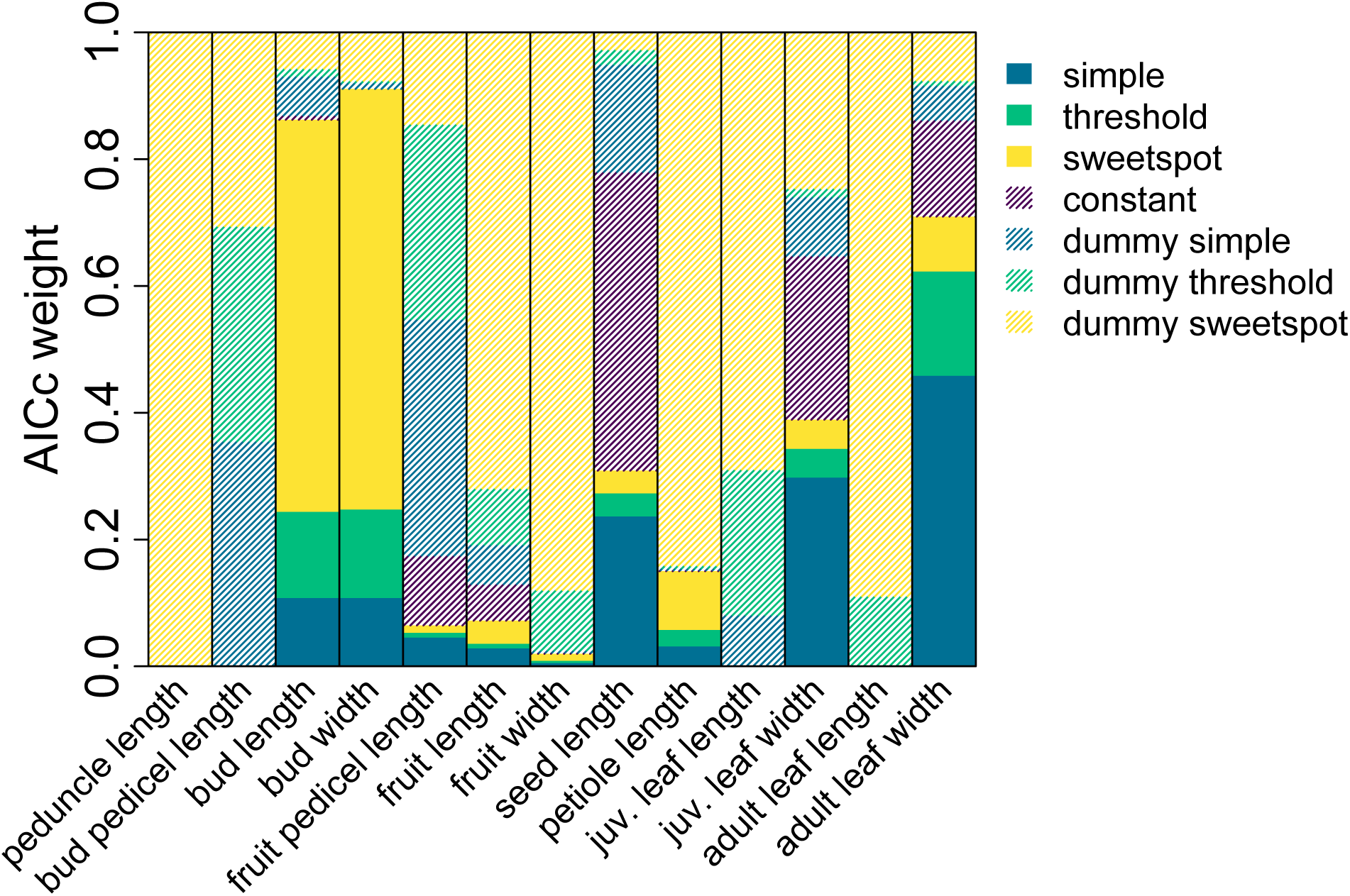
AICc weights of models assuming different factor-rate relationships for each eucalypt trait using our continuous stochastic character map-based pipeline (juv. = juvenile). Colors correspond to different kinds of factor-rate relationships (i.e., constant, exponential/simple, logistic/threshold, and Gaussian/sweetspot). Solid colors correspond to non-null models assuming rates depend on height while line-shaded colors represent null models assuming rates are either constant or depend on simulated dummy factors.

**Figure 8.**
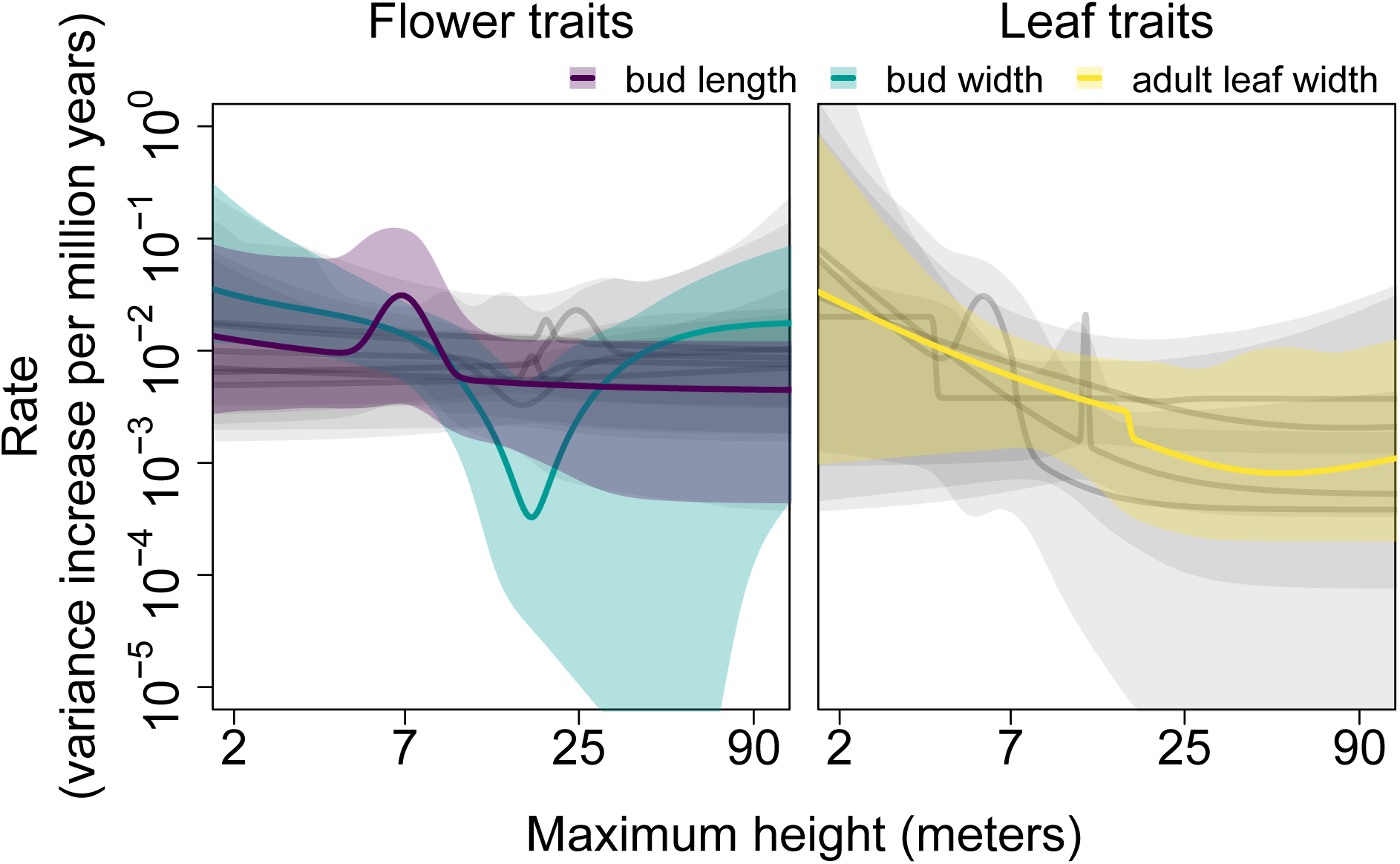
Model-averaged height-rate relationships for all eucalypt traits inferred using our continuous stochastic character map-based pipeline. Lines depict relationships based on maximum likelihood estimates while the sorrounding shaded regions represent 95% confidence intervals calculated using *dentist* (Boyko and O’Meara, 2024). Lines and shaded regions for traits exhibiting substantial evidence of factor-rate associations (i.e., summed AICc weight *>* 0.7) are drawn in color, while those for other traits are drawn in light gray. Note that all traits were log-transformed, such that rates are measure increases in *geometric* variance per million years.

While evidence for size-rate relationships was weak when considering traits individually, we consistently inferred a negative association between size and rates under simple models for all traits except inflorescence pedicel and seed length (Tables S7-S19), suggesting that rates of flower and leaf trait evolution are higher among smaller-bodied eucalypt lineages overall. We further investigated this trend by calculating fold-differences between average rates predicted for eucalypts greater and less than 15 m tall (close to the median height across all species) based on model-averaged relationships for each trait (Fig. 9). Averaging across all traits, rates of phenotypic evolution are ∼40% lower among eucalypts that grow taller than 15 m compared to those less than 15 m tall, though the confidence intervals for individual traits overlapped with 1 (i.e., no change) in all cases except bud length/width and petiole length.

**Figure 9.**
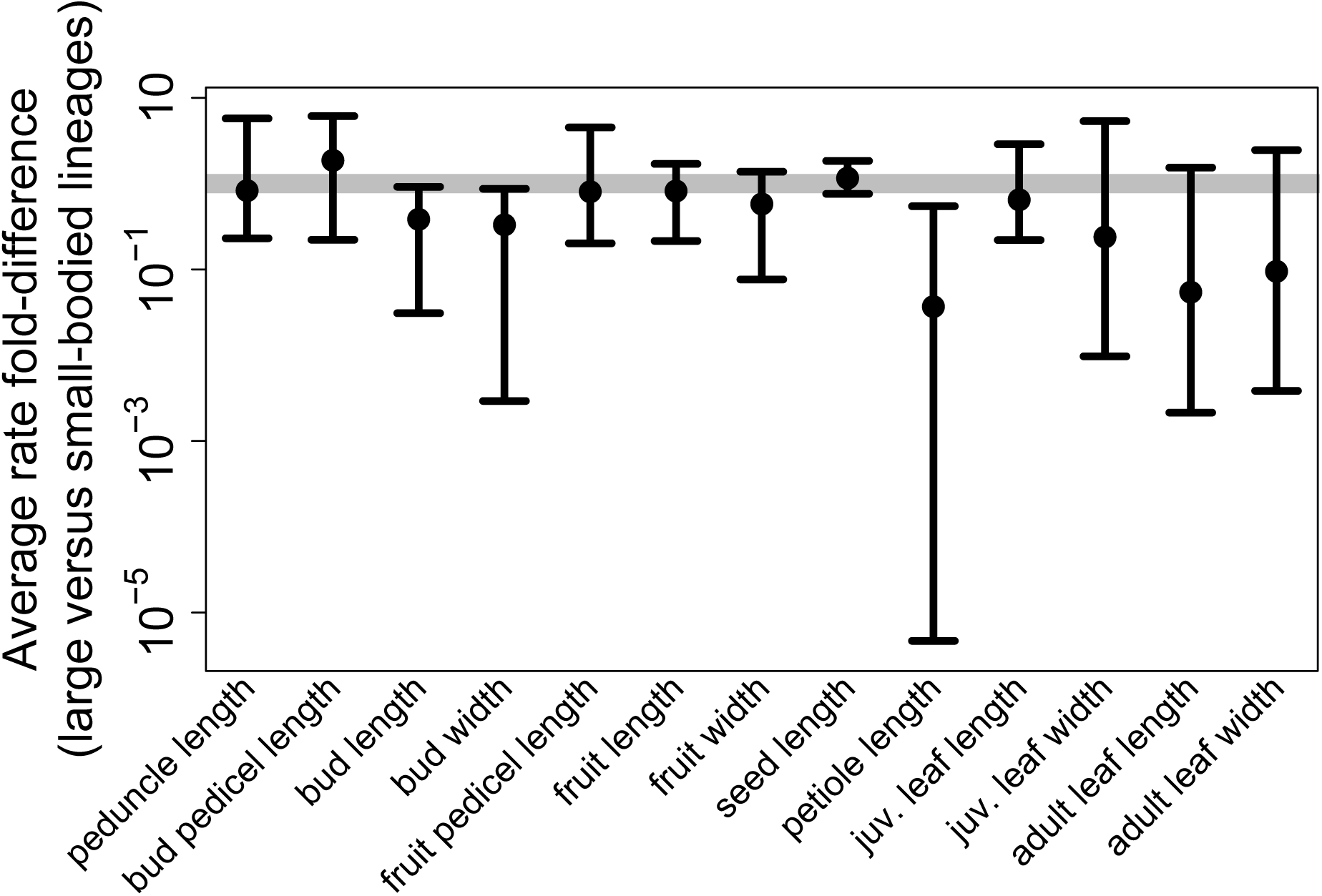
Model-averaged fold-differences in average rates among eucalypts greater and less than 15 meters tall for all traits inferred using our continuous stochastic character map-based pipeline. Points correspond to maximum likelihood estimates while error bars represent 95% confidence intervals on these differences calculated using *dentist* (Boyko and O’Meara, 2024). The thick gray line corresponds to a fold-difference of 1 (i.e., no change) for reference.

Despite substantial size-independent rate variation inferred by dummy factor-dependent models, mapping estimated rates onto the eucalypt phylogeny still revealed a negative association between size and rates of flower and leaf trait evolution overall, especially among smaller eucalypts growing ∼20 m tall or less (Fig. 10; see Figs. S6-S18 for individual traits and Figs. S19-S20 for separate summaries of flower and leaf traits). A phylogenetic generalized least squares regression including both a linear and quadratic term for height best explained variation in relative tip rates averaged across all traits, suggesting overall rates of phenotypic evolution dip to a minimum around 25 m and increase by about 200% and 50% among lineages around 3 and 75 m tall, respectively. Height explained ∼15% of variation in tip rates according to this model, with much of the residual variation stemming from a relatively small number of outliers characterized by rates some 2 to 10 times higher than expected.

**Figure 10.**
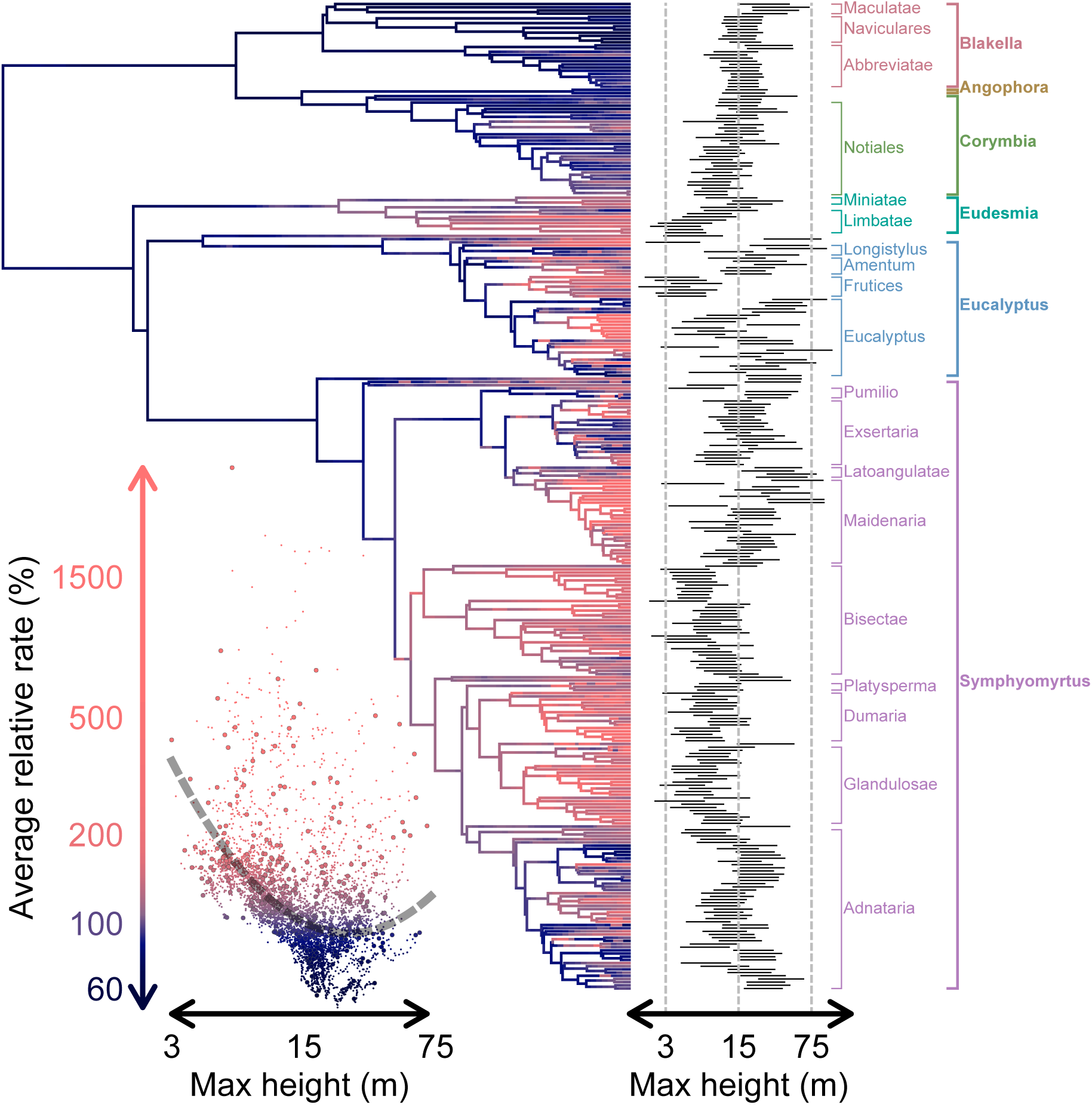
Relative rates of phenotypic evolution, averaged over all flower and leaf traits, mapped onto the eucalypt phylogeny, with dark blue and light red colors indicating relatively low and high rates, respectively (see y-axis of plot on bottom left for color scale; note that the color gradient is percentile-based due to the right-skewed distribution of relative rates on a log scale). Bars depicting 95% credible intervals on maximum height for each tip (inferred under an “evolving rates” or evorates model of height evolution) are arrayed along the right side of the phylogeny, along with clade labels indicating major eucalypt sections and subgenera/genera. The bottom left plot consists of mapped rates for each time point with respect to mean maximum heights (as inferred under the evorates model of height evolution), with larger points indicating tip heights/rates. The gray dashed line running through the plot represents a phylogenetic generalized least squares regression relating tip rates to maximum heights.

## Discussion

In this paper we introduce continuous stochastic character mapping, a flexible and efficient tool for sampling evolutionary histories of continuous variables along phylogenies based on Brownian Motion models of trait evolution. Like conventional stochastic character mapping for discrete variables, this technique can be used to generate distributions of probable evolutionary histories for continuous variables. Beyond their use in exploring continuous trait evolutionary dynamics while accounting for uncertainty in reconstructed ancestral states, these distributions enable the development of statistical pipelines for investigating how evolutionary patterns across phylogenies are associated with continuous variables. Such approaches will help researchers address long-standing questions regarding the impacts of continuously-varying factors like body size (e.g., Friedman et al., 2019), generation time (e.g., Gingerich, 2001), and/or climatic niche (e.g., Tribble et al., 2023) on macroevolutionary dynamics. To illustrate these applications, we implemented a pipeline capable of detecting and quantifying relationships between continuously-varying factors and rates of continuous trait evolution. We applied this novel approach to an empirical example, testing whether body size is related to rates of flower and leaf trait evolution in eucalypts. While our analyses revealed substantial size-independent variation in rates of phenotypic evolution across different traits and clades, we found that smaller-bodied eucalypt lineages exhibit elevated rates overall. Ultimately, our results demonstrate how continuous stochastic character maps provide a powerful toolkit for analyzing comparative data on continuous variables.

### Continuous stochastic character maps are useful for inferring factor-rate associations

Our simulation study revealed that continuous stochastic character mapping provides an effective and pragmatic approach for inferring relationships between rates of trait evolution and continuously-varying factors. Previous approaches for estimating these relationships were developed for particular empirical studies to circumvent the lack of a standard method (e.g., Cooper and Purvis, 2009; Uyeda et al., 2021). Accordingly, the general statistical performance of these methods remains largely unknown and their utility is limited by a lack of generalizable code for implementation. The *evorag* method developed by Weir and Lawson (2015) constitutes a notable exception as a well-studied method suitable for a variety of purposes, though it only uses sister pair contrasts to infer factor-rate relationships. By using entire phylogenies, our pipeline more fully utilizes the information available in phylogenetic comparative data to infer factor-rate relationships. More recently, Hansen et al. (2022) developed a regression-based approach for inferring correlations between rates and continuous variables, though this method is only capable of inferring linear or exponential relationships between a single continuous factor and rates for a single continuous trait (or intraspecific variances, which are interpreted as “microevolutionary rates” by the authors). By comparison, our method offers more flexibility, allowing researchers to incorporate explicit reconstructions of both discrete and multiple continuous factors into analyses, fit models to multivariate continuous trait data, and/or simultaneously link multiple parameters (i.e., rates, intraspecific variances, and/or evolutionary trends) to reconstructed factors via arbitrary parameter functions (computational/statistical tractability notwithstanding). Overall, our new method is a substantial generalization of previous ones, expanding the hypothesis-testing toolkit available for macroevolutionary analyses.

While our pipeline was generally accurate and robust, it was limited under certain conditions. For one, the approach struggled to detect relatively weak factor-rate relationships from smaller phylogenetic comparative datasets. Based on the simulation conditions examined here, our pipeline typically requires phylogenies with 100 tips to consistently detect ∼20-fold differences in rates associated with a continuous factor, and 200 tips for 5-fold differences. Nonetheless, our pipeline was able to reliably reject factor-rate associations under most simulation conditions—even when rates varied due to unobserved factors. Thus, while our approach may fail to detect weak factor-rate associations, it is generally unlikely to infer spurious associations (though, as with any correlative statistical method, associations may reflect unobserved factors confounded with observed factors/rates). Our pipeline also often failed to distinguish threshold relationships from simple or sweetspot relationships—even using large phylogenies with 200 tips. Interestingly, a previously-developed method for inferring relationships between continuously-varying factors and lineage diversification rates exhibited similar difficulties with distinguishing threshold relationships from linear ones (FitzJohn, 2010). Overall, the low power/differentiation rates of our pipeline under these conditions may reflect inherent limits to phylogenetic comparative inference of factor-rate relationships rather than shortcomings particular to our approach. Recent work suggests detecting associations between evolutionary rates and continuously-varying factors from phylogenetic comparative data may be fundamentally challenging (Hansen et al., 2022).

We also observed shrinkage of rates estimated under our pipeline in some scenarios, whereby high and low rates tend to be under- and overestimated, respectively. Shrinkage has also been found in previous work investigating methods for inferring variation in rates of trait evolution (Revell, 2013a; Martin et al., 2023). Here, shrinkage likely reflects a phenomenon first pointed out by Revell (2013a), who noted that uncertainty in factor histories may cause factor values associated with low rates to be inferred in lineages evolving at relatively high rates and vice versa. A joint inference approach, whereby the factor history and its effect on trait evolution processes are inferred simultaneously rather than sequentially, could help mitigate these issues and enable more accurate inference of factor-rate associations in the future. However, such methods entail significant mathematical, statistical, and computational challenges due to their complexity (e.g., FitzJohn, 2010, May and Moore, 2020, Boyko et al., 2023b).

Our work adds to a growing body of literature demonstrating that accounting for “background” or residual heterogeneity in evolutionary processes is often critical for robust phylogenetic comparative inference and hypothesis testing (Maddison and FitzJohn, 2015; Beaulieu and O’Meara, 2016; Uyeda et al., 2018; May and Moore, 2020; Boyko and Beaulieu, 2023; Boyko et al., 2023b; Tribble et al., 2023). Models based on simulated dummy factors were often crucial for controlling our method’s error rates by providing more competitive null models which were able to account for rate variation due to unobserved factors. However, elevated error rates under a few simulation conditions (e.g., noisy constant rates) suggest dummy factor-dependent models were less effective at accounting for primarily random rate variation that does not exhibit phylogenetic autocorrelation (i.e., noise, as opposed to the phylogenetically autocorrelated rate variation driven by observed/hidden factors in our simulations). Despite this, dummy factor-dependent models still reduced error rates quite substantially under such conditions—particularly as sample size increased (see Tables S2, S3, and S5).

### Smaller eucalypts exhibit higher rates of phenotypic evolution

Using our continuous stochastic character map-based pipeline for inferring relationships between rates of trait evolution and continuous factors, we show that rates of flower and leaf trait evolution are elevated among lineages of smaller eucalypts overall. As far as we know, this is the first published analysis to investigate the relationship between body size and rates of phenotypic evolution in any plant group. Previous work has been limited to vertebrates and yielded mixed results, suggesting body shape evolution is slower among larger fish and birds (Friedman et al., 2019; Zimova et al., 2023; but see Chira et al., 2018) while body size and cranial shape evolution are generally faster among larger mammals (Cooper and Purvis, 2009; Baker et al., 2015). Nonetheless, our results agree with broad empirical evidence for accelerated molecular evolution and lineage diversification among smaller-bodied lineages (Gillooly et al., 2005; Fontanillas et al., 2007; Bromham, 2011; Wollenberg et al., 2011; Etienne et al., 2012; Lanfear et al., 2013; Weber et al., 2014; Boucher et al., 2017; Igea et al., 2017; Tedesco et al., 2017; Friedman et al., 2019; Zimova et al., 2023). Smaller plants in particular tend to exhibit reduced seed dispersal capacity and greater microhabitat specialization, presumably leading to more frequent evolutionary divergence at finer spatial scales compared to larger plants (Boucher et al., 2017). These factors may play a particularly important role in shaping size-rate relationships across eucalypts, as their seeds are gravity-dispersed and typically travel no further than the height of their parent tree (Booth, 2017). Accordingly, the geographic ranges of most eucalypt species span 1% or less of Australia’s total land area (Hughes et al., 1996). While prior comparative studies suggest both range size (Mathews and Bonser, 2005) and lineage diversification rates (Vasconcelos et al., 2022) are largely unrelated to height variation across eucalypts, such patterns are worth re-investigating given the growing availability of eucalypt trait/occurrence data (e.g., Belbin et al., 2021; Falster et al., 2021) along with increasingly well-resolved phylogenetic resources (e.g., Ferguson et al., 2023; McLay et al., 2023; Crisp et al., 2024).

Despite the overall negative association between size and rates of phenotypic evolution, our results also revealed substantial size-independent rate variation among eucalypts. A fundamental difficulty in inferring factor-rate relationships from phylogenetic comparative data is that evolutionary rates are generally influenced by countless interacting factors, yet macroevolutionary researchers are practically limited to only modeling the effect(s) of one to several factors at a time. Even well-supported factor-rate relationships tend to only account for a small portion of apparent rate variation in any given clade (e.g., Hansen et al., 2022). Using rate maps, however, we can at least gauge which clades generally supported or contradicted the overall negative association between size and rates of phenotypic evolution across eucalypts. Based on rates averaged across all traits, we inferred relatively high rates of phenotypic evolution among several clades predominantly consisting of small eucalypt species (e.g., sections Limbatae, Frutices, Bisectae, Dumaria, and Glandulosae). However, larger eucalypts do not always exhibit low rates by comparison, with particularly high rates occurring among some larger-bodied lineages in the subgenus Eucalyptus and section Maidenaria. Notably, each of these clades exhibit exceptionally high rates of size evolution (Fig. S21) and encompass both the smallest and largest known eucalypt species (this may partially account for the inferred increase in rates among the largest eucalypt species compared to medium-sized ones). Ultimately, relatively high rates of phenotypic evolution seem to occur among eucalypt lineages of all sizes, though smaller-bodied lineages nonetheless stand out as exhibiting high rates rather consistently based on our analyses. Interestingly, while this trend is apparent across both major eucalypt clades, rates of flower and leaf trait evolution tend to be slower and less heterogeneous among *Corymbia* s.l. (consisting of (sub)genera *Angophora*, *Corymbia*, and *Blakella*) compared to *Eucalyptus* (consisting of the major subgenera Eucalyptus, Eudesmia, and Symphyomyrtus).

While our empirical investigation demonstrated the utility of our new pipeline overall, it also illustrates an important caveat regarding our model selection procedure. Broadly, our current approach appears effective at clarifying when factor-dependent rate variation exceeds factor-independent variation and vice versa. Our approach does not, however, provide results in a regression-like manner (i.e., inferring the amount of residual variation around an overall trend relating rates to a factor of interest), limiting its ability to reliably detect subtler factor-rate relationships. For example, while AICc weights provided weak to equivocal evidence for size-associated variation in rates of petiole length and juvenile leaf width evolution (Fig. 7), rate maps for the same traits revealed clear negative relationships with just a few extreme outliers (Figs. S14, S16). While careful investigation of estimated parameters and rate maps is useful for identifying cases where model selection is being overly conservative in this manner, explicitly modeling residual rate variation around inferred factor-rate relationships would likely yield more robust and powerful inferences which are less cumbersome to interpret. Unfortunately, modeling residual variation in rate parameters under continuous trait evolution models is not straightforward (which is why our pipeline uses continuous stochastic character maps in the first place), though we see this as a natural next step for improving inference of factor-rate relationships from macroevolutionary data.

### Conclusion and Future Extensions

Here, we developed a novel method for sampling histories of continuous variables on phylogenies under Brownian Motion models of trait evolution, thereby generalizing conventional discrete stochastic character mapping methods to work with continuous variables. We show how continuous stochastic character maps may be used to infer relationships between continuous trait evolution dynamics (with a focus on rates) and continuously-varying factors. Finally, we demonstrate the utility of this approach in an empirical example, showing that rates of flower and leaf trait evolution are generally higher among smaller eucalypts. Ultimately, continuous stochastic character mapping can empower researchers with new and innovative strategies for both analyzing the evolutionary dynamics of continuous phenotypes and testing hypotheses that require accounting for how continuous variables have changed across the evolutionary histories of clades.

Looking forward, continuous stochastic character mapping could be extended to sample evolutionary histories under other popular models of continuous trait evolution such as Ornstein-Uhlenbeck and Lévy processes, which are generally interpreted as models of adaptive and pulsed evolution, respectively. Another promising future direction would be the development of methods for fitting lineage diversification or discrete trait evolution models with parameters depending on mapped continuous variables. Such methods would allow macroevolutionary biologists to investigate the impact of continuously-varying factors on evolutionary dynamics with greater flexibility. Lastly, our algorithm for generating continuous stochastic character maps could benefit data augmentation schemes for Bayesian inference of complex phylogenetic comparative models involving continuously-varying factors (e.g., Lartillot and Poujol, 2011; Lartillot and Delsuc, 2012; Lartillot et al., 2016; Quintero and Landis, 2020; Quintero, 2025), as it can directly transform samples of uncorrelated normal random variables into phenotypic evolutionary histories under Brownian Motion models.

## Acknowledgements

We would like to thank the members of the corresponding author’s former dissertation committee—Luke Harmon, Gideon Bradburd, William Wetzel, and Jeffrey Conner—for their invaluable advice throughout the development of this work. We are especially grateful for their writing suggestions, which helped improve the clarity of this manuscript. Past and present members of the Leebens-Mack lab at the University of Georgia as well as the Weber and Bradburd labs at the University of Michigan (both formerly at Michigan State University) provided additional advice which was particularly helpful in refining how we communicated the work to broader audiences. We would also like to thank participants and organizers of the 2019 Nantucket phylogeny developeR workshop—in particular Nicholas Bone, Fabio Machado, and Josef Uyeda—for their helpful feedback on the very first implementation of continuous stochastic character mapping. Lastly, Isabel Sanmartín, Stacey Smith, Mike May, and two anonymous reviewers provided incredibly comprehensive and useful comments on an earlier version of this manuscript, leading to key refinements of our statistical methods, expansion of our empirical case study, and much more concise, accessible writing.

## Supplementary Material

Data will be made available from a Dryad Digital Repository upon publication. In the meantime, please reach out B.S.M. to request any of the data and/or scripts used in this work. The Online Appendix can be found under Supplementary Material on bioRxiv, while the current version of the *contsimmap* R package is available at the GitHub repository: https://github.com/bstaggmartin/contsimmap.

