## Supplementary material for "Stochastic Character Mapping of Continuous Traits on Phylogenies": Online Appendix

Table S1. *Table of parameter values for each version of simple, threshold, sweetspot, and null parameter functions used to generate data for the simulation study of our continuous stochastic character map-based pipeline for inferring relationships between rates of trait evolution and continuously-varying factors. In general, we consider the strong versions of each function the default, which are standardized to yield an overall rate around 4 that varies ∼20-fold as factor values range from -2 to 2. Weak functions are largely identical to strong functions but instead only cause rates to vary ∼5-fold over the -2 to 2 interval. On the other hand, wide functions still cause rates to vary ∼20-fold but over a wider range of factor values from -4 to 4 (note that there is no wide version of the simple function). Intercept and slope parameters were chosen to ensure simple functions reach the minimum/maximum values of corresponding threshold/sweetspot functions at factor values of -2.5 and 2.5.*

| | parameter | | | | | |
|---|---|---|---|---|---|---|
| function version | intercept ($\beta_0$) | slope ($\beta_1$) | mid-rate ($\alpha$) | location ($\theta$) | rate deviation ($\delta$) | width ($\omega$) |
| strong | $\ln 4 - \ln\cosh\left(\frac{\ln 20}{2}\right)$ | $\frac{\ln 20}{5}$ | $\ln 4$ | 0 | $\frac{\ln 20}{2}$ | $\ln 4$ |
| weak | $\ln 4 - \ln\cosh\left(\frac{\ln 5}{2}\right)$ | $\frac{\ln 5}{5}$ | $\ln 4$ | 0 | $\frac{\ln 5}{2}$ | $\ln 4$ |
| wide | — | — | $\ln 4$ | 0 | $\frac{\ln 20}{2}$ | $\ln 8$ |

Table S2. *Proportions of times (based on 120 replicates) our continuous stochastic character map-based pipeline for inferring relationships between rates of trait evolution and continuously-varying factors selected a model assuming a given kind of factor-rate relationship (i.e., constant, exponential/simple, logistic/threshold, or Gaussian/sweetspot; corresponding to different rows) as the best-fitting one (based on a summed AICc weight cutoff of 0.9). Here, d. and o. denote models based on simulated dummy factors and observed factors, respectively. Proportions for simulations with and without random variation around factor-rate relationships (noise) are given within and without parentheses, respectively.*

| | constant | hidden factor-dependent | | | simple | | threshold | | | sweetspot | | |
|---|---|---|---|---|---|---|---|---|---|---|---|---|
| | | simple | threshold | sweetspot | strong | weak | strong | weak | wide | strong | weak | wide |
| **50 tips** | | | | | | | | | | | | |
| constant | 0.87 (0.48) | 0.22 (0.12) | 0.22 (0.12) | 0.22 (0.08) | 0.15 (0.06) | 0.51 (0.26) | 0.12 (0.08) | 0.48 (0.21) | 0.32 (0.23) | 0.14 (0.03) | 0.52 (0.28) | 0.42 (0.21) |
| d. simple | 0.02 (0.06) | 0.32 (0.22) | 0.24 (0.16) | 0.23 (0.16) | 0.09 (0.07) | 0.06 (0.12) | 0.09 (0.05) | 0.07 (0.10) | 0.05 (0.07) | 0.05 (0.10) | 0.15 (0.11) | 0.07 (0.10) |
| d. threshold | 0.04 (0.22) | 0.07 (0.26) | 0.15 (0.23) | 0.11 (0.18) | 0.03 (0.15) | 0.07 (0.19) | 0.03 (0.10) | 0.05 (0.21) | 0.06 (0.20) | 0.07 (0.13) | 0.05 (0.20) | 0.07 (0.22) |
| d. sweetspot | 0.04 (0.18) | 0.32 (0.34) | 0.36 (0.42) | 0.38 (0.52) | 0.15 (0.12) | 0.12 (0.22) | 0.10 (0.22) | 0.08 (0.23) | 0.04 (0.19) | 0.22 (0.21) | 0.12 (0.25) | 0.15 (0.22) |
| o. simple | 0.03 (0.00) | 0.03 (0.02) | 0.00 (0.02) | 0.02 (0.01) | 0.52 (0.42) | 0.19 (0.12) | 0.39 (0.28) | 0.21 (0.10) | 0.38 (0.20) | 0.04 (0.04) | 0.05 (0.03) | 0.09 (0.08) |
| o. threshold | 0.01 (0.03) | 0.00 (0.02) | 0.01 (0.03) | 0.02 (0.02) | 0.03 (0.07) | 0.01 (0.06) | 0.12 (0.13) | 0.04 (0.08) | 0.08 (0.03) | 0.04 (0.09) | 0.03 (0.05) | 0.06 (0.07) |
| o. sweetspot | 0.00 (0.02) | 0.02 (0.03) | 0.02 (0.02) | 0.02 (0.03) | 0.03 (0.10) | 0.04 (0.03) | 0.14 (0.14) | 0.07 (0.07) | 0.07 (0.08) | 0.44 (0.39) | 0.08 (0.09) | 0.13 (0.10) |
| **100 tips** | | | | | | | | | | | | |
| constant | 0.79 (0.27) | 0.07 (0.02) | 0.08 (0.00) | 0.04 (0.02) | 0.01 (0.00) | 0.16 (0.07) | 0.05 (0.01) | 0.14 (0.03) | 0.11 (0.01) | 0.00 (0.00) | 0.24 (0.06) | 0.18 (0.06) |
| d. simple | 0.04 (0.07) | 0.28 (0.12) | 0.22 (0.08) | 0.15 (0.08) | 0.00 (0.01) | 0.04 (0.07) | 0.01 (0.03) | 0.11 (0.03) | 0.03 (0.03) | 0.01 (0.02) | 0.03 (0.04) | 0.07 (0.02) |
| d. threshold | 0.07 (0.28) | 0.22 (0.28) | 0.23 (0.31) | 0.22 (0.23) | 0.01 (0.02) | 0.04 (0.18) | 0.03 (0.03) | 0.05 (0.22) | 0.03 (0.17) | 0.01 (0.05) | 0.07 (0.20) | 0.04 (0.23) |
| d. sweetspot | 0.09 (0.29) | 0.38 (0.52) | 0.41 (0.54) | 0.53 (0.62) | 0.04 (0.09) | 0.11 (0.31) | 0.02 (0.09) | 0.12 (0.26) | 0.07 (0.22) | 0.01 (0.07) | 0.22 (0.36) | 0.09 (0.25) |
| o. simple | 0.00 (0.01) | 0.03 (0.03) | 0.00 (0.00) | 0.02 (0.00) | 0.82 (0.62) | 0.52 (0.21) | 0.19 (0.15) | 0.23 (0.10) | 0.38 (0.17) | 0.04 (0.01) | 0.04 (0.00) | 0.08 (0.05) |
| o. threshold | 0.01 (0.03) | 0.01 (0.00) | 0.03 (0.03) | 0.01 (0.02) | 0.04 (0.14) | 0.07 (0.07) | 0.40 (0.34) | 0.12 (0.18) | 0.18 (0.14) | 0.10 (0.10) | 0.02 (0.08) | 0.08 (0.12) |
| o. sweetspot | 0.00 (0.06) | 0.01 (0.03) | 0.03 (0.04) | 0.03 (0.03) | 0.07 (0.12) | 0.07 (0.08) | 0.31 (0.35) | 0.22 (0.17) | 0.19 (0.25) | 0.83 (0.75) | 0.38 (0.26) | 0.44 (0.27) |
| **200 tips** | | | | | | | | | | | | |
| constant | 0.72 (0.08) | 0.00 (0.00) | 0.03 (0.00) | 0.01 (0.00) | 0.00 (0.00) | 0.03 (0.00) | 0.00 (0.00) | 0.02 (0.00) | 0.01 (0.00) | 0.00 (0.00) | 0.03 (0.00) | 0.04 (0.00) |
| d. simple | 0.03 (0.07) | 0.18 (0.12) | 0.12 (0.07) | 0.10 (0.07) | 0.00 (0.00) | 0.03 (0.03) | 0.01 (0.00) | 0.02 (0.06) | 0.00 (0.02) | 0.00 (0.00) | 0.05 (0.01) | 0.01 (0.01) |
| d. threshold | 0.07 (0.24) | 0.29 (0.32) | 0.30 (0.28) | 0.26 (0.34) | 0.00 (0.00) | 0.01 (0.16) | 0.00 (0.01) | 0.03 (0.05) | 0.01 (0.05) | 0.00 (0.00) | 0.01 (0.12) | 0.00 (0.03) |
| d. sweetspot | 0.16 (0.52) | 0.48 (0.53) | 0.52 (0.56) | 0.61 (0.54) | 0.00 (0.02) | 0.03 (0.19) | 0.00 (0.03) | 0.04 (0.12) | 0.02 (0.08) | 0.01 (0.00) | 0.06 (0.19) | 0.03 (0.15) |
| o. simple | 0.00 (0.00) | 0.00 (0.00) | 0.00 (0.00) | 0.00 (0.00) | 0.85 (0.58) | 0.72 (0.36) | 0.05 (0.03) | 0.23 (0.19) | 0.24 (0.21) | 0.00 (0.00) | 0.02 (0.00) | 0.03 (0.01) |
| o. threshold | 0.01 (0.01) | 0.02 (0.01) | 0.01 (0.02) | 0.00 (0.02) | 0.07 (0.17) | 0.07 (0.11) | 0.65 (0.58) | 0.36 (0.25) | 0.35 (0.26) | 0.05 (0.10) | 0.12 (0.10) | 0.13 (0.19) |
| o. sweetspot | 0.02 (0.07) | 0.03 (0.02) | 0.03 (0.07) | 0.03 (0.03) | 0.08 (0.22) | 0.12 (0.16) | 0.29 (0.36) | 0.31 (0.32) | 0.38 (0.38) | 0.94 (0.90) | 0.72 (0.58) | 0.77 (0.62) |

Table S3. *Proportions of times (based on 120 replicates) our continuous stochastic character map-based pipeline for inferring relationships between rates of trait evolution and continuously-varying factors selected a model assuming a given kind of factor-rate relationship (i.e., constant, exponential/simple, logistic/threshold, or Gaussian/sweetspot; corresponding to different rows) as the best-fitting one (based on a summed AICc weight cutoff of 0.8). Here, d. and o. denote models based on simulated dummy factors and observed factors, respectively. Proportions for simulations with and without random variation around factor-rate relationships (noise) are given within and without parentheses, respectively.*

| | constant | hidden factor-dependent | | | simple | | threshold | | | sweetspot | | |
|---|---|---|---|---|---|---|---|---|---|---|---|---|
| | | simple | threshold | sweetspot | strong | weak | strong | weak | wide | strong | weak | wide |
| **50 tips** | | | | | | | | | | | | |
| constant | 0.84 (0.48) | 0.21 (0.11) | 0.20 (0.12) | 0.21 (0.07) | 0.11 (0.04) | 0.42 (0.24) | 0.09 (0.07) | 0.41 (0.17) | 0.24 (0.22) | 0.11 (0.01) | 0.48 (0.26) | 0.37 (0.18) |
| d. simple | 0.02 (0.06) | 0.32 (0.20) | 0.24 (0.16) | 0.23 (0.15) | 0.07 (0.05) | 0.03 (0.12) | 0.05 (0.04) | 0.03 (0.06) | 0.04 (0.04) | 0.03 (0.09) | 0.12 (0.10) | 0.06 (0.07) |
| d. threshold | 0.04 (0.22) | 0.07 (0.25) | 0.15 (0.22) | 0.11 (0.18) | 0.03 (0.13) | 0.04 (0.17) | 0.03 (0.08) | 0.05 (0.18) | 0.04 (0.18) | 0.05 (0.12) | 0.03 (0.17) | 0.03 (0.20) |
| d. sweetspot | 0.04 (0.18) | 0.31 (0.33) | 0.36 (0.42) | 0.38 (0.52) | 0.12 (0.07) | 0.11 (0.16) | 0.07 (0.18) | 0.08 (0.22) | 0.04 (0.17) | 0.17 (0.17) | 0.12 (0.20) | 0.13 (0.20) |
| o. simple | 0.03 (0.01) | 0.07 (0.03) | 0.02 (0.02) | 0.03 (0.01) | 0.61 (0.48) | 0.32 (0.17) | 0.46 (0.30) | 0.28 (0.14) | 0.45 (0.23) | 0.06 (0.04) | 0.07 (0.04) | 0.12 (0.08) |
| o. threshold | 0.02 (0.03) | 0.00 (0.02) | 0.01 (0.03) | 0.03 (0.03) | 0.03 (0.09) | 0.03 (0.08) | 0.15 (0.17) | 0.07 (0.12) | 0.09 (0.04) | 0.07 (0.12) | 0.03 (0.06) | 0.10 (0.10) |
| o. sweetspot | 0.01 (0.02) | 0.03 (0.07) | 0.03 (0.03) | 0.02 (0.04) | 0.04 (0.12) | 0.06 (0.07) | 0.16 (0.15) | 0.08 (0.11) | 0.09 (0.11) | 0.51 (0.45) | 0.14 (0.17) | 0.19 (0.16) |
| **100 tips** | | | | | | | | | | | | |
| constant | 0.78 (0.24) | 0.07 (0.02) | 0.08 (0.00) | 0.03 (0.02) | 0.01 (0.00) | 0.11 (0.05) | 0.04 (0.01) | 0.10 (0.03) | 0.09 (0.01) | 0.00 (0.00) | 0.20 (0.06) | 0.16 (0.06) |
| d. simple | 0.03 (0.07) | 0.27 (0.12) | 0.22 (0.07) | 0.15 (0.08) | 0.00 (0.01) | 0.03 (0.03) | 0.01 (0.02) | 0.08 (0.03) | 0.02 (0.03) | 0.00 (0.02) | 0.03 (0.03) | 0.06 (0.01) |
| d. threshold | 0.07 (0.27) | 0.22 (0.28) | 0.23 (0.30) | 0.22 (0.22) | 0.00 (0.02) | 0.02 (0.17) | 0.03 (0.02) | 0.03 (0.21) | 0.03 (0.12) | 0.00 (0.04) | 0.05 (0.19) | 0.03 (0.22) |
| d. sweetspot | 0.09 (0.28) | 0.38 (0.52) | 0.39 (0.51) | 0.52 (0.62) | 0.03 (0.07) | 0.07 (0.27) | 0.01 (0.07) | 0.09 (0.25) | 0.06 (0.20) | 0.01 (0.05) | 0.18 (0.30) | 0.05 (0.22) |
| o. simple | 0.00 (0.01) | 0.04 (0.03) | 0.01 (0.00) | 0.03 (0.00) | 0.84 (0.64) | 0.62 (0.24) | 0.20 (0.16) | 0.30 (0.10) | 0.42 (0.19) | 0.04 (0.01) | 0.05 (0.03) | 0.10 (0.05) |
| o. threshold | 0.01 (0.04) | 0.01 (0.01) | 0.03 (0.03) | 0.01 (0.02) | 0.04 (0.14) | 0.07 (0.08) | 0.40 (0.38) | 0.17 (0.19) | 0.18 (0.16) | 0.11 (0.11) | 0.03 (0.11) | 0.12 (0.13) |
| o. sweetspot | 0.03 (0.10) | 0.02 (0.04) | 0.03 (0.08) | 0.04 (0.04) | 0.08 (0.12) | 0.08 (0.15) | 0.32 (0.36) | 0.23 (0.20) | 0.21 (0.28) | 0.84 (0.78) | 0.47 (0.28) | 0.49 (0.31) |
| **200 tips** | | | | | | | | | | | | |
| constant | 0.70 (0.08) | 0.00 (0.00) | 0.03 (0.00) | 0.01 (0.00) | 0.00 (0.00) | 0.02 (0.00) | 0.00 (0.00) | 0.01 (0.00) | 0.01 (0.00) | 0.00 (0.00) | 0.01 (0.00) | 0.02 (0.00) |
| d. simple | 0.02 (0.07) | 0.18 (0.11) | 0.12 (0.07) | 0.10 (0.07) | 0.00 (0.00) | 0.02 (0.03) | 0.00 (0.00) | 0.02 (0.05) | 0.00 (0.02) | 0.00 (0.00) | 0.03 (0.01) | 0.00 (0.01) |
| d. threshold | 0.07 (0.24) | 0.29 (0.31) | 0.30 (0.28) | 0.26 (0.34) | 0.00 (0.00) | 0.00 (0.15) | 0.00 (0.01) | 0.01 (0.04) | 0.01 (0.05) | 0.00 (0.00) | 0.01 (0.10) | 0.00 (0.02) |
| d. sweetspot | 0.16 (0.49) | 0.48 (0.52) | 0.52 (0.56) | 0.60 (0.54) | 0.00 (0.02) | 0.02 (0.17) | 0.00 (0.02) | 0.02 (0.11) | 0.01 (0.07) | 0.01 (0.00) | 0.03 (0.17) | 0.01 (0.12) |
| o. simple | 0.01 (0.00) | 0.00 (0.00) | 0.00 (0.00) | 0.00 (0.00) | 0.85 (0.58) | 0.76 (0.37) | 0.05 (0.03) | 0.26 (0.19) | 0.24 (0.21) | 0.00 (0.00) | 0.02 (0.00) | 0.03 (0.01) |
| o. threshold | 0.02 (0.03) | 0.02 (0.03) | 0.01 (0.02) | 0.00 (0.02) | 0.07 (0.17) | 0.07 (0.12) | 0.65 (0.59) | 0.38 (0.26) | 0.36 (0.26) | 0.05 (0.10) | 0.13 (0.10) | 0.14 (0.21) |
| o. sweetspot | 0.03 (0.07) | 0.03 (0.04) | 0.03 (0.07) | 0.03 (0.03) | 0.08 (0.22) | 0.12 (0.17) | 0.30 (0.36) | 0.32 (0.35) | 0.38 (0.39) | 0.94 (0.90) | 0.78 (0.62) | 0.80 (0.64) |

Table S4. *Proportions of times (based on 120 replicates) our continuous stochastic character map-based pipeline for inferring relationships between rates of trait evolution and continuously-varying factors selected a model assuming a given kind of factor-rate relationship (i.e., null, exponential/simple, logistic/threshold, or Gaussian/sweetspot; corresponding to different rows) as the best-fitting one (based on a summed AICc weight cutoff of 0.8). Proportions for simulations with and without random variation around simulated factor-rate relationships (noise) are given within and without parentheses, respectively. Here, we consider all models assuming either constant or dummy factor-dependent rates "null".*

| | constant | hidden factor-dependent | | | simple | | threshold | | | sweetspot | | |
|---|---|---|---|---|---|---|---|---|---|---|---|---|
| | | simple | threshold | sweetspot | strong | weak | strong | weak | wide | strong | weak | wide |
| **50 tips** | | | | | | | | | | | | |
| null | 0.94 (0.94) | 0.91 (0.89) | 0.95 (0.92) | 0.92 (0.92) | 0.32 (0.30) | 0.60 (0.68) | 0.23 (0.38) | 0.57 (0.63) | 0.37 (0.62) | 0.36 (0.39) | 0.75 (0.73) | 0.58 (0.66) |
| simple | 0.03 (0.01) | 0.07 (0.03) | 0.02 (0.02) | 0.03 (0.01) | 0.61 (0.48) | 0.32 (0.17) | 0.46 (0.30) | 0.28 (0.14) | 0.45 (0.23) | 0.06 (0.04) | 0.07 (0.04) | 0.12 (0.08) |
| threshold | 0.02 (0.03) | 0.00 (0.02) | 0.01 (0.03) | 0.03 (0.03) | 0.03 (0.09) | 0.03 (0.08) | 0.15 (0.17) | 0.07 (0.12) | 0.09 (0.04) | 0.07 (0.12) | 0.03 (0.06) | 0.10 (0.10) |
| sweetspot | 0.01 (0.02) | 0.03 (0.07) | 0.03 (0.03) | 0.02 (0.04) | 0.04 (0.12) | 0.06 (0.07) | 0.16 (0.15) | 0.08 (0.11) | 0.09 (0.11) | 0.51 (0.45) | 0.14 (0.17) | 0.19 (0.16) |
| **100 tips** | | | | | | | | | | | | |
| null | 0.97 (0.85) | 0.93 (0.92) | 0.92 (0.88) | 0.92 (0.94) | 0.03 (0.09) | 0.22 (0.52) | 0.08 (0.11) | 0.30 (0.51) | 0.19 (0.37) | 0.01 (0.11) | 0.46 (0.58) | 0.29 (0.51) |
| simple | 0.00 (0.01) | 0.04 (0.03) | 0.01 (0.00) | 0.03 (0.00) | 0.84 (0.64) | 0.62 (0.24) | 0.20 (0.16) | 0.30 (0.10) | 0.42 (0.19) | 0.04 (0.01) | 0.05 (0.03) | 0.10 (0.05) |
| threshold | 0.01 (0.04) | 0.01 (0.01) | 0.03 (0.03) | 0.01 (0.02) | 0.04 (0.14) | 0.07 (0.08) | 0.40 (0.38) | 0.17 (0.19) | 0.18 (0.16) | 0.11 (0.11) | 0.03 (0.11) | 0.12 (0.13) |
| sweetspot | 0.03 (0.10) | 0.02 (0.04) | 0.03 (0.08) | 0.04 (0.04) | 0.08 (0.12) | 0.08 (0.15) | 0.32 (0.36) | 0.23 (0.20) | 0.21 (0.28) | 0.84 (0.78) | 0.47 (0.28) | 0.49 (0.31) |
| **200 tips** | | | | | | | | | | | | |
| null | 0.94 (0.89) | 0.96 (0.93) | 0.96 (0.91) | 0.97 (0.96) | 0.00 (0.02) | 0.05 (0.34) | 0.00 (0.03) | 0.05 (0.20) | 0.03 (0.14) | 0.01 (0.00) | 0.07 (0.28) | 0.03 (0.14) |
| simple | 0.01 (0.00) | 0.00 (0.00) | 0.00 (0.00) | 0.00 (0.00) | 0.85 (0.58) | 0.76 (0.37) | 0.05 (0.03) | 0.26 (0.19) | 0.24 (0.21) | 0.00 (0.00) | 0.02 (0.00) | 0.03 (0.01) |
| threshold | 0.02 (0.03) | 0.02 (0.03) | 0.01 (0.02) | 0.00 (0.02) | 0.07 (0.17) | 0.07 (0.12) | 0.65 (0.59) | 0.38 (0.26) | 0.36 (0.26) | 0.05 (0.10) | 0.13 (0.10) | 0.14 (0.21) |
| sweetspot | 0.03 (0.07) | 0.03 (0.04) | 0.03 (0.07) | 0.03 (0.03) | 0.08 (0.22) | 0.12 (0.17) | 0.30 (0.36) | 0.32 (0.35) | 0.38 (0.39) | 0.94 (0.90) | 0.78 (0.62) | 0.80 (0.64) |

Table S5. *Proportions of times (based on 120 replicates) our continuous stochastic character map-based pipeline for inferring relationships between rates of trait evolution and continuously-varying factors selected a model assuming a given kind of factor-rate relationship (i.e., constant, exponential/simple, logistic/threshold, or Gaussian/sweetspot; corresponding to different rows) as the best-fitting one (based on a summed AICc weight cutoff of 0.7). Here, d. and o. denote models based on simulated dummy factors and observed factors, respectively. Proportions for simulations with and without random variation around factor-rate relationships (noise) are given within and without parentheses, respectively.*

| | constant | hidden factor-dependent | | | simple | | threshold | | | sweetspot | | |
|---|---|---|---|---|---|---|---|---|---|---|---|---|
| | | simple | threshold | sweetspot | strong | weak | strong | weak | wide | strong | weak | wide |
| **50 tips** | | | | | | | | | | | | |
| constant | 0.80 (0.43) | 0.20 (0.11) | 0.17 (0.11) | 0.18 (0.07) | 0.07 (0.03) | 0.35 (0.16) | 0.08 (0.06) | 0.34 (0.16) | 0.16 (0.19) | 0.07 (0.01) | 0.41 (0.21) | 0.32 (0.17) |
| d. simple | 0.02 (0.05) | 0.28 (0.18) | 0.24 (0.16) | 0.22 (0.12) | 0.06 (0.03) | 0.02 (0.09) | 0.03 (0.03) | 0.03 (0.04) | 0.03 (0.03) | 0.03 (0.06) | 0.10 (0.09) | 0.03 (0.06) |
| d. threshold | 0.04 (0.22) | 0.06 (0.23) | 0.14 (0.21) | 0.11 (0.18) | 0.03 (0.12) | 0.04 (0.15) | 0.03 (0.08) | 0.04 (0.17) | 0.03 (0.16) | 0.04 (0.12) | 0.03 (0.17) | 0.01 (0.17) |
| d. sweetspot | 0.04 (0.17) | 0.30 (0.32) | 0.33 (0.42) | 0.37 (0.47) | 0.07 (0.06) | 0.09 (0.13) | 0.05 (0.16) | 0.06 (0.19) | 0.04 (0.16) | 0.14 (0.14) | 0.12 (0.20) | 0.11 (0.17) |
| o. simple | 0.05 (0.03) | 0.12 (0.03) | 0.05 (0.03) | 0.03 (0.01) | 0.68 (0.54) | 0.40 (0.26) | 0.48 (0.31) | 0.36 (0.18) | 0.52 (0.25) | 0.07 (0.05) | 0.12 (0.05) | 0.16 (0.09) |
| o. threshold | 0.03 (0.05) | 0.00 (0.02) | 0.03 (0.03) | 0.03 (0.04) | 0.03 (0.10) | 0.04 (0.11) | 0.15 (0.17) | 0.08 (0.13) | 0.10 (0.07) | 0.07 (0.12) | 0.05 (0.08) | 0.11 (0.14) |
| o. sweetspot | 0.03 (0.04) | 0.04 (0.10) | 0.03 (0.05) | 0.05 (0.11) | 0.05 (0.12) | 0.06 (0.10) | 0.18 (0.18) | 0.09 (0.12) | 0.13 (0.14) | 0.58 (0.49) | 0.18 (0.20) | 0.26 (0.20) |
| **100 tips** | | | | | | | | | | | | |
| constant | 0.77 (0.19) | 0.07 (0.02) | 0.07 (0.00) | 0.03 (0.01) | 0.01 (0.00) | 0.09 (0.03) | 0.03 (0.01) | 0.07 (0.02) | 0.07 (0.01) | 0.00 (0.00) | 0.13 (0.04) | 0.11 (0.05) |
| d. simple | 0.03 (0.07) | 0.25 (0.11) | 0.22 (0.07) | 0.15 (0.08) | 0.00 (0.01) | 0.03 (0.03) | 0.01 (0.02) | 0.07 (0.02) | 0.01 (0.02) | 0.00 (0.02) | 0.03 (0.03) | 0.05 (0.01) |
| d. threshold | 0.06 (0.26) | 0.20 (0.27) | 0.22 (0.29) | 0.21 (0.21) | 0.00 (0.02) | 0.02 (0.13) | 0.03 (0.02) | 0.03 (0.20) | 0.02 (0.11) | 0.00 (0.04) | 0.04 (0.17) | 0.03 (0.19) |
| d. sweetspot | 0.07 (0.27) | 0.37 (0.51) | 0.38 (0.51) | 0.49 (0.60) | 0.02 (0.06) | 0.05 (0.22) | 0.00 (0.07) | 0.08 (0.23) | 0.04 (0.19) | 0.01 (0.05) | 0.12 (0.21) | 0.05 (0.18) |
| o. simple | 0.00 (0.01) | 0.06 (0.03) | 0.02 (0.00) | 0.03 (0.01) | 0.85 (0.64) | 0.63 (0.28) | 0.22 (0.16) | 0.32 (0.12) | 0.44 (0.20) | 0.04 (0.01) | 0.07 (0.04) | 0.11 (0.06) |
| o. threshold | 0.03 (0.07) | 0.02 (0.02) | 0.04 (0.04) | 0.03 (0.03) | 0.04 (0.14) | 0.07 (0.10) | 0.40 (0.38) | 0.17 (0.19) | 0.20 (0.18) | 0.11 (0.11) | 0.05 (0.12) | 0.12 (0.13) |
| o. sweetspot | 0.04 (0.13) | 0.04 (0.05) | 0.05 (0.09) | 0.07 (0.07) | 0.08 (0.13) | 0.10 (0.20) | 0.32 (0.36) | 0.26 (0.22) | 0.22 (0.29) | 0.84 (0.78) | 0.55 (0.38) | 0.53 (0.38) |
| **200 tips** | | | | | | | | | | | | |
| constant | 0.67 (0.07) | 0.00 (0.00) | 0.03 (0.00) | 0.01 (0.00) | 0.00 (0.00) | 0.02 (0.00) | 0.00 (0.00) | 0.01 (0.00) | 0.01 (0.00) | 0.00 (0.00) | 0.00 (0.00) | 0.02 (0.00) |
| d. simple | 0.01 (0.07) | 0.18 (0.11) | 0.12 (0.07) | 0.10 (0.07) | 0.00 (0.00) | 0.01 (0.03) | 0.00 (0.00) | 0.01 (0.04) | 0.00 (0.02) | 0.00 (0.00) | 0.01 (0.01) | 0.00 (0.01) |
| d. threshold | 0.07 (0.22) | 0.29 (0.30) | 0.30 (0.28) | 0.26 (0.33) | 0.00 (0.00) | 0.00 (0.14) | 0.00 (0.01) | 0.01 (0.03) | 0.01 (0.04) | 0.00 (0.00) | 0.01 (0.09) | 0.00 (0.02) |
| d. sweetspot | 0.15 (0.47) | 0.48 (0.52) | 0.51 (0.55) | 0.60 (0.54) | 0.00 (0.02) | 0.02 (0.14) | 0.00 (0.02) | 0.00 (0.09) | 0.01 (0.07) | 0.00 (0.00) | 0.03 (0.16) | 0.00 (0.11) |
| o. simple | 0.02 (0.01) | 0.00 (0.00) | 0.00 (0.00) | 0.00 (0.00) | 0.85 (0.58) | 0.77 (0.38) | 0.05 (0.03) | 0.27 (0.20) | 0.24 (0.22) | 0.00 (0.00) | 0.03 (0.00) | 0.03 (0.01) |
| o. threshold | 0.03 (0.04) | 0.02 (0.03) | 0.01 (0.02) | 0.00 (0.02) | 0.07 (0.17) | 0.07 (0.13) | 0.65 (0.59) | 0.38 (0.27) | 0.36 (0.27) | 0.06 (0.10) | 0.13 (0.10) | 0.15 (0.21) |
| o. sweetspot | 0.06 (0.12) | 0.03 (0.05) | 0.04 (0.08) | 0.03 (0.03) | 0.08 (0.22) | 0.12 (0.18) | 0.30 (0.36) | 0.33 (0.38) | 0.38 (0.39) | 0.94 (0.90) | 0.80 (0.64) | 0.80 (0.65) |

Table 6. *Proportions of times our continuous stochastic character map-based pipeline for inferring relationships between rates of trait evolution and continuously-varying factors selected a model assuming a given kind of factor-rate relationship (i.e., null, exponential/simple, logistic/threshold, or Gaussian/sweetspot; corresponding to different rows) as the best-fitting one (based on a summed AICc weight cutoff of 0.7). Proportions for simulations with and without random variation around simulated factor-rate relationships (noise) are given within and without parentheses, respectively. Here, we consider all models assuming either constant or dummy factor-dependent rates "null".*

| | constant | hidden factor-dependent | | | simple | | threshold | | | sweetspot | | |
|---|---|---|---|---|---|---|---|---|---|---|---|---|
| | | simple | threshold | sweetspot | strong | weak | strong | weak | wide | strong | weak | wide |
| **50 tips** | | | | | | | | | | | | |
| null | 0.90 (0.88) | 0.83 (0.85) | 0.89 (0.89) | 0.88 (0.84) | 0.23 (0.23) | 0.50 (0.53) | 0.19 (0.33) | 0.47 (0.56) | 0.25 (0.54) | 0.28 (0.33) | 0.65 (0.67) | 0.48 (0.57) |
| simple | 0.05 (0.03) | 0.12 (0.03) | 0.05 (0.03) | 0.03 (0.01) | 0.68 (0.54) | 0.40 (0.26) | 0.48 (0.31) | 0.36 (0.18) | 0.52 (0.25) | 0.07 (0.05) | 0.12 (0.05) | 0.16 (0.09) |
| threshold | 0.03 (0.05) | 0.00 (0.02) | 0.03 (0.03) | 0.03 (0.04) | 0.03 (0.10) | 0.04 (0.11) | 0.15 (0.17) | 0.08 (0.13) | 0.10 (0.07) | 0.07 (0.12) | 0.05 (0.08) | 0.11 (0.14) |
| sweetspot | 0.03 (0.04) | 0.04 (0.10) | 0.03 (0.05) | 0.05 (0.11) | 0.05 (0.12) | 0.06 (0.10) | 0.18 (0.18) | 0.09 (0.12) | 0.13 (0.14) | 0.58 (0.49) | 0.18 (0.20) | 0.26 (0.20) |
| **100 tips** | | | | | | | | | | | | |
| null | 0.93 (0.78) | 0.88 (0.90) | 0.89 (0.87) | 0.88 (0.90) | 0.03 (0.08) | 0.19 (0.42) | 0.07 (0.11) | 0.25 (0.47) | 0.13 (0.32) | 0.01 (0.11) | 0.32 (0.46) | 0.23 (0.43) |
| simple | 0.00 (0.01) | 0.06 (0.03) | 0.02 (0.00) | 0.03 (0.01) | 0.85 (0.64) | 0.63 (0.28) | 0.22 (0.16) | 0.32 (0.12) | 0.44 (0.20) | 0.04 (0.01) | 0.07 (0.04) | 0.11 (0.06) |
| threshold | 0.03 (0.07) | 0.02 (0.02) | 0.04 (0.04) | 0.03 (0.03) | 0.04 (0.14) | 0.07 (0.10) | 0.40 (0.38) | 0.17 (0.19) | 0.20 (0.18) | 0.11 (0.11) | 0.05 (0.12) | 0.12 (0.13) |
| sweetspot | 0.04 (0.13) | 0.04 (0.05) | 0.05 (0.09) | 0.07 (0.07) | 0.08 (0.13) | 0.10 (0.20) | 0.32 (0.36) | 0.26 (0.22) | 0.22 (0.29) | 0.84 (0.78) | 0.55 (0.38) | 0.53 (0.38) |
| **200 tips** | | | | | | | | | | | | |
| null | 0.89 (0.83) | 0.95 (0.92) | 0.95 (0.90) | 0.97 (0.95) | 0.00 (0.02) | 0.04 (0.31) | 0.00 (0.03) | 0.03 (0.16) | 0.03 (0.12) | 0.00 (0.00) | 0.04 (0.26) | 0.02 (0.13) |
| simple | 0.02 (0.01) | 0.00 (0.00) | 0.00 (0.00) | 0.00 (0.00) | 0.85 (0.58) | 0.77 (0.38) | 0.05 (0.03) | 0.27 (0.20) | 0.24 (0.22) | 0.00 (0.00) | 0.03 (0.00) | 0.03 (0.01) |
| threshold | 0.03 (0.04) | 0.02 (0.03) | 0.01 (0.02) | 0.00 (0.02) | 0.07 (0.17) | 0.07 (0.13) | 0.65 (0.59) | 0.38 (0.27) | 0.36 (0.27) | 0.06 (0.10) | 0.13 (0.10) | 0.15 (0.21) |
| sweetspot | 0.06 (0.12) | 0.03 (0.05) | 0.04 (0.08) | 0.03 (0.03) | 0.08 (0.22) | 0.12 (0.18) | 0.30 (0.36) | 0.33 (0.38) | 0.38 (0.39) | 0.94 (0.90) | 0.80 (0.64) | 0.80 (0.65) |

Table S7. *Maximum likelihood parameter estimates and associated AICc for models of eucalpyt* ***peduncle length*** *evolution. Here, d. and h. denote models based on simulated dummy factors and maximum height, respectively. Parenthesized numbers represent lower and upper ends of 95% confidence intervals calculated via dentist (Boyko and O'Meara, 2024). All rates are per million years; height/dummy factor measured in millimeters on natural log scale ($\beta_0$ corresponds to rate at mean height/dummy factor value, ~9.6 or 15 meters).*

| | parameter | | | | | | | |
|---|---|---|---|---|---|---|---|---|
| model | intercept ($\beta_0$) | slope ($\beta_1$) | mid-rate ($\alpha$) | location ($\theta$) | rate deviation ($\delta$) | width ($\omega$) | tip error ($\epsilon$) | AICc |
| constant | — | — | -5.15 (-5.94, -4.50) | — | — | — | 0.46 (0.40, 0.53) | 516.90 |
| h. simple | -5.12 (-6.12, -4.41) | -0.32 (-1.84, 1.67) | — | — | — | — | 0.46 (0.38, 0.54) | 518.67 |
| h. threshold | — | — | -5.64 (-6.56, -4.52) | 10.07 (8.93, 10.66) | -10.00 (-10.00, 10.00) | -1.84 (-2.61, 3.60) | 0.46 (0.37, 0.55) | 521.34 |
| h. sweetspot | — | — | -3.96 (-6.76, -3.86) | 9.78 (7.58, 14.10) | 1.18 (-10.00, 9.91) | -1.45 (-2.61, 3.59) | 0.46 (0.38, 0.56) | 520.24 |
| d. simple | -3.98 | 1.89 | — | — | — | — | 0.45 | 514.52 |
| d. threshold | — | — | 6.69 | 6.55 | -6.23 | -1.76 | 0.43 | 510.63 |
| d. sweetspot | — | — | 1.22 | 9.89 | -4.86 | 4.36 | 0.36 | ***497.14*** |

Table S8. *Maximum likelihood parameter estimates and associated AICc for models of eucalpyt* ***inflorescence pedicel length*** *evolution. Here, d. and h. denote models based on simulated dummy factors and maximum height, respectively. Parenthesized numbers represent lower and upper ends of 95% confidence intervals calculated via dentist (Boyko and O'Meara, 2024). All rates are per million years; height/dummy factor measured in millimeters on natural log scale ($\beta_0$ corresponds to rate at mean height/dummy factor value, ~9.6 or 15 meters).*

| | parameter | | | | | | | |
|---|---|---|---|---|---|---|---|---|
| model | intercept ($\beta_0$) | slope ($\beta_1$) | mid-rate ($\alpha$) | location ($\theta$) | rate deviation ($\delta$) | width ($\omega$) | tip error ($\epsilon$) | AICc |
| constant | — | — | -4.59 (-5.49, -3.91) | — | — | — | 0.56 (0.46, 0.65) | 647.77 |
| h. simple | -4.66 (-6.50, -3.83) | 0.36 (-2.00, 2.78) | — | — | — | — | 0.56 (0.45, 0.67) | 649.67 |
| h. threshold | — | — | -4.36 (-5.11, -4.10) | 9.83 (9.03, 10.15) | 0.85 (-4.52, 1.23) | -2.62 (-2.62, 3.60) | 0.57 (0.46, 0.66) | 651.54 |
| h. sweetspot | — | — | -3.70 (-6.10, -0.84) | 10.11 (2.22, 16.83) | 1.38 (-6.20, 2.33) | -0.33 (-1.68, 1.10) | 0.58 (0.47, 0.68) | 648.38 |
| d. simple | -6.40 | 2.18 | — | — | — | — | 0.54 | ***631.50*** |
| d. threshold | — | — | -1.30 | 11.71 | 3.20 | 1.12 | 0.55 | **631.59** |
| d. sweetspot | — | — | -1.33 | 12.28 | 2.61 | 1.21 | 0.54 | **631.80** |

Table S9. *Maximum likelihood parameter estimates and associated AICc for models of eucalpyt* ***bud length*** *evolution. Here, d. and h. denote models based on simulated dummy factors and maximum height, respectively. Parenthesized numbers represent lower and upper ends of 95% confidence intervals calculated via dentist (Boyko and O'Meara, 2024). All rates are per million years; height/dummy factor measured in millimeters on natural log scale ($\beta_0$ corresponds to rate at mean height/dummy factor value, ~9.6 or 15 meters).*

| | parameter | | | | | | | |
|---|---|---|---|---|---|---|---|---|
| model | intercept ($\beta_0$) | slope ($\beta_1$) | mid-rate ($\alpha$) | location ($\theta$) | rate deviation ($\delta$) | width ($\omega$) | tip error ($\epsilon$) | AICc |
| constant | — | — | -4.73 (-5.36, -4.23) | — | — | — | 0.29 (0.22, 0.37) | 379.10 |
| h. simple | -4.91 (-5.83, -4.30) | -0.97 (-2.24, -0.01) | — | — | — | — | 0.30 (0.22, 0.37) | 373.56 |
| h. threshold | — | — | -4.39 (-5.57, -2.94) | 9.15 (6.48, 10.60) | -0.72 (-4.33, -0.05) | -0.96 (-2.47, 2.11) | 0.29 (0.20, 0.39) | 373.09 |
| h. sweetspot | — | — | -3.87 (-5.10, -2.42) | 8.83 (7.96, 9.12) | 1.00 (0.25, 2.01) | -0.18 (-2.59, 2.05) | 0.29 (0.20, 0.38) | ***370.06*** |
| d. simple | -5.89 | -0.84 | — | — | — | — | 0.30 | 374.67 |
| d. threshold | — | — | 7.90 | -6.45 | -7.50 | 2.47 | 0.29 | 377.99 |
| d. sweetspot | — | — | -3.76 | 6.63 | 1.11 | 1.46 | 0.28 | 374.82 |

Table S10. *Maximum likelihood parameter estimates and associated AICc for models of eucalpyt* ***bud width*** *evolution. Here, d. and h. denote models based on simulated dummy factors and maximum height, respectively. Parenthesized numbers represent lower and upper ends of 95% confidence intervals calculated via dentist (Boyko and O'Meara, 2024). All rates are per million years; height/dummy factor measured in millimeters on natural log scale ($\beta_0$ corresponds to rate at mean height/dummy factor value, ~9.6 or 15 meters).*

| | parameter | | | | | | | |
|---|---|---|---|---|---|---|---|---|
| model | intercept ($\beta_0$) | slope ($\beta_1$) | mid-rate ($\alpha$) | location ($\theta$) | rate deviation ($\delta$) | width ($\omega$) | tip error ($\epsilon$) | AICc |
| constant | — | — | -5.20 (-5.89, -4.62) | — | — | — | 0.29 (0.22, 0.35) | 310.30 |
| h. simple | -5.90 (-7.21, -4.91) | -1.88 (-3.60, -0.54) | — | — | — | — | 0.30 (0.24, 0.36) | 297.01 |
| h. threshold | — | — | -4.93 (-6.05, -4.11) | 9.22 (8.90, 9.78) | -1.38 (-10.00, -0.43) | -0.87 (-1.95, -0.46) | 0.30 (0.25, 0.37) | 296.50 |
| h. sweetspot | — | — | -4.40 (-5.02, -0.88) | 9.77 (9.50, 11.10) | -9.99 (-9.99, -0.81) | 1.40 (0.43, 3.60) | 0.27 (0.21, 0.35) | ***293.38*** |
| d. simple | -5.96 | 0.96 | — | — | — | — | 0.28 | 301.82 |
| d. threshold | — | — | -4.80 | 9.67 | 1.12 | -1.85 | 0.29 | 304.29 |
| d. sweetspot | — | — | -4.54 | 9.13 | -10.00 | 1.56 | 0.24 | 297.70 |

Table S11. *Maximum likelihood parameter estimates and associated AICc for models of eucalypt* ***infructescence pedicel length*** *evolution. Here, d. and h. denote models based on simulated dummy factors and maximum height, respectively. Parenthesized numbers represent lower and upper ends of 95% confidence intervals calculated via dentist (Boyko and O'Meara, 2024). All rates are per million years; height/dummy factor measured in millimeters on natural log scale ($\beta_0$ corresponds to rate at mean height/dummy factor value, ~9.6 or 15 meters).*

| | parameter | | | | | | | |
|---|---|---|---|---|---|---|---|---|
| model | intercept ($\beta_0$) | slope ($\beta_1$) | mid-rate ($\alpha$) | location ($\theta$) | rate deviation ($\delta$) | width ($\omega$) | tip error ($\epsilon$) | AICc |
| constant | — | — | -4.33 (-5.28, -3.60) | — | — | — | 0.64 (0.53, 0.75) | 736.74 |
| h. simple | -4.25 (-5.72, -3.42) | -0.41 (-2.02, 2.08) | — | — | — | — | 0.63 (0.47, 0.77) | 738.52 |
| h. threshold | — | — | -4.82 (-6.36, -2.08) | 10.16 (5.92, 12.86) | -9.36 (-9.98, 9.99) | -2.08 (-2.61, 3.60) | 0.63 (0.48, 0.79) | 742.05 |
| h. sweetspot | — | — | -2.91 (-6.25, -2.01) | 9.89 (8.20, 10.53) | 1.43 (-10.00, 9.99) | -1.57 (-2.53, 3.54) | 0.66 (0.50, 0.78) | 741.28 |
| d. simple | -5.30 | 1.90 | — | — | — | — | 0.59 | ***734.31*** |
| d. threshold | — | — | -1.57 | 11.29 | 1.78 | 0.49 | 0.56 | **734.69** |
| d. sweetspot | — | — | 2.20 | 15.43 | 3.94 | 2.06 | 0.57 | **736.20** |

Table S12. *Maximum likelihood parameter estimates and associated AICc for models of eucalypt* ***fruit length*** *evolution. Here, d. and h. denote models based on simulated dummy factors and maximum height, respectively. Parenthesized numbers represent lower and upper ends of 95% confidence intervals calculated via dentist (Boyko and O'Meara, 2024). All rates are per million years; height/dummy factor measured in millimeters on natural log scale ($\beta_0$ corresponds to rate at mean height/dummy factor value, ~9.6 or 15 meters).*

| | parameter | | | | | | | |
|---|---|---|---|---|---|---|---|---|
| model | intercept ($\beta_0$) | slope ($\beta_1$) | mid-rate ($\alpha$) | location ($\theta$) | rate deviation ($\delta$) | width ($\omega$) | tip error ($\epsilon$) | AICc |
| constant | — | — | -4.92 (-5.51, -4.42) | — | — | — | 0.28 (0.21, 0.34) | 333.32 |
| h. simple | -4.92 (-5.64, -4.36) | -0.30 (-1.43, 0.82) | — | — | — | — | 0.28 (0.20, 0.35) | 334.74 |
| h. threshold | — | — | -4.89 (-5.76, -3.91) | 9.49 (8.91, 10.50) | -0.36 (-4.03, 1.07) | -2.62 (-2.62, 3.60) | 0.27 (0.19, 0.36) | 337.46 |
| h. sweetspot | — | — | -5.17 (-6.12, -4.58) | 9.70 (8.97, 9.90) | -10.00 (-10.00, 9.80) | 0.11 (-2.54, 1.96) | 0.27 (0.21, 0.35) | 334.25 |
| d. simple | -5.24 | 0.68 | — | — | — | — | 0.26 | 333.17 |
| d. threshold | — | — | -4.62 | 10.27 | 0.83 | 0.45 | 0.24 | 332.47 |
| d. sweetspot | — | — | -4.46 | 9.10 | -1.53 | 1.45 | 0.22 | ***328.28*** |

Table S13. *Maximum likelihood parameter estimates and associated AICc for models of eucalpyt* ***fruit width*** *evolution. Here, d. and h. denote models based on simulated dummy factors and maximum height, respectively. Parenthesized numbers represent lower and upper ends of 95% confidence intervals calculated via dentist (Boyko and O'Meara, 2024). All rates are per million years; height/dummy factor measured in millimeters on natural log scale ($\beta_0$ corresponds to rate at mean height/dummy factor value, ~9.6 or 15 meters).*

| | parameter | | | | | | | |
|---|---|---|---|---|---|---|---|---|
| model | intercept ($\beta_0$) | slope ($\beta_1$) | mid-rate ($\alpha$) | location ($\theta$) | rate deviation ($\delta$) | width ($\omega$) | tip error ($\epsilon$) | AICc |
| constant | — | — | -4.88 (-5.41, -4.45) | — | — | — | 0.24 (0.18, 0.30) | 287.75 |
| h. simple | -5.02 (-5.85, -4.49) | -0.70 (-1.89, 0.23) | — | — | — | — | 0.24 (0.18, 0.31) | 285.43 |
| h. threshold | — | — | -4.45 (-5.90, -4.21) | 8.95 (4.09, 16.83) | -0.58 (-3.20, 0.56) | -1.49 (-2.62, 3.59) | 0.23 (0.17, 0.31) | 286.34 |
| h. sweetspot | — | — | -4.96 (-5.85, -2.25) | 9.70 (9.10, 12.13) | -1.42 (-10.00, -0.01) | 0.60 (-2.55, 3.60) | 0.23 (0.15, 0.31) | 283.96 |
| d. simple | -5.34 | 0.97 | — | — | — | — | 0.22 | 286.21 |
| d. threshold | — | — | -5.26 | 9.17 | 10.00 | -1.85 | 0.23 | 279.66 |
| d. sweetspot | — | — | -4.70 | 9.18 | -10.00 | 1.25 | 0.20 | ***275.21*** |

Table S14. *Maximum likelihood parameter estimates and associated AICc for models of eucalpyt* ***seed length*** *evolution. Here, d. and h. denote models based on simulated dummy factors and maximum height, respectively. Parenthesized numbers represent lower and upper ends of 95% confidence intervals calculated via dentist (Boyko and O'Meara, 2024). All rates are per million years; height/dummy factor measured in millimeters on natural log scale ($\beta_0$ corresponds to rate at mean height/dummy factor value, ~9.6 or 15 meters).*

| | parameter | | | | | | | |
|---|---|---|---|---|---|---|---|---|
| model | intercept ($\beta_0$) | slope ($\beta_1$) | mid-rate ($\alpha$) | location ($\theta$) | rate deviation ($\delta$) | width ($\omega$) | tip error ($\epsilon$) | AICc |
| constant | — | — | -5.21 (-5.60, -4.87) | — | — | — | 0.12 (0.07, 0.16) | ***54.86*** |
| h. simple | -5.21 (-5.66, -4.82) | 0.18 (-0.44, 0.82) | — | — | — | — | 0.11 (0.06, 0.17) | **56.24** |
| h. threshold | — | — | -5.04 (-5.62, -3.91) | 10.02 (8.74, 16.83) | 0.22 (-0.19, 0.90) | -2.62 (-2.62, 3.60) | 0.11 (0.05, 0.18) | 59.97 |
| h. sweetspot | — | — | -4.39 (-4.49, -2.68) | 11.79 (9.83, 16.00) | 0.67 (0.52, 1.58) | 1.42 (-2.62, 2.41) | 0.11 (0.05, 0.18) | 60.02 |
| d. simple | -5.21 | 0.00 | — | — | — | — | 0.12 | 56.90 |
| d. threshold | — | — | -5.90 | 15.23 | -10.00 | -1.85 | 0.12 | 60.88 |
| d. sweetspot | — | — | -5.84 | 11.04 | -10.00 | -0.48 | 0.11 | 60.56 |

Table S15. *Maximum likelihood parameter estimates and associated AICc for models of eucalpyt* ***petiole length*** *evolution. Here, d. and h. denote models based on simulated dummy factors and maximum height, respectively. Parenthesized numbers represent lower and upper ends of 95% confidence intervals calculated via dentist (Boyko and O'Meara, 2024). All rates are per million years; height/dummy factor measured in millimeters on natural log scale ($\beta_0$ corresponds to rate at mean height/dummy factor value, ~9.6 or 15 meters).*

| | parameter | | | | | | | |
|---|---|---|---|---|---|---|---|---|
| model | intercept ($\beta_0$) | slope ($\beta_1$) | mid-rate ($\alpha$) | location ($\theta$) | rate deviation ($\delta$) | width ($\omega$) | tip error ($\epsilon$) | AICc |
| constant | — | — | -7.00 (-8.57, -5.91) | — | — | — | 0.37 (0.33, 0.42) | 297.24 |
| h. simple | -7.13 (-9.43, -5.73) | -2.88 (-5.29, -0.12) | — | — | — | — | 0.34 (0.29, 0.40) | 291.24 |
| h. threshold | — | — | -4.70 (-5.79, -4.19) | 8.87 (7.91, 9.46) | -1.85 (-7.11, -0.90) | -2.62 (-2.62, 2.16) | 0.34 (0.30, 0.40) | 291.62 |
| h. sweetspot | — | — | -3.85 (-6.81, -1.35) | 8.66 (6.53, 9.05) | 2.23 (0.70, 9.96) | -0.40 (-0.85, 2.15) | 0.33 (0.28, 0.40) | 289.08 |
| d. simple | -14.96 | -4.45 | — | — | — | — | 0.35 | 295.90 |
| d. threshold | — | — | 0.56 | 16.49 | 4.72 | 1.59 | 0.36 | 295.33 |
| d. sweetspot | — | — | -2.59 | 7.31 | 2.77 | -1.85 | 0.33 | ***284.67*** |

Table S16. *Maximum likelihood parameter estimates and associated AICc for models of eucalpyt* ***juvenile leaf length*** *evolution. Here, d. and h. denote models based on simulated dummy factors and maximum height, respectively. Parenthesized numbers represent lower and upper ends of 95% confidence intervals calculated via dentist (Boyko and O'Meara, 2024). All rates are per million years; height/dummy factor measured in millimeters on natural log scale ($\beta_0$ corresponds to rate at mean height/dummy factor value, ~9.6 or 15 meters).*

| | parameter | | | | | | | |
|---|---|---|---|---|---|---|---|---|
| model | intercept ($\beta_0$) | slope ($\beta_1$) | mid-rate ($\alpha$) | location ($\theta$) | rate deviation ($\delta$) | width ($\omega$) | tip error ($\epsilon$) | AICc |
| constant | — | — | -5.58 (-6.42, -4.94) | — | — | — | 0.29 (0.23, 0.34) | 246.19 |
| h. simple | -5.58 (-6.61, -4.86) | -0.03 (-2.22, 1.77) | — | — | — | — | 0.29 (0.22, 0.35) | 248.23 |
| h. threshold | — | — | -3.03 (-7.20, -3.03) | 8.30 (2.23, 9.62) | -1.64 (-1.64, 4.87) | -2.62 (-2.62, 3.60) | 0.27 (0.21, 0.35) | 248.48 |
| h. sweetspot | — | — | -6.03 (-7.40, -4.46) | 9.94 (7.58, 12.05) | -10.00 (-10.00, 9.77) | -0.20 (-0.37, 0.82) | 0.28 (0.21, 0.36) | 251.27 |
| d. simple | -6.26 | -1.16 | — | — | — | — | 0.27 | 234.92 |
| d. threshold | — | — | -2.77 | 11.77 | 1.75 | -1.85 | 0.25 | 232.82 |
| d. sweetspot | — | — | -0.77 | 12.10 | 2.73 | -0.93 | 0.25 | ***230.62*** |

Table S17. *Maximum likelihood parameter estimates and associated AICc for models of eucalpyt* ***juvenile leaf width*** *evolution. Here, d. and h. denote models based on simulated dummy factors and maximum height, respectively. Parenthesized numbers represent lower and upper ends of 95% confidence intervals calculated via dentist (Boyko and O'Meara, 2024). All rates are per million years; height/dummy factor measured in millimeters on natural log scale ($\beta_0$ corresponds to rate at mean height/dummy factor value, ~9.6 or 15 meters).*

| | parameter | | | | | | | |
|---|---|---|---|---|---|---|---|---|
| model | intercept ($\beta_0$) | slope ($\beta_1$) | mid-rate ($\alpha$) | location ($\theta$) | rate deviation ($\delta$) | width ($\omega$) | tip error ($\epsilon$) | AICc |
| constant | — | — | -5.43 (-7.10, -4.37) | — | — | — | 0.61 (0.52, 0.70) | **584.17** |
| h. simple | -5.40 (-7.29, -4.30) | -1.23 (-3.51, 1.44) | — | — | — | — | 0.59 (0.50, 0.70) | ***583.88*** |
| h. threshold | — | — | -5.28 (-7.33, -4.12) | 9.65 (8.82, 9.68) | -10.00 (-10.00, 10.00) | 0.83 (-2.56, 3.59) | 0.59 (0.50, 0.72) | 587.65 |
| h. sweetspot | — | — | -4.15 (-4.18, -4.15) | 10.77 (3.93, 14.67) | -10.00 (-10.00, 9.98) | 2.53 (-1.70, 3.53) | 0.59 (0.48, 0.71) | 587.62 |
| d. simple | -5.43 | 0.07 | — | — | — | — | 0.61 | 586.21 |
| d. threshold | — | — | -5.07 | 10.89 | 0.40 | -1.85 | 0.60 | 590.18 |
| d. sweetspot | — | — | -1.68 | 10.89 | 2.57 | -1.85 | 0.57 | **584.26** |

Table S18. *Maximum likelihood parameter estimates and associated AICc for models of eucalpyt* ***adult leaf length*** *evolution. Here, d. and h. denote models based on simulated dummy factors and maximum height, respectively. Parenthesized numbers represent lower and upper ends of 95% confidence intervals calculated via dentist (Boyko and O'Meara, 2024). All rates are per million years; height/dummy factor measured in millimeters on natural log scale ($\beta_0$ corresponds to rate at mean height/dummy factor value, ~9.6 or 15 meters).*

| | parameter | | | | | | | |
|---|---|---|---|---|---|---|---|---|
| model | intercept ($\beta_0$) | slope ($\beta_1$) | mid-rate ($\alpha$) | location ($\theta$) | rate deviation ($\delta$) | width ($\omega$) | tip error ($\epsilon$) | AICc |
| constant | — | — | -6.41 (-7.26, -5.65) | — | — | — | 0.25 (0.21, 0.29) | 122.15 |
| h. simple | -6.71 (-8.05, -5.66) | -2.15 (-3.98, 0.49) | — | — | — | — | 0.23 (0.19, 0.28) | 118.65 |
| h. threshold | — | — | 7.05 (1.71, 11.93) | 4.10 (2.23, 7.64) | -7.53 (-9.81, -4.53) | 1.36 (-2.61, 1.70) | 0.22 (0.17, 0.28) | 121.54 |
| h. sweetspot | — | — | -3.14 (-4.94, -2.48) | 9.40 (9.21, 9.43) | 2.13 (1.45, 3.58) | -2.62 (-2.62, -0.57) | 0.23 (0.17, 0.28) | 120.36 |
| d. simple | -5.68 | -1.02 | — | — | — | — | 0.17 | 102.57 |
| d. threshold | — | — | -2.02 | 11.95 | 2.38 | -1.85 | 0.20 | 91.58 |
| d. sweetspot | — | — | -1.05 | 12.00 | 2.87 | -1.85 | 0.20 | ***87.40*** |

Table S19. *Maximum likelihood parameter estimates and associated AICc for models of eucalpyt* ***adult leaf width*** *evolution. Here, d. and h. denote models based on simulated dummy factors and maximum height, respectively. Parenthesized numbers represent lower and upper ends of 95% confidence intervals calculated via dentist (Boyko and O'Meara, 2024). All rates are per million years; height/dummy factor measured in millimeters on natural log scale ($\beta_0$ corresponds to rate at mean height/dummy factor value, ~9.6 or 15 meters).*

| | parameter | | | | | | | |
|---|---|---|---|---|---|---|---|---|
| model | intercept ($\beta_0$) | slope ($\beta_1$) | mid-rate ($\alpha$) | location ($\theta$) | rate deviation ($\delta$) | width ($\omega$) | tip error ($\epsilon$) | AICc |
| constant | — | — | -5.70 (-6.79, -4.90) | — | — | — | 0.41 (0.35, 0.47) | 401.85 |
| h. simple | -6.02 (-7.93, -4.94) | -1.52 (-3.89, 0.57) | — | — | — | — | 0.41 (0.34, 0.47) | ***399.65*** |
| h. threshold | — | — | -5.79 (-7.40, -4.96) | 9.72 (9.50, 10.01) | -10.00 (-10.00, 0.46) | -2.36 (-2.61, 0.66) | 0.40 (0.33, 0.49) | 401.69 |
| h. sweetspot | — | — | -3.98 (-4.05, -3.70) | 10.44 (9.96, 12.74) | -10.00 (-10.00, 1.84) | 2.67 (-2.41, 3.60) | 0.41 (0.33, 0.47) | 402.98 |
| d. simple | -5.70 | 0.01 | — | — | — | — | 0.41 | 403.89 |
| d. threshold | — | — | -5.50 | 11.03 | 0.25 | -1.85 | 0.41 | 407.89 |
| d. sweetspot | — | — | -4.07 | 9.71 | -10.00 | 1.88 | 0.38 | 403.24 |

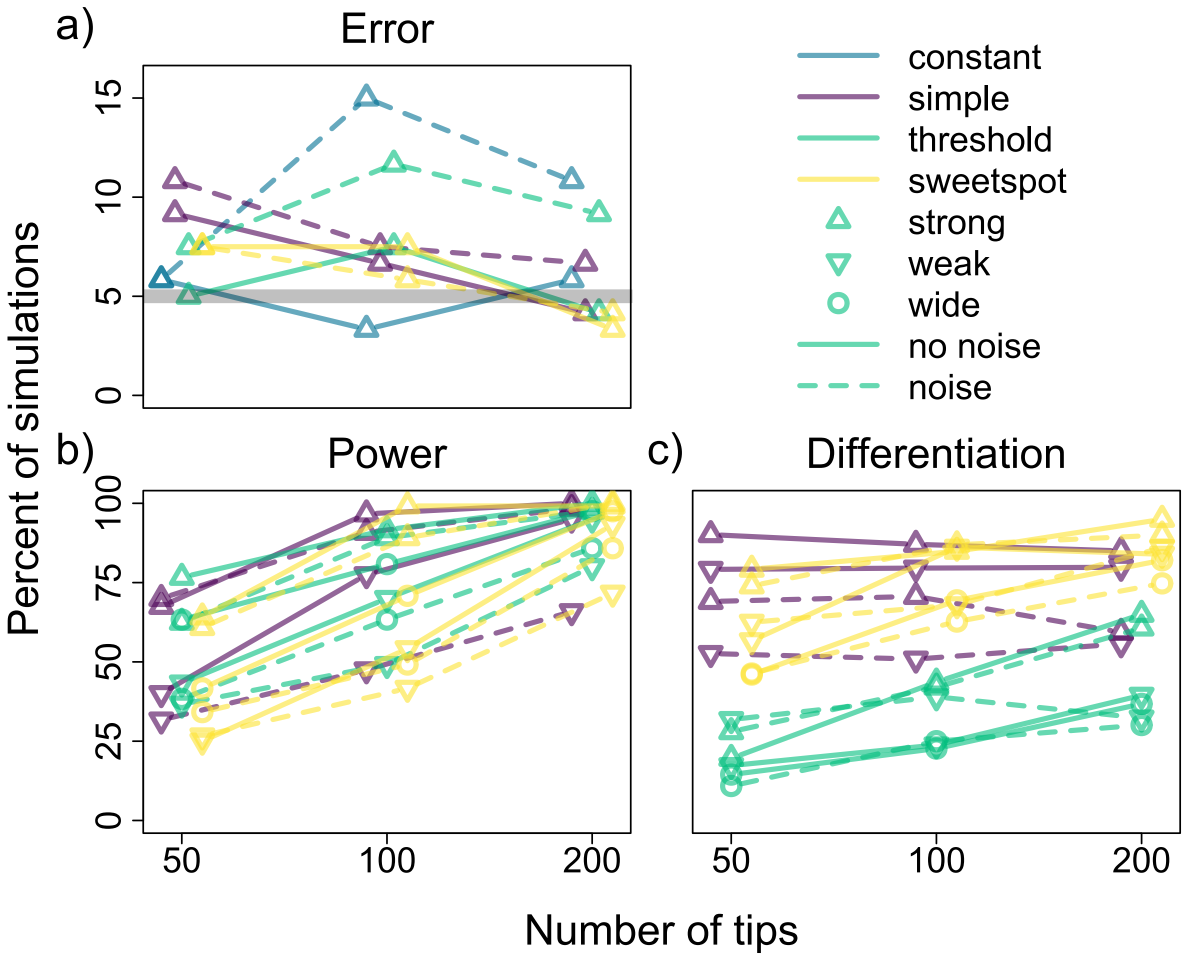

Figure S1. Error, power, and differentiation rates of our continuous stochastic character map-based pipeline for inferring relationships between rates of trait evolution and continuously-varying factors based on a summed AICc weight cutoff of 0.8. Different colors correspond to simulations with different factor-rate relationships (i.e., constant, exponential/simple, logistic/threshold, and Gaussian/sweetspot), different symbols to simulations with differing relationship strength and—in the case of threshold and sweetspot models—width, and dashed versus solid lines to simulations with versus without random variation in rates (noise) around simulated factor-rate relationships. Panel a: percent of simulations with either constant or hidden factor-dependent rates for which the best-fitting model is an observed factor-dependent model (i.e., error rates), with a thick gray line indicating a rate of 5%. Panel b: percent of simulations with observed factor-dependent rates for which the best-fitting model is also an observed factor-dependent model (i.e., power rates). Panel c: percent of observed-factor dependent simulations exhibiting significant evidence for a factor-rate relationship for which the best fitting model assumed the same kind factor-rate relationship used to simulate the data (i.e., differentiation rates).

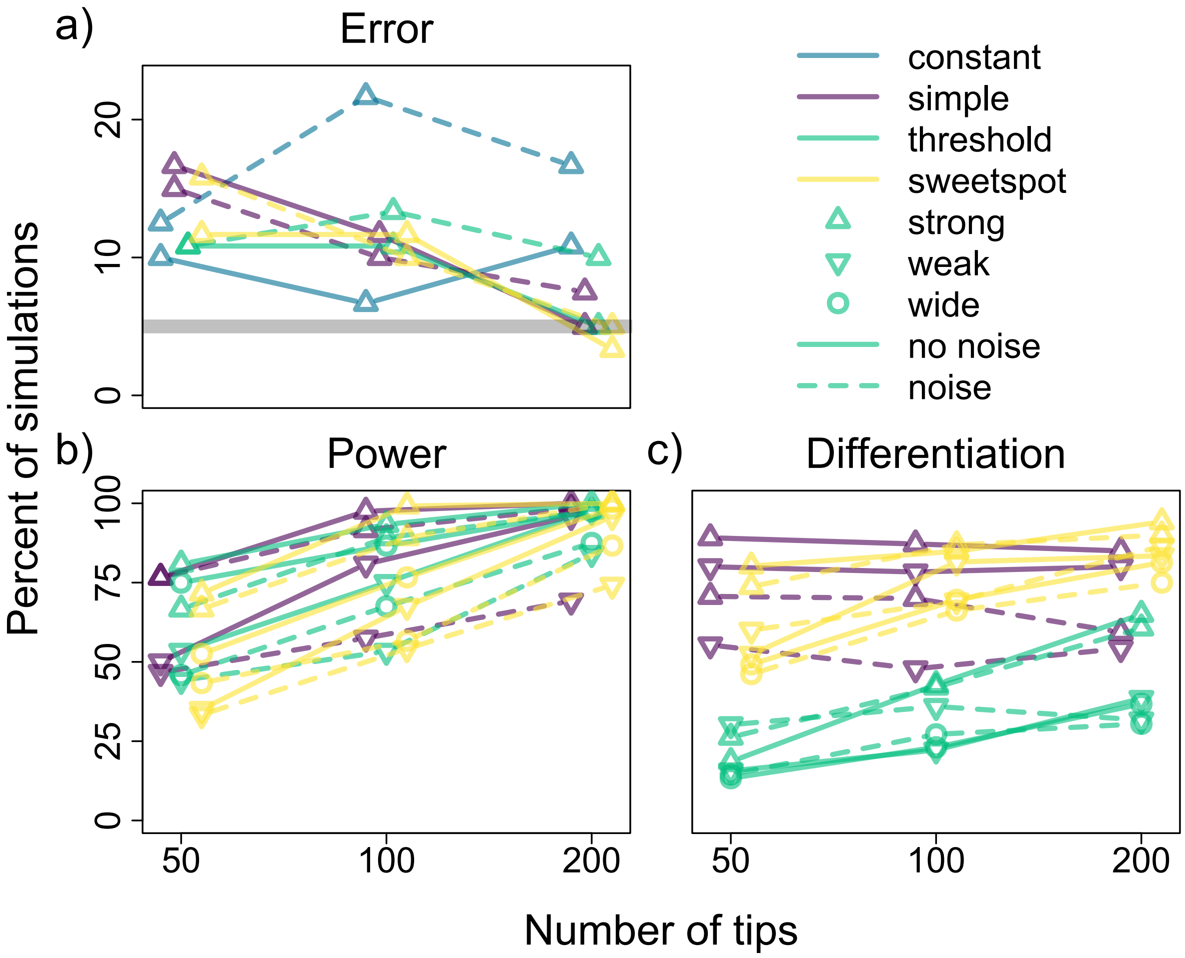

Figure S2. Error, power, and differentiation rates of our continuous stochastic character map-based pipeline for inferring relationships between rates of trait evolution and continuously-varying factors based on a summed AICc weight cutoff of 0.7. Different colors correspond to simulations with different factor-rate relationships (i.e., constant, exponential/simple, logistic/threshold, and Gaussian/sweetspot), different symbols to simulations with differing relationship strength and—in the case of threshold and sweetspot models—width, and dashed versus solid lines to simulations with versus without random variation in rates (noise) around simulated factor-rate relationships. Panel a: percent of simulations with either constant or hidden factor-dependent rates for which the best-fitting model is an observed factor-dependent model (i.e., error rates), with a thick gray line indicating a rate of 5%. Panel b: percent of simulations with observed factor-dependent rates for which the best-fitting model is also an observed factor-dependent model (i.e., power rates). Panel c: percent of observed-factor dependent simulations exhibiting significant evidence for a factor-rate relationship for which the best fitting model assumed the same kind factor-rate relationship used to simulate the data (i.e., differentiation rates).

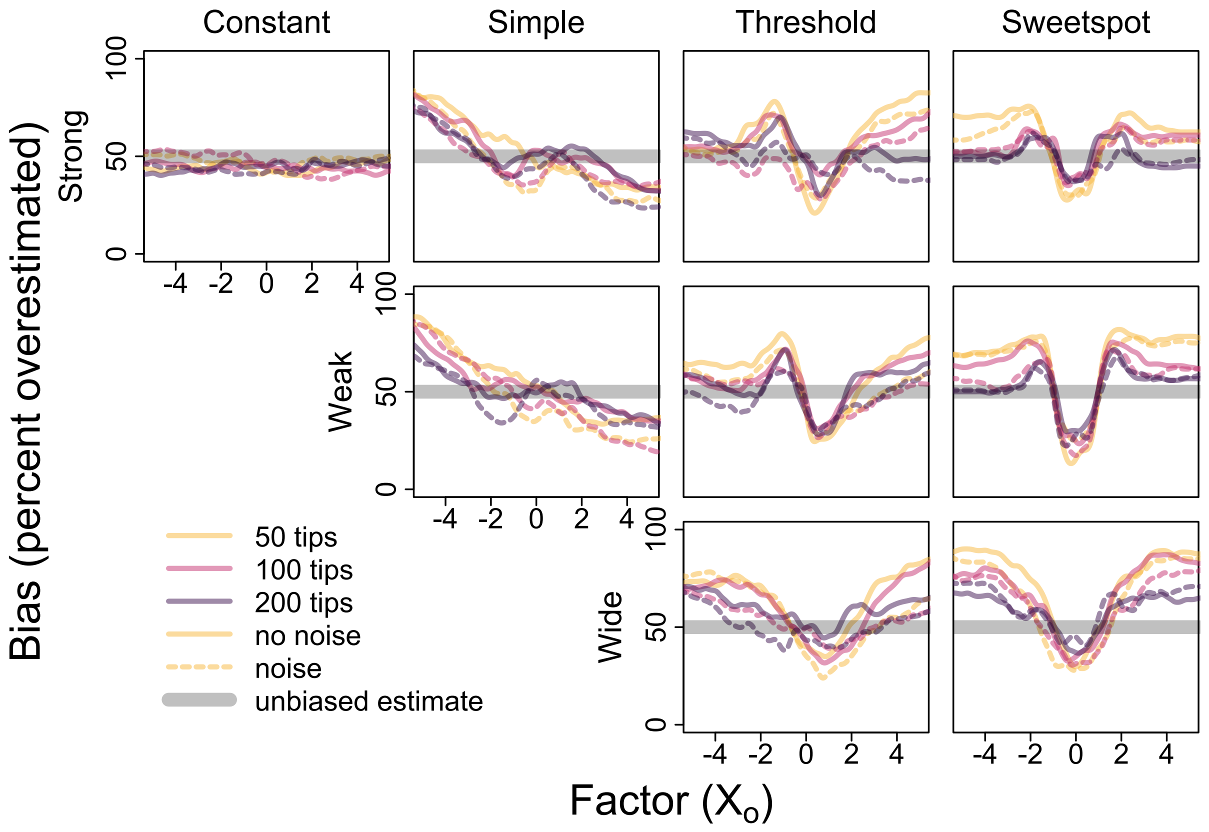

Figure S3. Bias of model-averaged factor-rate relationships (i.e., percent of overestimated rates) estimated using our continuous stochastic character map-based pipeline for inferring relationships between rates of trait evolution and continuously-varying factors for all simulations with either constant or observed factor-dependent rates. Different colors correspond to different sample sizes (i.e., number of tips in simulated phylogeny) and solid versus dashed lines to simulations without versus with random variation in rates (noise) around inferred relationships. Position of unbiased estimation depicted with thick gray line.

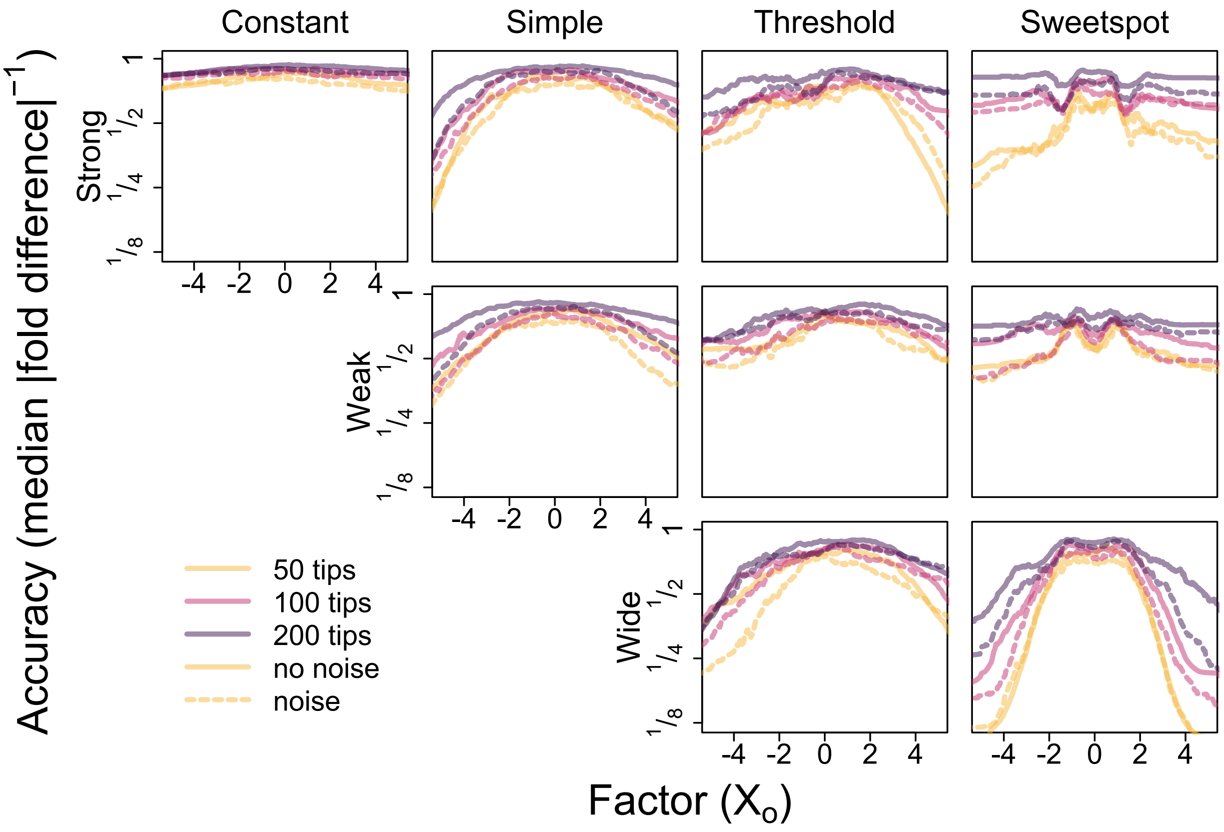

Figure S4. Accuracy of model-averaged factor-rate relationships (i.e., median absolute differences between estimated and simulated rates on log scale, negated such that higher values correspond to greater accuracy) estimated using our continuous stochastic character map-based pipeline for inferring relationships between rates of trait evolution and continuously-varying factors for all simulations with either constant or observed factor-dependent rates. Different colors correspond to different sample sizes (i.e., number of tips in simulated phylogeny) and solid versus dashed lines to simulations without versus with random variation in rates (noise) around inferred relationships.

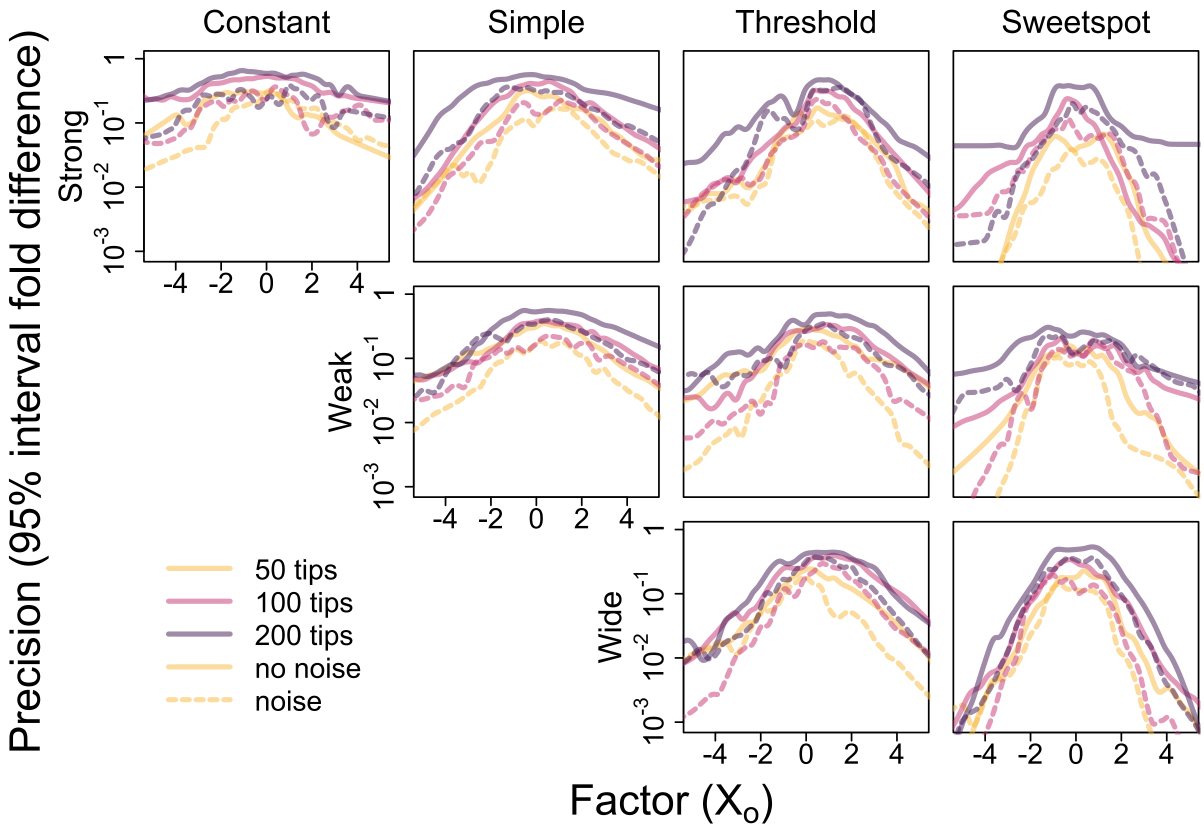

Figure S5. Precision of model-averaged factor-rate relationships (i.e., the 2.5% empirical quantile of estimated rates divided by the corresponding 97.5% empirical quantile) estimated using our continuous stochastic character map-based pipeline for inferring relationships between rates of trait evolution and continuously-varying factors for all simulations with either constant or observed factor-dependent rates. Different colors correspond to different sample sizes (i.e., number of tips in simulated phylogeny) and solid versus dashed lines to simulations without versus with random variation in rates (noise) around inferred relationships.

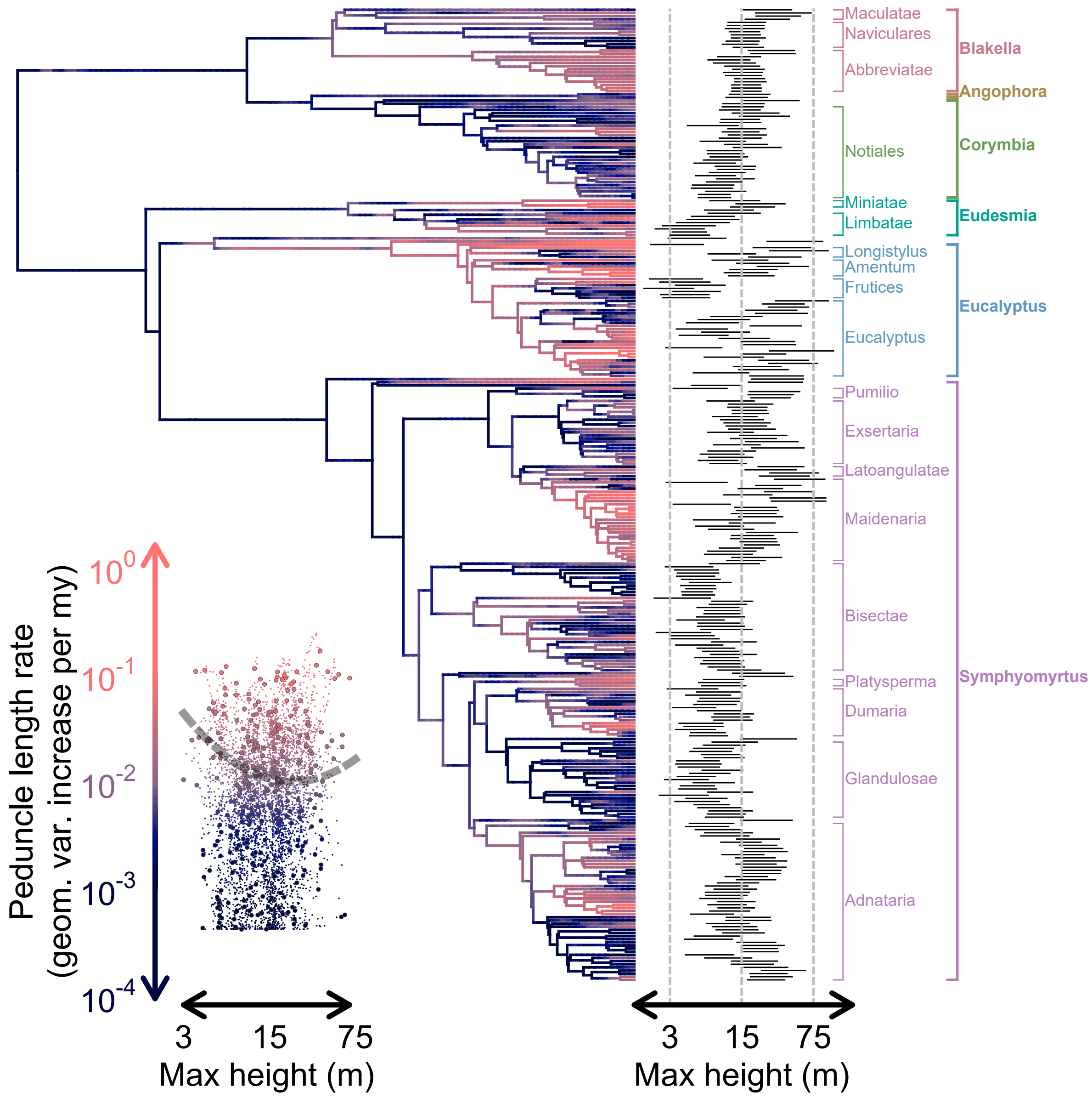

Figure S6. Marginal rate estimates for **peduncle length** evolution mapped onto the eucalypt phylogeny, with dark blue and light red colors indicating low and high rates, respectively (see y-axis of plot on bottom left for color scale; note that the color gradient is standardized across all rate maps for individual traits). Bars depicting 95% credible intervals on maximum height for each tip (inferred under an "evolving rates" or evorates model of height evolution) are arrayed along the right side of the phylogeny, along with clade labels indicating major eucalypt sections and subgenera/genera. The bottom left plot consists of mapped rates for each time point with respect to mean maximum heights (as inferred under the evorates model of height evolution), with larger points indicating tip heights/rates. The gray dashed line running through the plot represents the best-fitting phylogenetic generalized least squares regression relating tip rates to maximum heights (best-fitting based on likelihood ratio tests comparing models assuming either a flat, linear, or quadratic relationship with respect to height).

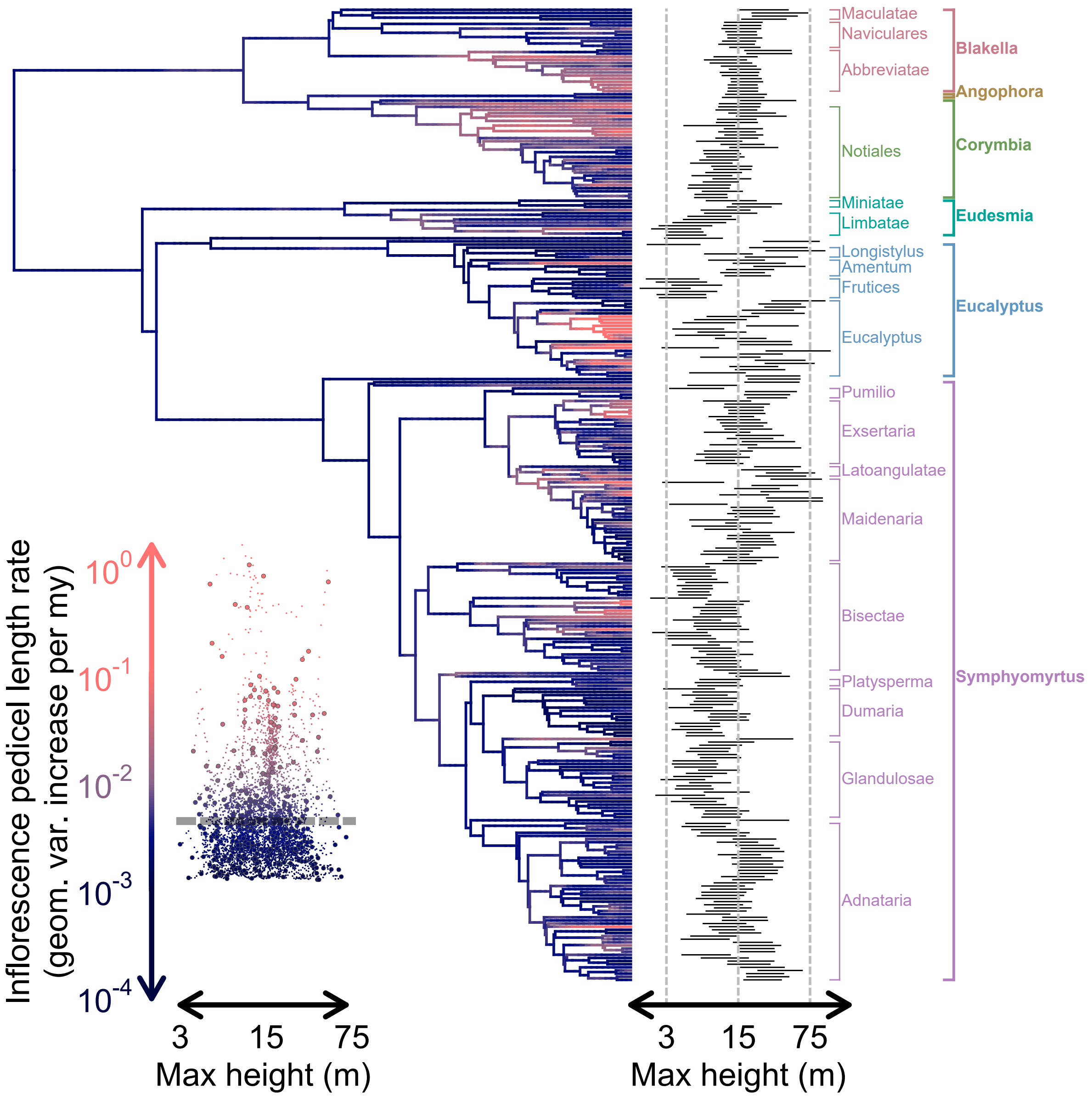

Figure S7. Marginal rate estimates for **inflorescence pedicel length** evolution mapped onto the eucalypt phylogeny, with dark blue and light red colors indicating low and high rates, respectively (see y-axis of plot on bottom left for color scale; note that the color gradient is standardized across all rate maps for individual traits). Bars depicting 95% credible intervals on maximum height for each tip (inferred under an "evolving rates" or evorates model of height evolution) are arrayed along the right side of the phylogeny, along with clade labels indicating major eucalypt sections and subgenera/genera. The bottom left plot consists of mapped rates for each time point with respect to mean maximum heights (as inferred under the evorates model of height evolution), with larger points indicating tip heights/rates. The gray dashed line running through the plot represents the best-fitting phylogenetic generalized least squares regression relating tip rates to maximum heights (best-fitting based on likelihood ratio tests comparing models assuming either a flat, linear, or quadratic relationship with respect to height).

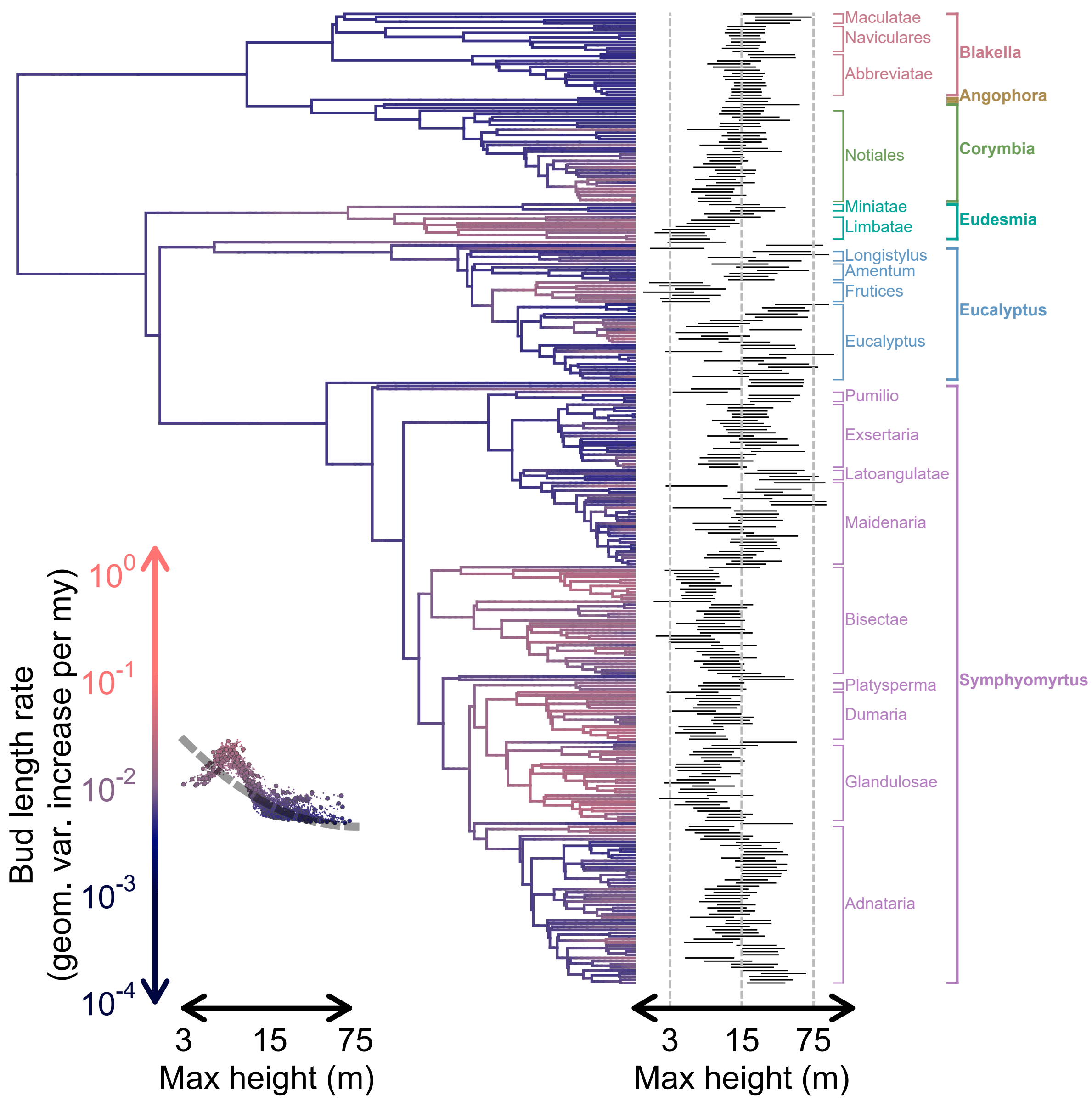

Figure S8. Marginal rate estimates for **bud length** evolution mapped onto the eucalypt phylogeny, with dark blue and light red colors indicating low and high rates, respectively (see y-axis of plot on bottom left for color scale; note that the color gradient is standardized across all rate maps for individual traits). Bars depicting 95% credible intervals on maximum height for each tip (inferred under an "evolving rates" or evorates model of height evolution) are arrayed along the right side of the phylogeny, along with clade labels indicating major eucalypt sections and subgenera/genera. The bottom left plot consists of mapped rates for each time point with respect to mean maximum heights (as inferred under the evorates model of height evolution), with larger points indicating tip heights/rates. The gray dashed line running through the plot represents the best-fitting phylogenetic generalized least squares regression relating tip rates to maximum heights (best-fitting based on likelihood ratio tests comparing models assuming either a flat, linear, or quadratic relationship with respect to height).

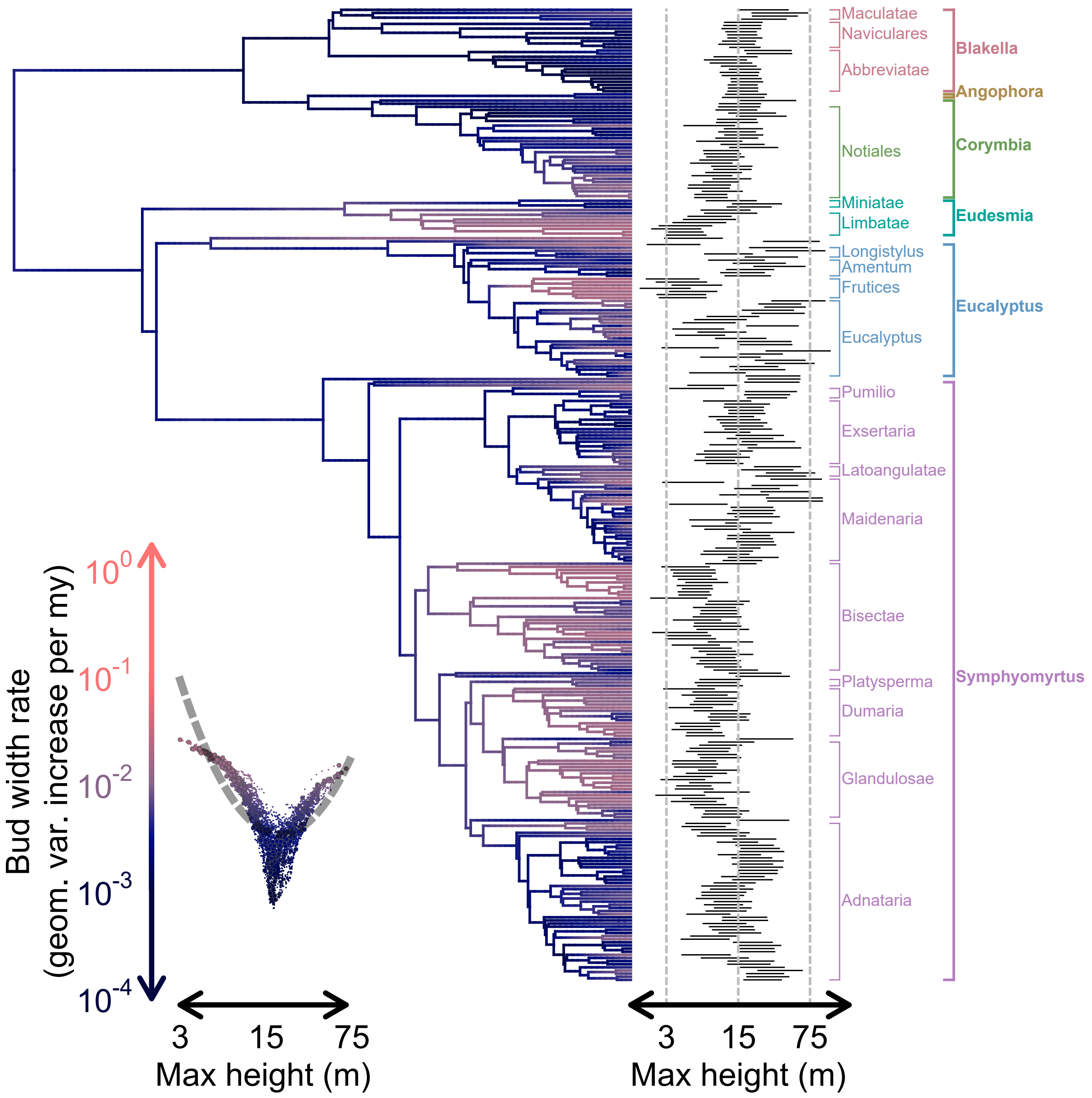

Figure S9. Marginal rate estimates for **bud width** evolution mapped onto the eucalypt phylogeny, with dark blue and light red colors indicating low and high rates, respectively (see y-axis of plot on bottom left for color scale; note that the color gradient is standardized across all rate maps for individual traits). Bars depicting 95% credible intervals on maximum height for each tip (inferred under an "evolving rates" or evorates model of height evolution) are arrayed along the right side of the phylogeny, along with clade labels indicating major eucalypt sections and subgenera/genera. The bottom left plot consists of mapped rates for each time point with respect to mean maximum heights (as inferred under the evorates model of height evolution), with larger points indicating tip heights/rates. The gray dashed line running through the plot represents the best-fitting phylogenetic generalized least squares regression relating tip rates to maximum heights (best-fitting based on likelihood ratio tests comparing models assuming either a flat, linear, or quadratic relationship with respect to height).

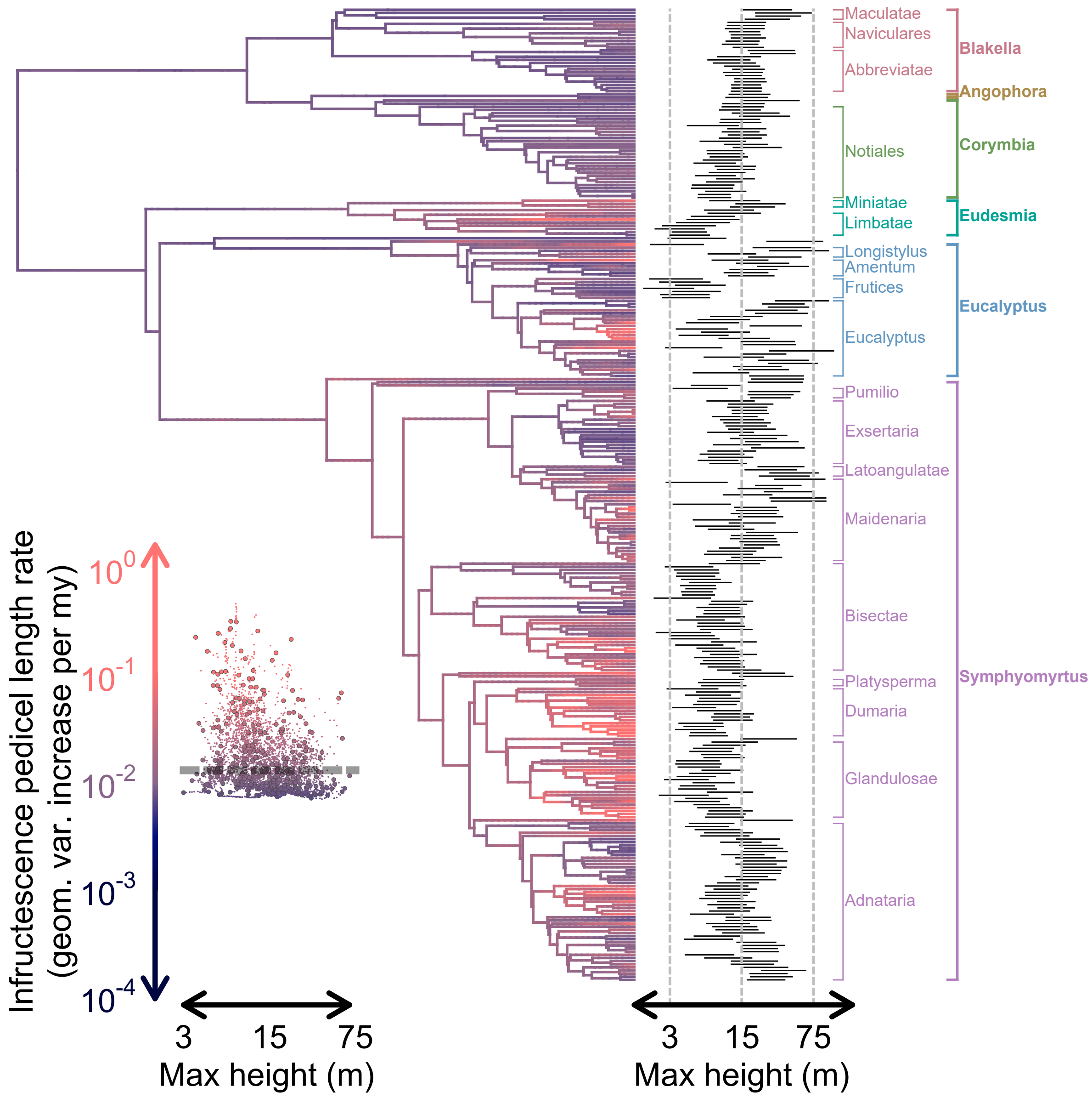

Figure S10. Marginal rate estimates for **infructescence pedicel length** evolution mapped onto the eucalypt phylogeny, with dark blue and light red colors indicating low and high rates, respectively (see y-axis of plot on bottom left for color scale; note that the color gradient is standardized across all rate maps for individual traits). Bars depicting 95% credible intervals on maximum height for each tip (inferred under an "evolving rates" or evorates model of height evolution) are arrayed along the right side of the phylogeny, along with clade labels indicating major eucalypt sections and subgenera/genera. The bottom left plot consists of mapped rates for each time point with respect to mean maximum heights (as inferred under the evorates model of height evolution), with larger points indicating tip heights/rates. The gray dashed line running through the plot represents the best-fitting phylogenetic generalized least squares regression relating tip rates to maximum heights (best-fitting based on likelihood ratio tests comparing models assuming either a flat, linear, or quadratic relationship with respect to height).

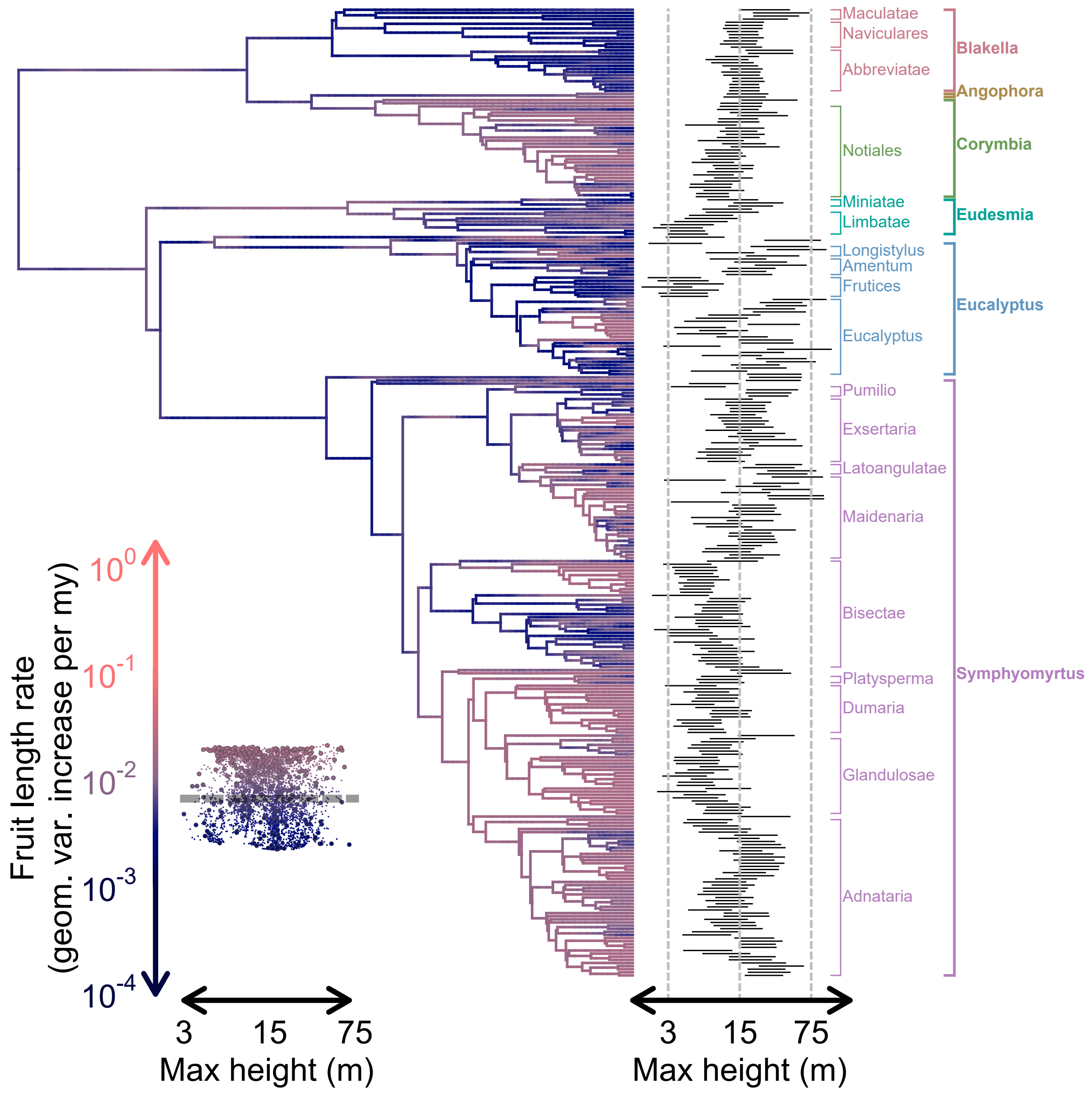

Figure S11. Marginal rate estimates for **fruit length** evolution mapped onto the eucalypt phylogeny, with dark blue and light red colors indicating low and high rates, respectively (see y-axis of plot on bottom left for color scale; note that the color gradient is standardized across all rate maps for individual traits). Bars depicting 95% credible intervals on maximum height for each tip (inferred under an "evolving rates" or evorates model of height evolution) are arrayed along the right side of the phylogeny, along with clade labels indicating major eucalypt sections and subgenera/genera. The bottom left plot consists of mapped rates for each time point with respect to mean maximum heights (as inferred under the evorates model of height evolution), with larger points indicating tip heights/rates. The gray dashed line running through the plot represents the best-fitting phylogenetic generalized least squares regression relating tip rates to maximum heights (best-fitting based on likelihood ratio tests comparing models assuming either a flat, linear, or quadratic relationship with respect to height).

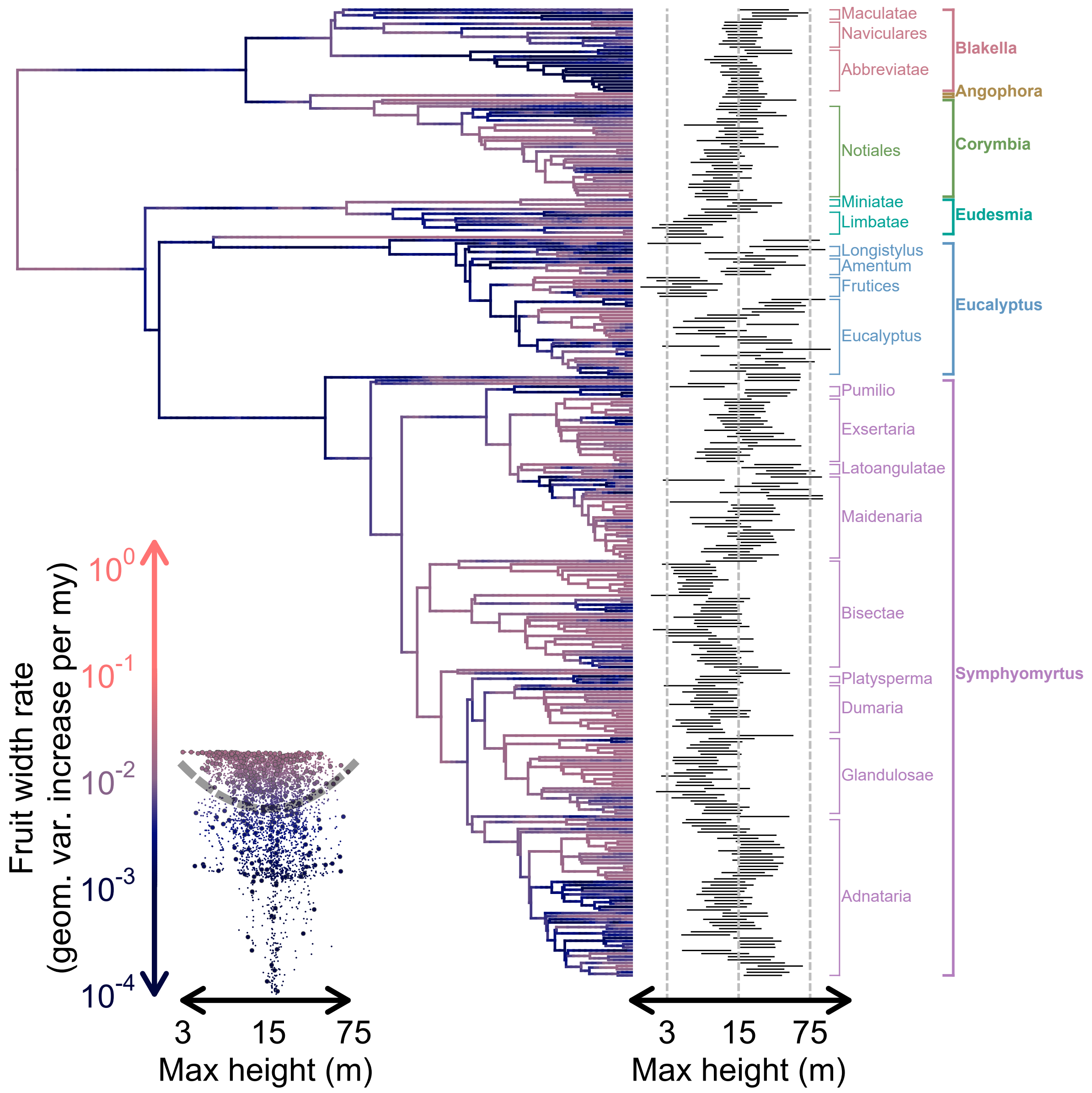

Figure S12. Marginal rate estimates for **fruit width** evolution mapped onto the eucalypt phylogeny, with dark blue and light red colors indicating low and high rates, respectively (see y-axis of plot on bottom left for color scale; note that the color gradient is standardized across all rate maps for individual traits). Bars depicting 95% credible intervals on maximum height for each tip (inferred under an "evolving rates" or evorates model of height evolution) are arrayed along the right side of the phylogeny, along with clade labels indicating major eucalypt sections and subgenera/genera. The bottom left plot consists of mapped rates for each time point with respect to mean maximum heights (as inferred under the evorates model of height evolution), with larger points indicating tip heights/rates. The gray dashed line running through the plot represents the best-fitting phylogenetic generalized least squares regression relating tip rates to maximum heights (best-fitting based on likelihood ratio tests comparing models assuming either a flat, linear, or quadratic relationship with respect to height).

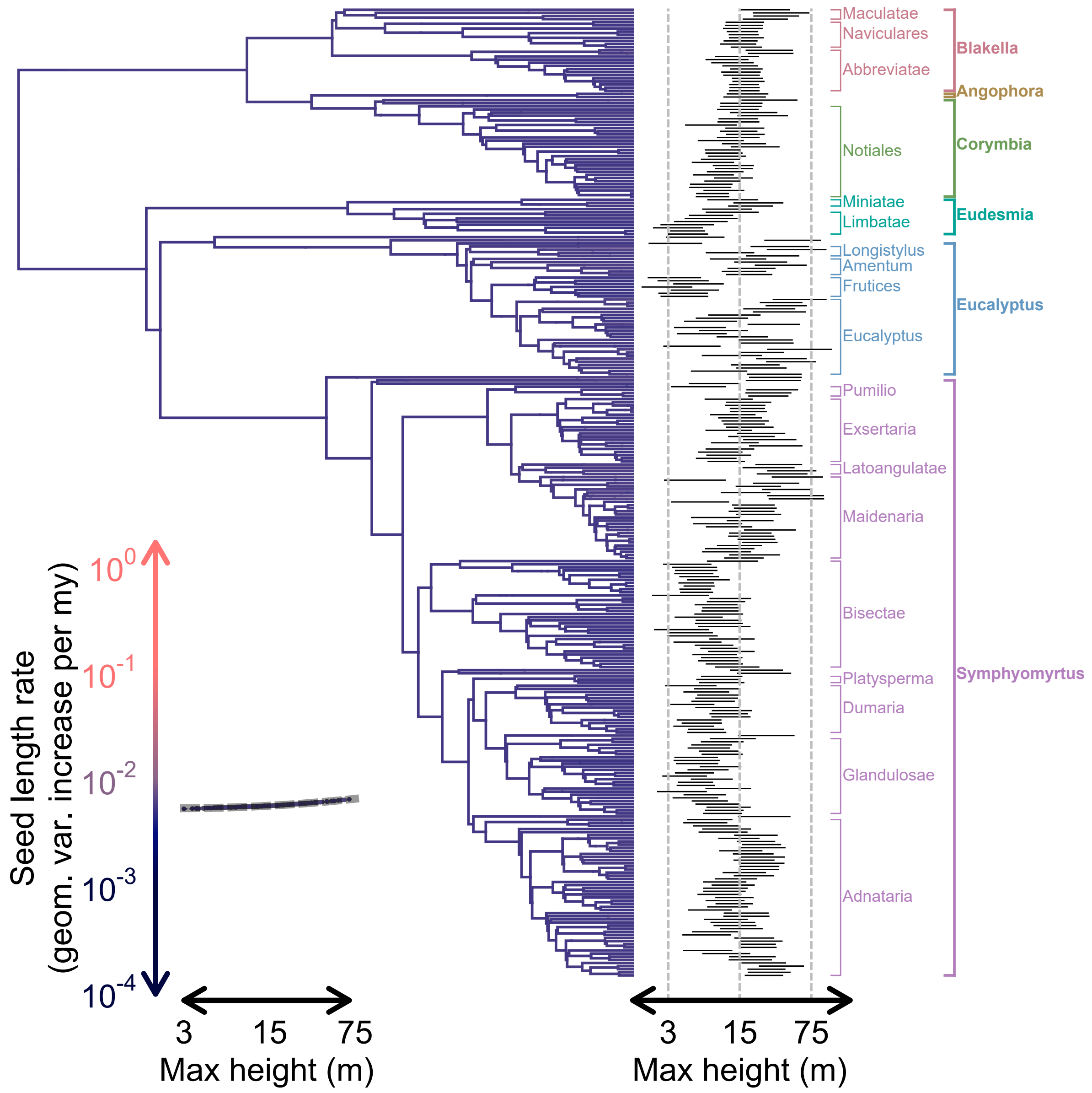

Figure S13. Marginal rate estimates for **seed length** evolution mapped onto the eucalypt phylogeny, with dark blue and light red colors indicating low and high rates, respectively (see y-axis of plot on bottom left for color scale; note that the color gradient is standardized across all rate maps for individual traits). Bars depicting 95% credible intervals on maximum height for each tip (inferred under an "evolving rates" or evorates model of height evolution) are arrayed along the right side of the phylogeny, along with clade labels indicating major eucalypt sections and subgenera/genera. The bottom left plot consists of mapped rates for each time point with respect to mean maximum heights (as inferred under the evorates model of height evolution), with larger points indicating tip heights/rates. The gray dashed line running through the plot represents the best-fitting phylogenetic generalized least squares regression relating tip rates to maximum heights (best-fitting based on likelihood ratio tests comparing models assuming either a flat, linear, or quadratic relationship with respect to height).

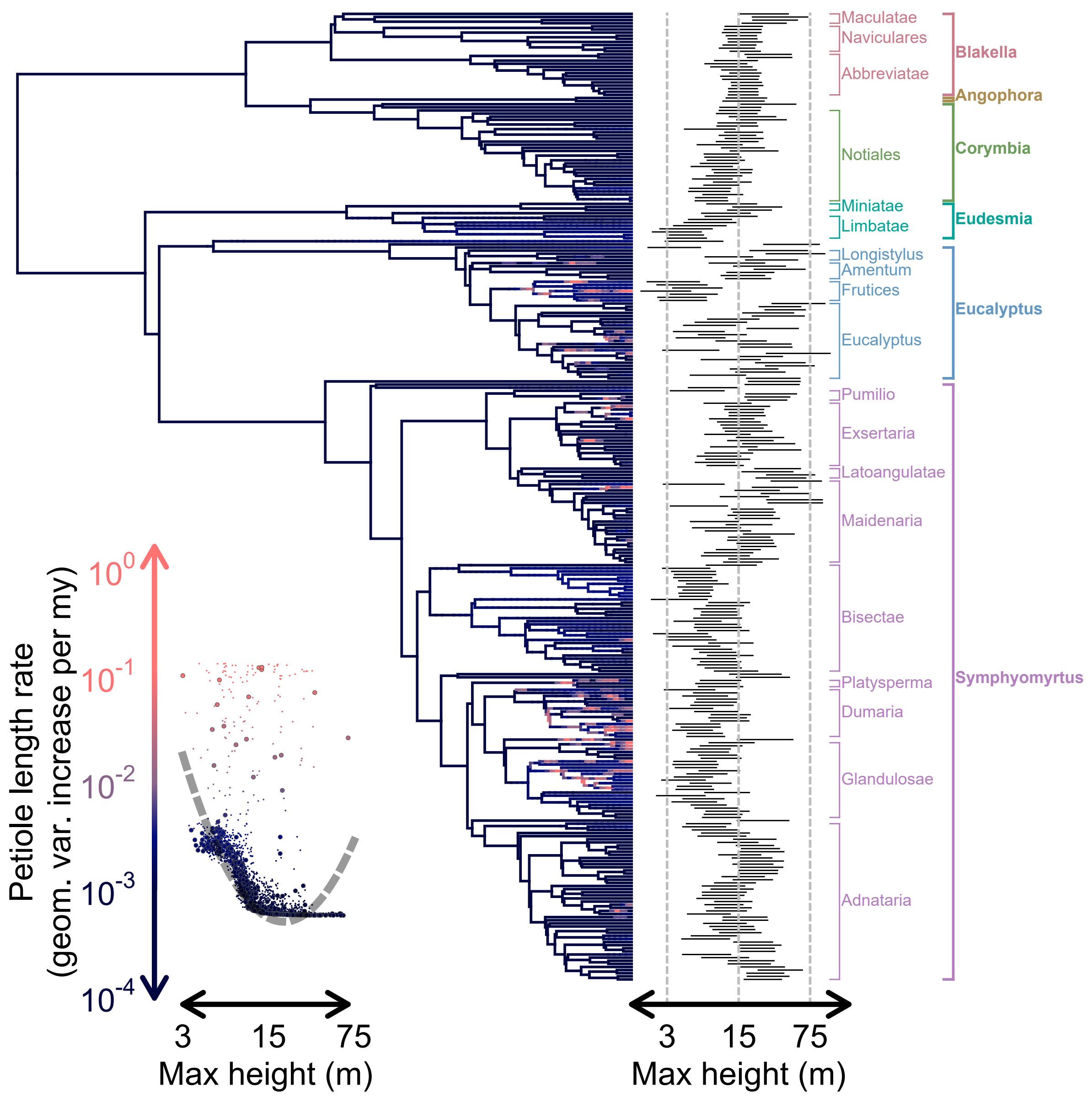

Figure S14. Marginal rate estimates for **petiole length** evolution mapped onto the eucalypt phylogeny, with dark blue and light red colors indicating low and high rates, respectively (see y-axis of plot on bottom left for color scale; note that the color gradient is standardized across all rate maps for individual traits). Bars depicting 95% credible intervals on maximum height for each tip (inferred under an "evolving rates" or evorates model of height evolution) are arrayed along the right side of the phylogeny, along with clade labels indicating major eucalypt sections and subgenera/genera. The bottom left plot consists of mapped rates for each time point with respect to mean maximum heights (as inferred under the evorates model of height evolution), with larger points indicating tip heights/rates. The gray dashed line running through the plot represents the best-fitting phylogenetic generalized least squares regression relating tip rates to maximum heights (best-fitting based on likelihood ratio tests comparing models assuming either a flat, linear, or quadratic relationship with respect to height).

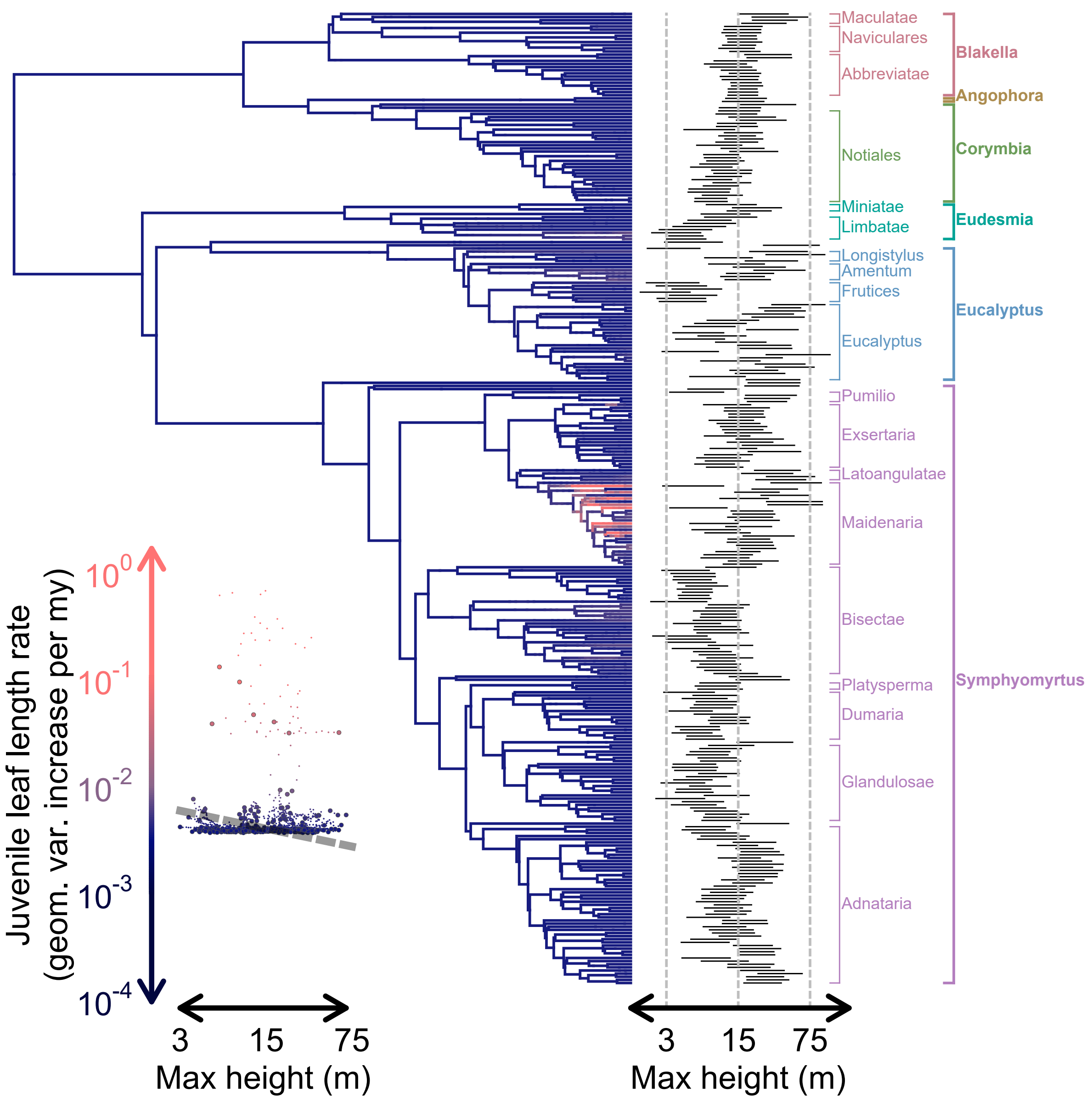

Figure S15. Marginal rate estimates for **juvenile leaf length** evolution mapped onto the eucalypt phylogeny, with dark blue and light red colors indicating low and high rates, respectively (see y-axis of plot on bottom left for color scale; note that the color gradient is standardized across all rate maps for individual traits). Bars depicting 95% credible intervals on maximum height for each tip (inferred under an "evolving rates" or evorates model of height evolution) are arrayed along the right side of the phylogeny, along with clade labels indicating major eucalypt sections and subgenera/genera. The bottom left plot consists of mapped rates for each time point with respect to mean maximum heights (as inferred under the evorates model of height evolution), with larger points indicating tip heights/rates. The gray dashed line running through the plot represents the best-fitting phylogenetic generalized least squares regression relating tip rates to maximum heights (best-fitting based on likelihood ratio tests comparing models assuming either a flat, linear, or quadratic relationship with respect to height).

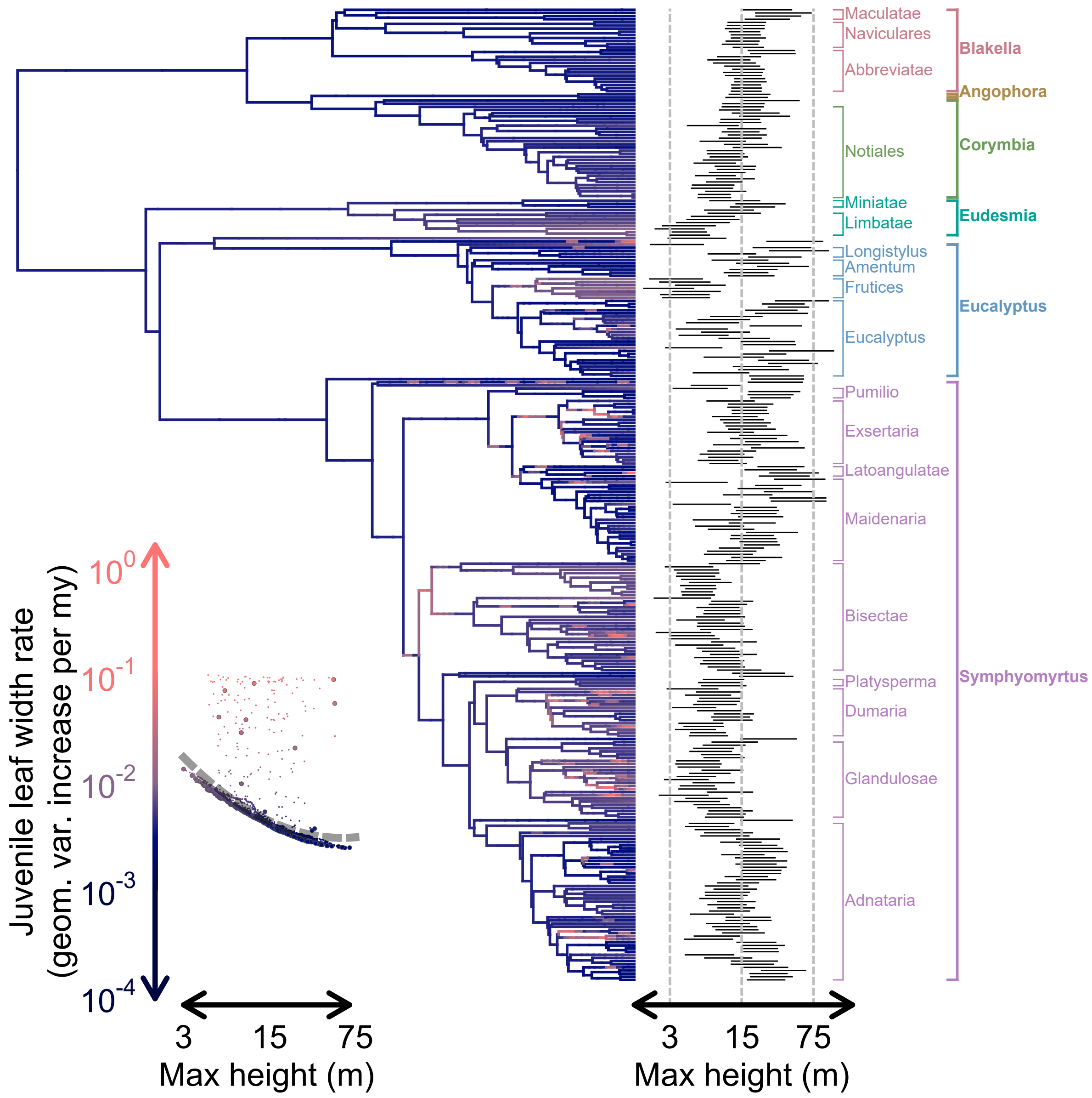

Figure S16. Marginal rate estimates for **juvenile leaf width** evolution mapped onto the eucalypt phylogeny, with dark blue and light red colors indicating low and high rates, respectively (see y-axis of plot on bottom left for color scale; note that the color gradient is standardized across all rate maps for individual traits). Bars depicting 95% credible intervals on maximum height for each tip (inferred under an "evolving rates" or evorates model of height evolution) are arrayed along the right side of the phylogeny, along with clade labels indicating major eucalypt sections and subgenera/genera. The bottom left plot consists of mapped rates for each time point with respect to mean maximum heights (as inferred under the evorates model of height evolution), with larger points indicating tip heights/rates. The gray dashed line running through the plot represents the best-fitting phylogenetic generalized least squares regression relating tip rates to maximum heights (best-fitting based on likelihood ratio tests comparing models assuming either a flat, linear, or quadratic relationship with respect to height).

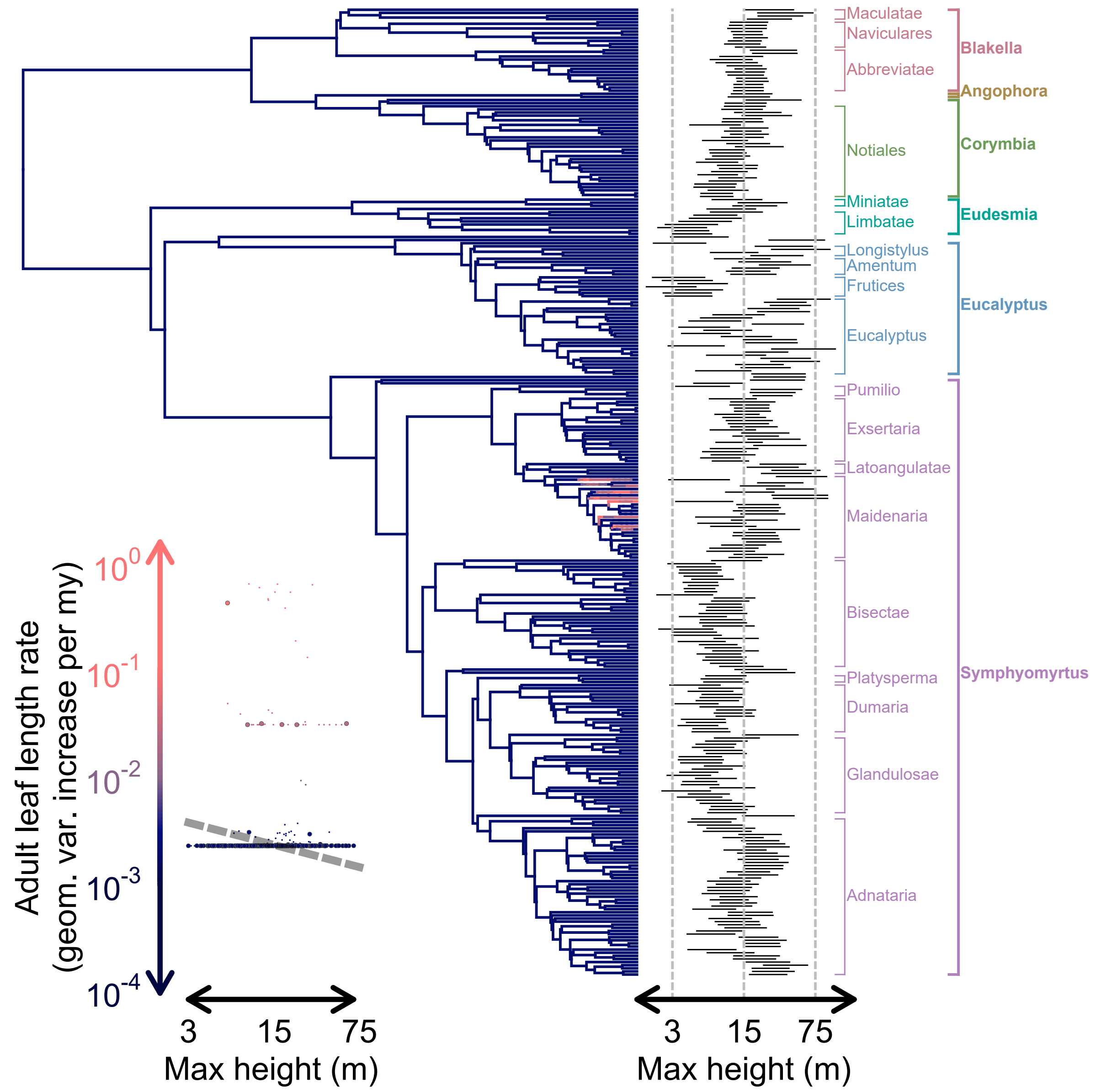

Figure S17. Marginal rate estimates for **adult leaf length** evolution mapped onto the eucalypt phylogeny, with dark blue and light red colors indicating low and high rates, respectively (see y-axis of plot on bottom left for color scale; note that the color gradient is standardized across all rate maps for individual traits). Bars depicting 95% credible intervals on maximum height for each tip (inferred under an "evolving rates" or evorates model of height evolution) are arrayed along the right side of the phylogeny, along with clade labels indicating major eucalypt sections and subgenera/genera. The bottom left plot consists of mapped rates for each time point with respect to mean maximum heights (as inferred under the evorates model of height evolution), with larger points indicating tip heights/rates. The gray dashed line running through the plot represents the best-fitting phylogenetic generalized least squares regression relating tip rates to maximum heights (best-fitting based on likelihood ratio tests comparing models assuming either a flat, linear, or quadratic relationship with respect to height).

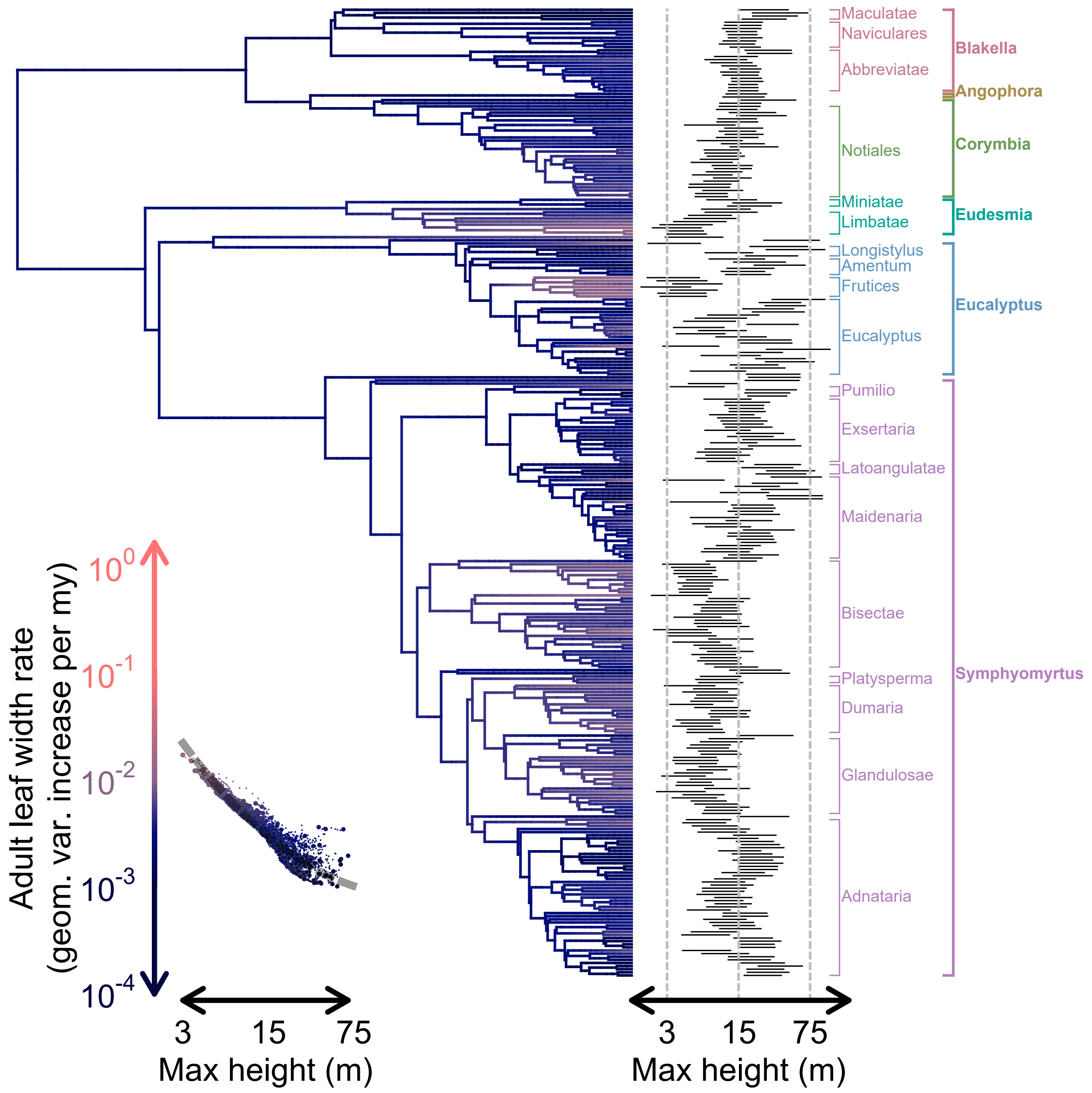

Figure S18. Marginal rate estimates for **adult leaf width** evolution mapped onto the eucalypt phylogeny, with dark blue and light red colors indicating low and high rates, respectively (see y-axis of plot on bottom left for color scale; note that the color gradient is standardized across all rate maps for individual traits). Bars depicting 95% credible intervals on maximum height for each tip (inferred under an "evolving rates" or evorates model of height evolution) are arrayed along the right side of the phylogeny, along with clade labels indicating major eucalypt sections and subgenera/genera. The bottom left plot consists of mapped rates for each time point with respect to mean maximum heights (as inferred under the evorates model of height evolution), with larger points indicating tip heights/rates. The gray dashed line running through the plot represents the best-fitting phylogenetic generalized least squares regression relating tip rates to maximum heights (best-fitting based on likelihood ratio tests comparing models assuming either a flat, linear, or quadratic relationship with respect to height).

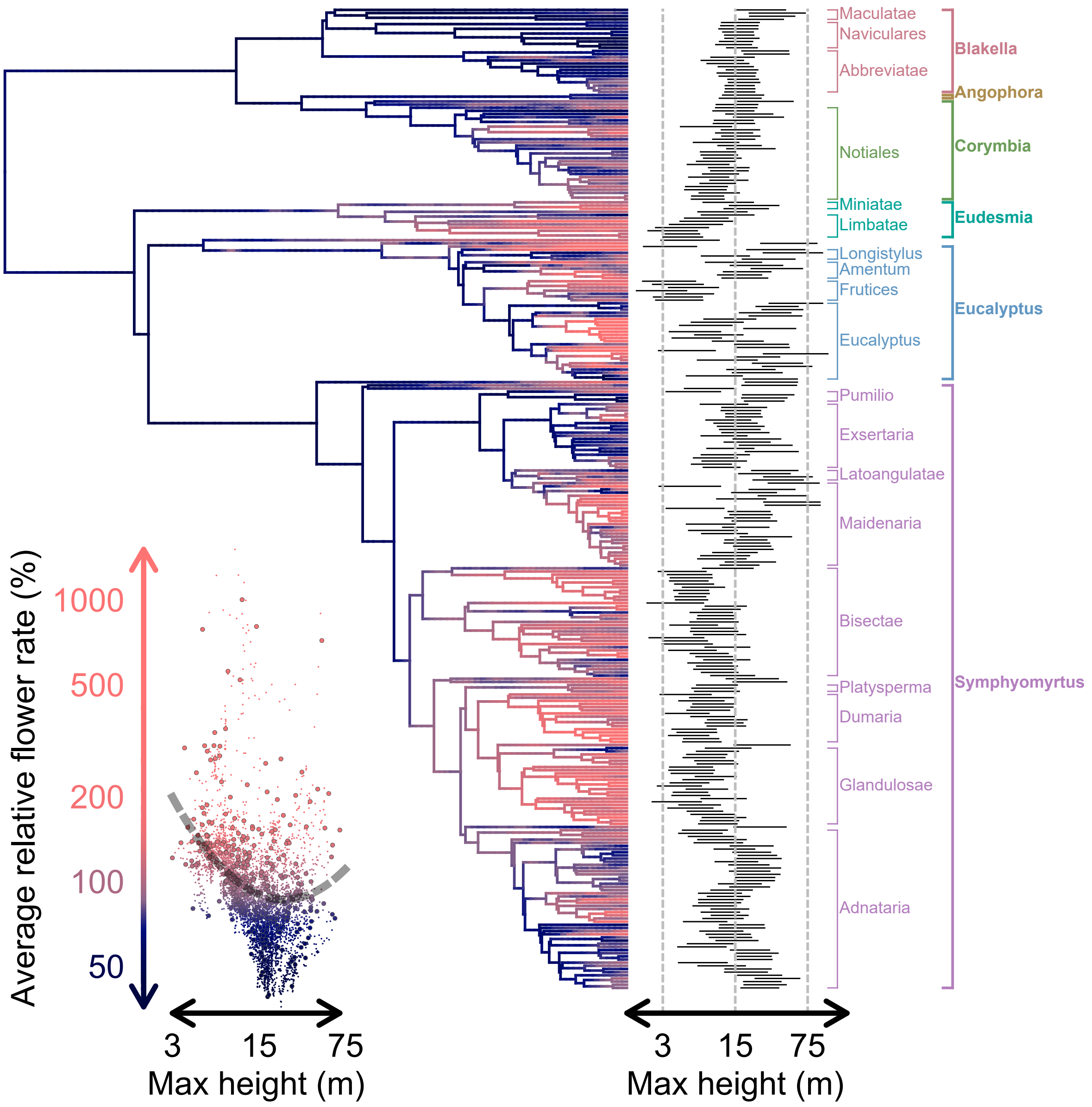

Figure S19. Relative rates of phenotypic evolution, averaged over all **flower traits**, mapped onto the eucalypt phylogeny, with dark blue and light red colors indicating relatively low and high rates, respectively (see y-axis of plot on bottom left for color scale; note that the color gradient is percentile-based due to the right-skewed distribution of relative rates on a log scale). Bars depicting 95% credible intervals on maximum height for each tip (inferred under an "evolving rates" or evorates model of height evolution) are arrayed along the right side of the phylogeny, along with clade labels indicating major eucalypt sections and subgenera/genera. The bottom left plot consists of mapped rates for each time point with respect to mean maximum heights (as inferred under the evorates model of height evolution), with larger points indicating tip heights/rates. The gray dashed line running through the plot represents the best-fitting phylogenetic generalized least squares regression relating tip rates to maximum heights (best-fitting based on likelihood ratio tests comparing models assuming either a flat, linear, or quadratic relationship with respect to height).

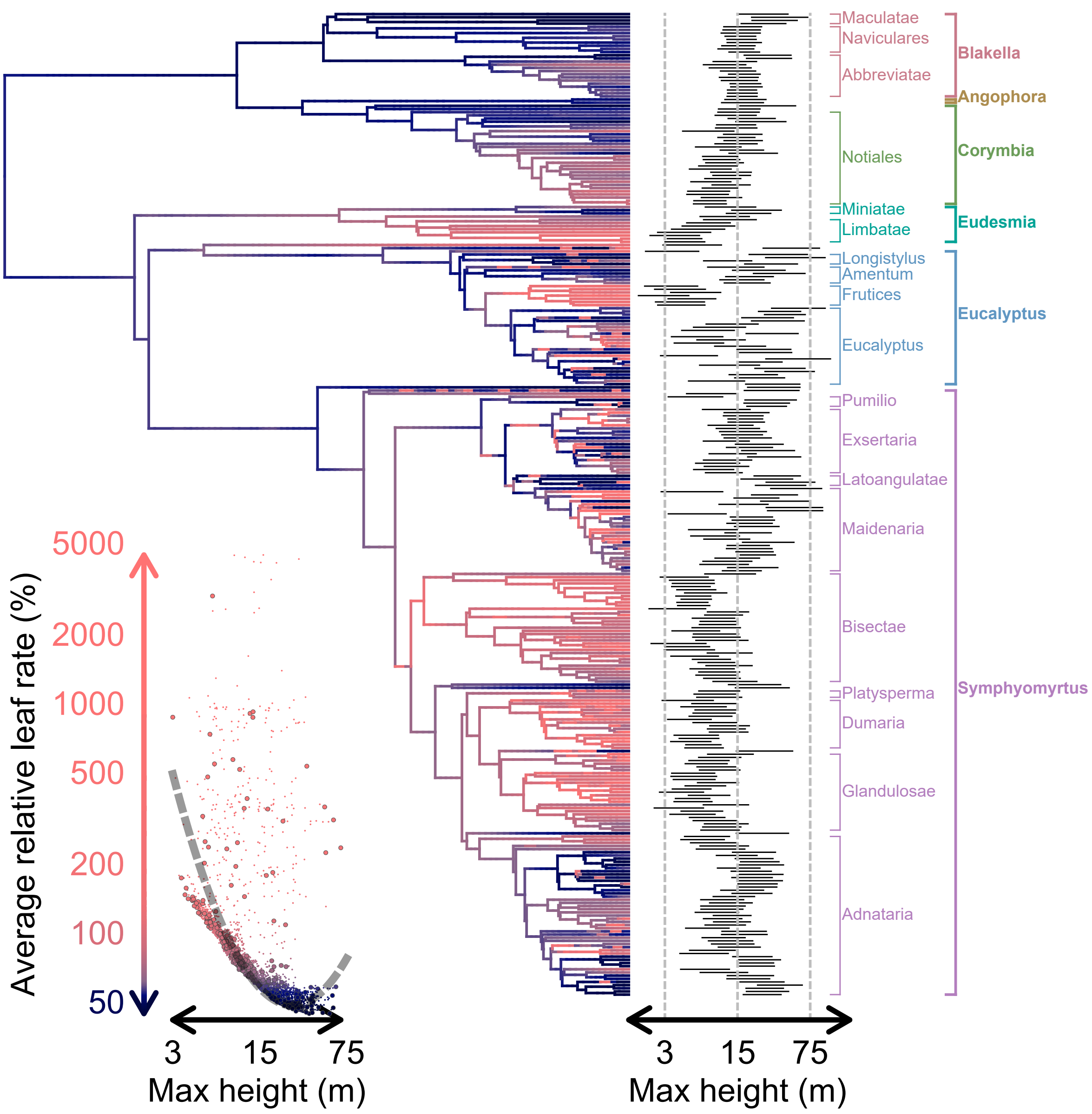

Figure S20. Relative rates of phenotypic evolution, averaged over all **leaf traits**, mapped onto the eucalypt phylogeny, with dark blue and light red colors indicating relatively low and high rates, respectively (see y-axis of plot on bottom left for color scale; note that the color gradient is percentile-based due to the right-skewed distribution of relative rates on a log scale). Bars depicting 95% credible intervals on maximum height for each tip (inferred under an "evolving rates" or evorates model of height evolution) are arrayed along the right side of the phylogeny, along with clade labels indicating major eucalypt sections and subgenera/genera. The bottom left plot consists of mapped rates for each time point with respect to mean maximum heights (as inferred under the evorates model of height evolution), with larger points indicating tip heights/rates. The gray dashed line running through the plot represents the best-fitting phylogenetic generalized least squares regression relating tip rates to maximum heights (best-fitting based on likelihood ratio tests comparing models assuming either a flat, linear, or quadratic relationship with respect to height).

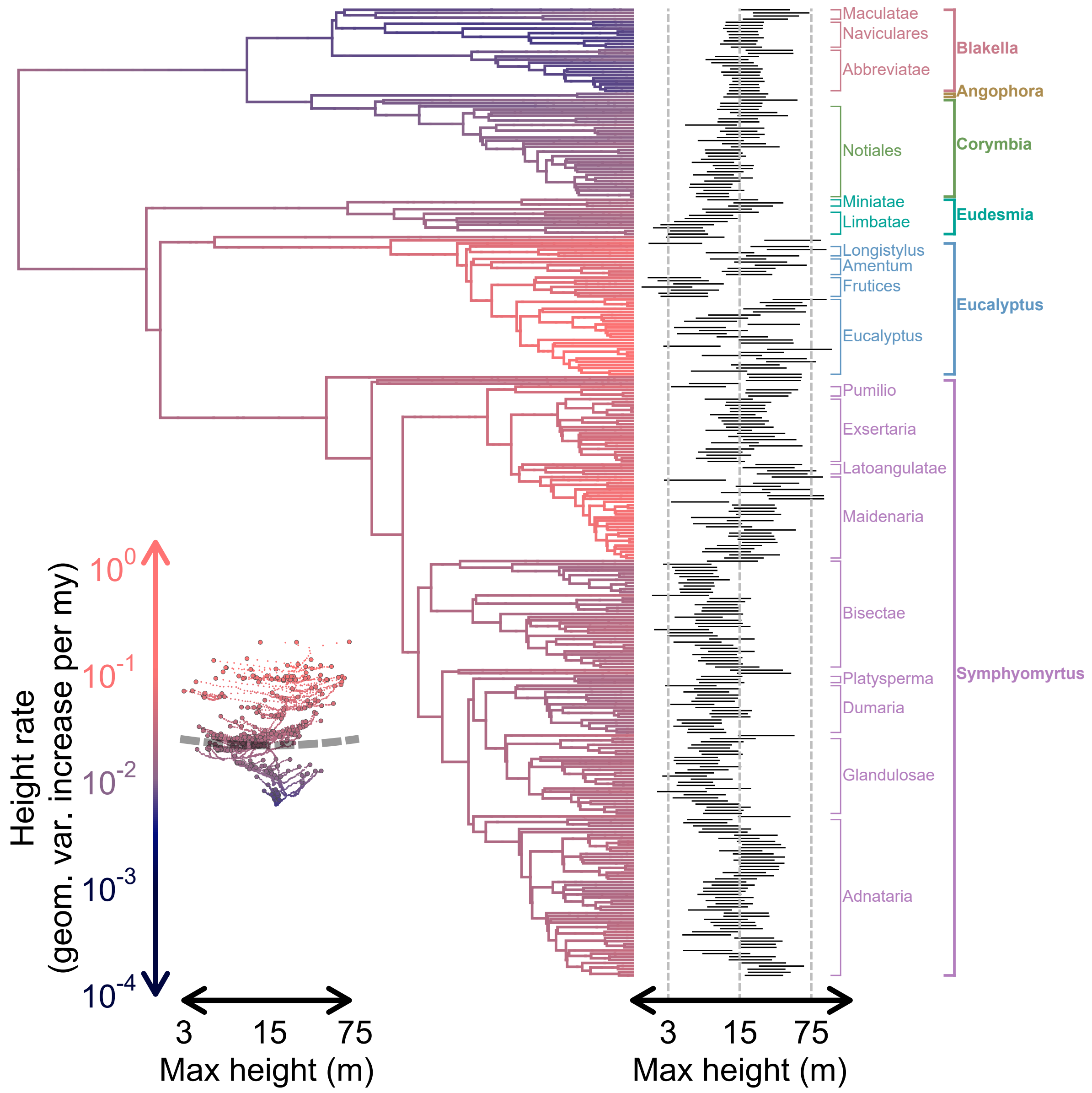

Figure S21. Marginal rate estimates for **maximum height** evolution mapped onto the eucalypt phylogeny, with dark blue and light red colors indicating low and high rates, respectively (see y-axis of plot on bottom left for color scale; note that the color gradient is standardized across all rate maps for individual traits). Note that these rates are *not* estimated from continuous stochastic character map-based models as they are for other individual traits—instead, they are based on continuous stochastic character maps under an "evolving rates" or evorates model of height evolution. Bars depicting 95% credible intervals on maximum height for each tip inferred under the very same model are arrayed along the right side of the phylogeny, along with clade labels indicating major eucalypt sections and subgenera/genera. The bottom left plot consists of mapped rates for each time point with respect to mean maximum heights, with larger points indicating tip heights/rates. The gray dashed line running through the plot represents the best-fitting phylogenetic generalized least squares regression relating tip rates to maximum heights (best-fitting based on likelihood ratio tests comparing models assuming either a flat, linear, or quadratic relationship with respect to height). Note that there is no substantial evidence for any kind of "autocorrelated rate variation" (e.g., rates of height evolution increasing among larger-bodied lineages).

### Technical Details of Continuous Stochastic Character Mapping Algorithm

Our continuous stochastic character mapping algorithm is primarily based on two observations. First, phenotypic changes under Brownian Motion models are normally distributed and independent among distinct lineages (i.e., non-overlapping portions of a phylogeny such as different edges; Karatzas and Shreve, 2004). Second, given multiple independent measurements of some phenotypic value with measurement errors, the distribution of the phenotypic value itself is given by the normalized product of the probability density functions associated with each measurement. This second observation is particularly convenient in the case of normally distributed measurement errors, as the product of multiple normal probability density functions is proportional to another normal probability density function (even in the case of multivariate normal distributions; see Petersen and Pedersen, 2012). Together, these two observations imply that distributions of phenotypic values evolving under Brownian Motion are normal at any point on a phylogeny—even if conditioning on observed measurements (at least assuming all measurement errors are normally distributed). Further, the mean and variance of a phenotypic distribution at any given focal time point on a phylogeny may be calculated by treating phenotypic distributions at all directly adjacent time points as independent "measurements" of the phenotype at the focal point. Note that it suffices to only keep track of the mean and variance of each phenotypic distribution because normal distributions are completely defined by these two parameters (Petersen and Pedersen, 2012).

Given the above observations, the phenotypic distributions at all nodes in a phylogeny—along with distributions at arbitrary time points along each edge—can be efficiently computed and sampled from by visiting each time point on the phylogeny twice. Once for each node during a "backwards" or "postorder" traversal from tips to root, and again for each node—as well as time points along each edge—during the "forwards" or "preorder" traversal from root to tips (this general approach is used by many other rapid ancestral state reconstruction algorithms, e.g.,Yang, 2006; Hiscott et al., 2016; Goolsby, 2017; Hassler et al., 2022).

*The simplest case: exact univariate measurements at tips*

We start off by describing the detailed steps of our algorithm under a relatively simple scenario: a phylogenetic comparative dataset on a single continuous phenotype with one exact measurement per tip, assuming a homogeneous Brownian Motion model with constant evolutionary rates/trends and no intraspecifc variation/measurement error (i.e., exact measurements). Let $y$ denote an $n$-length vector of observed phenotypic measurements, where $n$ represents the number of tips on a rooted phylogeny consisting of $e$ edges with branch lengths in units of time (given by an $e$-length vector $l$). Further, let $x$ and $v$ be $e$-length vectors of the (currently unknown) means and variances, respectively, of the phenotypic distributions at each node of the phylogeny, with the $i$th entries of $x/v$ corresponding to the node immediately descending from edge $i$ (note that this notation assumes the phylogeny has a "stem" edge ancestral to the root). Overall, our algorithm aims to recursively compute $x/v$ and sample values from these distributions (plus additional values along each edge of the phylogeny) given the measurements $y$ along with an estimated evolutionary rate $\sigma^2$ and trend $\mu$.

The initial backwards traversal of the algorithm functions to compute $x$ and $v$ conditional *only* on measurements associated with the descendants of each node. To begin, all entries of $x$ corresponding to tips are set to the observed measurements given by $y$. Further, because $y$ is assumed to represent exact measurements here, the corresponding entries of $v$ are set to 0. From here, the algorithm visits each internal node in the phylogeny, making sure to only visit a given node after all of its direct descendants have been visited (note that all tips were already visited in the initial step). For a given internal node descending from edge $i$, we can calculate $x_i$ and $v_i$ by treating each edge directly descending from $i$ as an independent "measurement" of the phenotype at the given node.

Specifically, each descending edge $j$ implies a normal distribution of measurements at the given node with mean $x_j - \mu l_j$ and variance $v_j + \sigma^2 l_j$, which follows from the fact that phenotypic changes over edge $j$ are normally distributed with mean $\mu l_j$ and variance $\sigma^2 l_j$ under the assumed Brownian Motion model. $x_i$ and $v_i$ are then given by the mean and variance of the product of normal probability density functions associated with each descending edge. In general, multiplying together the probability density functions of $c$ normal distributions with means $m$ and variances $s$ yields an unnormalized normal probability density function with variance $1/\sum_{j=1}^{c} s_j^{-1}$ (i.e., the reciprocal of the summed precisions—that is, the reciprocal of variance—for each distribution) and mean $(\sum_{j=1}^{c} s_j^{-1} m_j)/\sum_{j=1}^{c} s_j^{-1}$ (i.e., the precision-weighted average of means for each distribution; Petersen and Pedersen, 2012). Thus, the mean and variance of the phenotypic distribution at the focal node are ultimately given by:

$$x_i = \frac{\sum_{j \in \text{des}(i)} (v_j + \sigma^2 l_j)^{-1} (x_j - \mu l_j)}{\sum_{j \in \text{des}(i)} (v_j + \sigma^2 l_j)^{-1}} \tag{1}$$

$$v_i = \frac{1}{\sum_{j \in \text{des}(i)} (v_j + \sigma^2 l_j)^{-1}} \tag{2}$$

Where des($i$) is a shorthand for all edges directly descending from edge $i$.

After visiting every node in the phylogeny and calculating the mean and variance of the phenotypic distribution at the root, the algorithm moves on to the subsequent forwards traversal. Here, the algorithm updates $x/v$ based on measurements associated with tips that do *not* descend from each node and samples random values from these updated distributions, one node at a time. Beginning with the root descending from stem edge indexed $b$, note that $x_b$ and $v_b$ do not need to be updated because all tips in the phylogeny descend from the root by definition. Thus, the root-to-tips traversal begins with sampling a random phenotypic value at the root, $z$, from a normal distribution with mean $x_b$ and variance $v_b$, then replacing $x_b$ with $z$ and setting $v_b$ to 0 (reflecting the fact that we now assume the phenotypic value at the root is exactly $z$). From here, the algorithm proceeds to visit all other nodes in the phylogeny, ensuring that each node is visited only after its direct ancestor has been visited. For a given node descending from edge $i$, we can now think of edge $i$ itself as constituting an additional independent measurement of the phenotype at the focal node and update $x_i$ and $v_i$ accordingly:

$$x_i = \frac{v_i^{-1} x_i + (v_{\mathrm{anc}(i)} + \sigma^2 l_i)^{-1} (x_{\mathrm{anc}(i)} + \mu l_i)}{v_i^{-1} + (v_{\mathrm{anc}(i)} + \sigma^2 l_i)^{-1}} \quad (3)$$

$$v_i = \frac{1}{v_i^{-1} + (v_{\mathrm{anc}(i)} + \sigma^2 l_i)^{-1}} \quad (4)$$

Where anc($i$) is a shorthand for the edge directly ancestral to edge $i$. Note that the algorithm should skip updating $x_i/v_i$ if $v_i = 0$ (e.g., when edge $i$ is a tip), as $x_i$ corresponds to an exact phenotypic measurement in such cases and Eq. (4) would yield undefined results (more complicated procedures are needed in more general cases involving multivariate data, intraspecific variation/measurement error, and/or missing data, which we describe in the following subsection). As with the root node, the algorithm then samples a random phenotypic value for the given node, $z$, from a normal distribution with mean $x_i$ and variance $v_i$, replaces $x_i$ with $z$, and sets $v_i$ to 0. The last step highlights a subtle yet important point: distributions associated with edge $i$ are not necessarily independent of distributions for $i$'s sister edges. Specifically, because the means of these distributions all directly depend on $x_{\mathrm{anc}(i)}$ (i.e., the phenotypic value at the node ancestral to edge $i$), they will exhibit a covariance of $v_{\mathrm{anc}(i)}$. While most conventional ancestral state reconstruction methods must account for this non-independence among sister nodes, sampling exact phenotypic values for ancestral nodes ensures $v_{\mathrm{anc}(i)} = 0$, such that phenotypic distributions do not covary among sister nodes and the algorithm can update and sample from phenotypic distributions for each node independently.

After sampling a phenotypic value for the given node, the algorithm samples phenotypic values along edge $i$ before moving onto the next node. To do so, we must first define a set of time points at which to sample phenotypic values along edge $i$. Here, we place $p^* = \lceil l_i/dt - 1 \rceil$ equally-spaced time points along the edge, where $\lceil k \rceil$ denotes $k$ rounded up to the nearest integer. This ensures the interval between consecutive time points never exceeds a given "target" time interval of $dt$. For convenience, our implementation calculates $dt$ as $h/\xi$, where $h$ denotes the overall height of the phylogeny and $\xi$ is a user-specified "resolution" parameter controlling the overall density of time points across the phylogeny. To keep track of the specific times (measured forwards here, as in elapsed time since the root of the phylogeny) and sampled phenotypes associated with the time points along a given edge, let $t^*$ and $x^*$ denote $p^*$-length vectors of these respective quantities.

To calculate the means and variances of the phenotypic distributions for each time point along $i$, we can think of each time point as an additional node in the phylogeny with a single pair of adjacent edges—one ancestral and the other descendant. To begin, consider the first point on edge $i$ at time $t_1^*$ and let $t_a$ and $t_d$ denote the respective times at which the nodes immediately ancestral to and descending from $i$ occur. The mean and variance of the phenotypic distribution at $t_1$ based on the ancestral segment of edge $i$ are $x_{\text{anc}(i)} + \mu(t_1^* - t_a)$ and $\sigma^2(t_1^* - t_a)$, respectively, while those based on the descendant segment are $x_i - \mu(t_d - t_1^*)$ and $\sigma^2(t_d - t_1^*)$. Thus, the final mean and variance of the phenotypic distribution at $t_1^*$ are given by $m = (\frac{x_{\text{anc}(i)} + \mu(t_1^* - t_a)}{\sigma^2(t_1^* - t_a)} + \frac{x_i - \mu(t_d - t_1^*)}{\sigma^2(t_d - t_1^*)})/(\frac{1}{\sigma^2(t_1^* - t_a)} + \frac{1}{\sigma^2(t_d - t_1^*)})$ and $s = (\frac{1}{\sigma^2(t_1^* - t_a)} + \frac{1}{\sigma^2(t_d - t_1^*)})^{-1}$, respectively. After simplifying:

$$m = \frac{(t_d - t_1^*)x_{\text{anc}(i)} + (t_1^* - t_a)x_i}{t_d - t_a} \tag{5}$$

$$s = \frac{\sigma^2(t_1^* - t_a)(t_d - t_1^*)}{t_d - t_a} \tag{6}$$

These expressions correspond to the mean and variance of samples from a general Brownian bridge—that is, a Brownian Motion process with arbitrary rate/trend conditioned to start and end at particular values over a given time interval (Karatzas and

Shreve, 2004). We can now sample a random phenotypic value for $x_1^*$ from a normal distribution with mean $m$ and variance $s$ as given above. To sample phenotypic values for the other time points along edge $i$, we keep moving onto the next time point along the edge and repeating this procedure until the phenotypic value for the final $p^*$th time point has been sampled. Provided that the phenotypic value at the previous time point has been sampled already, the mean and variance of the phenotypic distribution at the $j$th time point are given by Eq. (6) after replacing $t_1^*$ with $t_j^*$, $t_a$ with $t_{j-1}^*$, and $x_{\text{anc}(i)}$ with $x_{j-1}^*$.

After completing the forwards traversal and sampling phenotypic values along the final edge of the phylogeny, the algorithm is complete. The final continuous stochastic character map is formed by concatenating the vector $x$ together with the $e$ distinct $x^*$ vectors (i.e., one for each edge). In our implementation, we organize the resulting map as a collection of $e$ vectors consisting of the sampled phenotypes along each edge. Specifically, the $i$th vector consists of the $x^*$ vector for the $i$th edge with $x_i$ tacked onto the end. Our implementation also returns the corresponding collection of $t^*$ vectors giving the times at which phenotypic values were sampled along each edge.

#### *Generalizing the algorithm: multivariate data and heterogeneity*

To effectively clarify the details of our continuous stochastic character mapping algorithm in its most flexible and general form, we will first establish some notation to help keep track of the various parameters and quantities generated/used by the algorithm. Assume we again have $e$ edges in our phylogeny (once again including a stem or root edge with lengths given by $e$-length vector $l$), though we now also have $s$ distinct regimes "painted" onto the phylogeny and data on $m$ continuous phenotypic variables. Each regime represents a discrete state with a potentially unique evolutionary rate/trend parameter, enabling modeling of among lineage trend/rate heterogeneity.

To outline our algorithm, it is useful to focus on the quantities associated with an individual edge denoted $i$. First, we define *two* sets of time points along edge $i$. First, we have $p_i^*$ "interpolant" points at equally-spaced times $t_i^*$, which directly correspond to the time points along each edge described in the previous subsection. Second, we now also define $p_i$ "critical" time points at times $t_i$, which consist of times corresponding to any regime shifts along edge $i$ as well as the descendant node of edge $i$. To keep track of regime information, let $r_i$ be another $p_i$-length vector of the regime preceding each critical time point along edge $i$. Broadly, the critical time points serve to simplify calculations involving rate/trend parameters by breaking up edges associated with multiple regimes into multiple edges each associated with a single regime (e.g., note how the specific steps of the forwards traversal described in step 2 below are repeated for each critical time point along each edge, rather than each edge). Because this blurs the distinction between nodes and points along edges, we generally found it most convenient to treat regime shift points and nodes together as a special class of time points in describing our algorithm.

Now let $X_i^*$ and $X_i$ be $p_i^* \times m$ and $p_i \times m$ matrices, respectively, of means of phenotypic distributions for each interpolant/critical time point along $i$ (i.e., each row corresponds to a time point, each column to a phenotype). The last row of $X_i$ correspond $x_i$ as defined in the previous section, while the rows of $X_i^*$ correspond to the individual entries of the $x^*$ vector for edge $i$. Finally, note that now any edge $i$—regardless of whether it is a tip or not—may have measurements associated with its descendant node: let $o$ be an $e$-length vector of integers representing the number of observations for each edge's descendant node (i.e., generally the number of sampled individuals/specimens, each with up to $m$ associated measurements for each phenotype; in the case that there are no observations for edge $i$, $o_i = 0$). Accordingly, let $Y_i$ be an $o_i \times m$ matrix of phenotypic measurements for each edge $i$.

Our algorithm samples values of $X_i$ and $X_i^*$ for each edge $i$ conditional on the phenotypic measurements/regime mappings described above ($Y_i$ and $r_i$, respectively, for each edge $i$). These phenotypic values are sampled from their joint distribution under an assumed multivariate Brownian Motion model with three main parameters: 1) $e$ $m \times m$

variance-covariance matrices describing intraspecific variation/measurement error at descendant nodes for each edge ($\Gamma$, which we term "node error" here as a generalization of the term "tip error" *sensu* Landis and Schraiber, 2017; see also Felsenstein, 2008; Hansen and Bartoszek, 2012); 2) $s$ $m$-length vectors describing deterministic changes in phenotypes per unit time for each regime ($\mu$, often referred to as evolutionary trends as in the previous subsection; see Hansen and Martins, 1996), and 3) $s$ $m \times m$ variance-covariance matrices describing stochastic changes in phenotypes per unit time for each regime ($\Sigma$, sometimes referred to as "evolutionary rate matrices" or "character variance-covariance matrices"). To do this, we additionally keep track of the $p_i$ variance-covariance matrices at the critical time points along each edge $i$, denoted $V_i$. Notably, the last matrix in $V_i$ corresponds to $v_i$ as defined in the prior subsection.

Below, we adopt a general notation of using subscripts to denote edge/regime indices first, followed by time points next (if applicable), and lastly specific phenotypic variables. For example, $X_i$ refers to the matrix of phenotypic values at critical time points along the $i$th edge, $X_{i,j}$ to its $j$th row (corresponding to the $j$th entry of $t_i$ or time $t_{i,j}$), and $X^*_{i,j,p}$ to $p$th value in this row (corresponding to the $p$th phenotypic variable). Similarly, $\mu_i$ would refer to evolutionary trends for the $i$th regime and $\mu_{i,j}$ to the trend for the $j$th phenotypic variable specifically.

Notably, our description repeatedly redefines/uses the labels $Z$ and $W$ to refer to temporary variables used in intermediate calculations. $Z$ and $W$ generally denote (matrices of) phenotypic means and (vectors of) variance-covariance matrices, respectively, at a given focal time points conditional on the phenotypic information associated with a particular observation or adjacent edge. As in the previous subsection, these (multivariate) normal probability density functions are implicitly multiplied together to calculate the final phenotypic distribution for the focal time point. Similarly to the univariate case, the product of $c$ multivariate normal density functions with means $\vec{m}$ and variance-covariance matrices $S$ is given by an unnormalized multivariate normal density function with variance-covariance matrix $(\sum_{j=1}^{c} S_j^{-1})^{-1}$ (i.e., the inverse of the summed precision matrices, which themselves are given by the inverse of the variance-covariance matrices) and mean $(\sum_{j=1}^{c} S_j^{-1})^{-1} \sum_{j=1}^{c} S_j^{-1}\vec{m}_j$ (the precision matrix-weighted average of means).

1) Complete the backwards traversal over all edges until reaching the "stem edge"/root. For each edge denoted $i$ with $c_i$ immediate descendant edges:

1a) Initialize calculations for edge $i$ by defining $Z$ as an $(o_i + c_i) \times m$ matrix and setting the first $o_i$ rows of $Z$ to $Y_i$. Similarly, define $W$ as $o_i + c_i$-length vector of $m \times m$ matrices and set the first $o_i$ matrices to $\Gamma_i$. Any missing entries in $Z$ are set to 0, and corresponding diagonal and off-diagonal entries of the associated $W$ matrices are set to $\infty$ and 0, respectively (Hassler et al., 2022). Accordingly, if edge $i$ is a tip with no associated measurements (i.e., $o_i + c_i = 0$), we can encode this missing data by setting $o_i$ to 1 and initializing $Z$ as a $1 \times m$ matrix of 0s and $W$ as a single $m \times m$ matrix with $\infty$ along the diagonal. Otherwise, if edge $i$ has immediate descendant edges (i.e., not a tip with $c_i > 0$), for each descendant edge indexed $j$ from 1 to $c_i$:

$$Z'_{o_i+j} = X'_{d_j,1} - \mu_{r_{d_j,1}}(t_{d_j,1} - t_{i,p_i}) \tag{7}$$

$$W_{o_i+j} = V_{d_j,1} + \Sigma_{r_{d_j,1}}(t_{d_j,1} - t_{i,p_i}) \tag{8}$$

Where $M'$ denotes the transpose of a matrix/vector (only used here to convert between row and column vectors), $Z_j$ denotes the $j$th row of $Z$, $W_j$ denotes the $j$th matrix of $W$, and $d$ is a $c_i$-length vector of indices corresponding to the edges immediately descending from $i$. Now the rows of $Z$ and matrices of $W$ correspond to the means and variance-covariance matrices of the phenotypic distribution at edge $i$'s descendant node conditional on any given observation for or edge descending from $i$. Thus, these represent the "measurement distributions" that must be multiplied together to compute the phenotypic distribution at $i$'s descendant node.

1b) Calculate the phenotypic distribution for $i$'s descendant node. As in the previous section, this is given by product of the probability density functions associated with each descending edge and/or observation given above. Again, we can calculate this multivariate normal distribution's mean and variance-covariance matrix by taking a precision matrix-weighted (i.e., the inverse of matrices in $W$) average of the means (i.e., the rows of $Z$) and the inverse of the summed precision matrices:

$$V_{i,p_i} = \left( \sum_{j=1}^{o_i+c_i} W_j^{-1} \right)^{-1} \tag{9}$$

$$X'_{i,p_i} = V_{i,p_i} \sum_{j=1}^{o_i+c_i} W_j^{-1} Z'_j \tag{10}$$

Where $W_j^{-1}$ specifically denotes the "pseudo-inverse" of $W_j$ (Hassler et al., 2022). Notably, Eq. (10) may be undefined when one or more measurements are missing or assumed to be exact, which can cause entries of $V^*_{i,p_i}$ and $W^{-1}$, respectively, to be $\infty$. The former case can be addressed by simply defining $\infty * 0 = 0$ (Hassler et al., 2022). The latter case does not admit a simple workaround, so our algorithm accurately approximates Eq. (10) by replacing all infinite entries in $W^{-1}$ with an arbitrarily large number $g$. To ensure accuracy, $g$ should generally be at least 2–4 orders magnitude larger than the reciprocal of the lowest non-zero evolutionary rate and/or node error variance under the assumed Brownian Motion model (by default, our implementation sets $g = 10^{10}$). If we use $\tilde{W}^{-1}$ to denote $W^{-1}$ with all infinite entries replaced by $g$, we can write a more robust version of Eq. (10):

$$X'_{i,p_i} = \left( \sum_{j=1}^{o_i+c_i} \tilde{W}_j^{-1} \right)^{-1} \sum_{j=1}^{o_i+c_i} \tilde{W}_j^{-1} Z'_j \tag{11}$$

Using $\tilde{W}^{-1}$ to calculate $X$ but not $V$ results in averaging any exact yet contradictory measurements while ensuring the resulting average is also considered exact. While somewhat paradoxical, this prevents approximation errors from accumulating in $V$ as the algorithm proceeds to other nodes in the phylogeny.

1c) If edge $i$ includes any regime shifts (i.e., $p_i \geqslant 2$), calculate phenotypic distributions for each preceding critical time point along edge $i$ using equations that follow from how phenotypic changes over time are distributed under the assumed Brownian Motion model. For each critical point $j$ from $p_i - 1$ to 1 (note that $p_i - 1$ will always be $\geqslant 1$ here because this step only occurs if $p_i \geqslant 2$):

$$V_{i,j} = V_{i,j+1} + \Sigma_{r_{i,j+1}}(t_{i,j+1} - t_{i,j}) \tag{12}$$

$$X'_{i,j} = X'_{i,j+1} - \mu_{r_{i,j+1}}(t_{i,j+1} - t_{i,j}) \tag{13}$$

2) Complete the forward traversal over all edges. Starting with the stem edge indexed $b$, sample a value for the root node from a multivariate normal distribution with mean $X_{b,1}$ and variance-covariance matrix $V_{b,1}$ (we assume the stem edge is of 0 length so it consists of a single critical time point corresponding to the root of the phylogeny and no interpolant points). As in the prior subsection, there is no need to update the phenotypic distribution at the root prior to sampling because all nodes in the phylogeny descend from the root by definition. Then, for each critical time point $j$ along edge $i$:

2a) Initialize calculations by redefining $Z$ as an $m$-length row vector and $W$ as a single $m \times m$ matrix. Here, $Z$ and $W$ represent counterparts to $X_{i,j}$ and $V_{i,j}$.

While $X_{i,j}$ and $V_{i,j}$ define the distribution of phenotypic values at the $j$th critical time point along edge $i$ based *only* on descendants of edge $i$, $Z$ and $W$ are based *only* on all *non*-descendants of $i$. Because this is a forwards traversal, phenotypic values for the immediately previous critical time point have already been sampled and $Z/W$ are given by:

$$Z' = X'_{i,j-1} + \mu_{r_{i,j}}(t_{i,j} - t_{i,j-1}) \tag{14}$$

$$W = \Sigma_{r_{i,j}}(t_{i,j} - t_{i,j-1}) \tag{15}$$

In this is the first critical time point along edge $i$ (i.e., $j - 1 = 0$), we must instead use the phenotypic values and time associated with the last critical time point (i.e., descendant node) of the edge directly ancestral to $i$, denoted $a$. Specifically, $X_{i,j-1}$ and $t_{i,j-1}$ are substituted with $X_{a,p_a}$ and $t_{a,p_a}$, respectively, in the expression above.

2b) Update the phenotypic distribution for the $j$th critical time point along edge $i$. Once again, this corresponds to taking the product of the existing phenotypic distribution at the $j$th critical time point—calculated during the backwards traversal in step 1—with the new phenotypic distribution based on the phenotypic value sampled for the previous critical time point:

$$V_{i,j} = \left(W^{-1} + V_{i,j}^{-1}\right)^{-1} \tag{16}$$

$$X'_{i,j} = \left(\tilde{W}^{-1} + \tilde{V}_{i,j}^{-1}\right)^{-1} \left(\tilde{W}^{-1} Z' + \tilde{V}_{i,j}^{-1} X'_{i,j}\right) \tag{17}$$

Where, as in Eq. (11), $\tilde{W}^{-1}$ and $\tilde{V}_{i,j}^{-1}$ respectively denote $W^{-1}$ and $V_{i,j}^{-1}$ but with any infinite entries replaced by an arbitrarily large number $g$ ($10^{10}$ in our implementation by default).

2c) Sample phenotypic values at the $j$th critical time point along edge $i$ by replacing

$X_{i,j}$ with new values sampled from a multivariate normal distribution with mean $X_{i,j}$ and variance-covariance matrix $V_{i,j}$.

2d) Sample phenotypic values at all interpolant time points along edge $i$ lying between the $j$th and last critical time point. As in step 2.1, if this is the first critical time point along edge $i$ (i.e., $j-1=0$), we use the last critical time point/descendant node of ancestral edge $a$. For each interpolant time point $k$ such $t_{i,j-1} < t^*_{i,k} < t_{i,j}$ (if $j-1=0$, swap $t_{i,j-1}$ with $t_{a,p_a}$ as indicated above), sample $X^*_{i,k}$ from a multivariate normal distribution with mean $Z$ and variance-covariance matrix $W$, now redefined as:

$$Z = \frac{(X_{i,j} - X^*_{i,k-1})(t^*_{i,k} - t^*_{i,k-1})}{t_{i,j} - t^*_{i,k-1}} + X^*_{i,k-1} \tag{18}$$

$$W = \frac{(t_{i,j} - t^*_{i,p})(t^*_{i,k} - t^*_{i,k-1})}{t_{i,j} - t^*_{i,k-1}} \Sigma_{r_{i,j}} \tag{19}$$

If this is the first interpolant time point along edge $i$, both $k-1$ and $j-1=0$, so $X^*_{i,k-1}$ and $t^*_{i,k-1}$ are accordingly swapped with the phenotypic values and time associated with the last critical time point/descendant node of ancestral edge $a$ as above (i.e., $X_{a,p_a}$ and $t_{a,p_a}$, respectively). Otherwise, in the case that the last interpolant time point precedes the previous critical time point (i.e., $t^*_{i,k-1} < t_{i,j-1}$), we instead use the phenotypic values and time associated with the previous critical time point $j-1$. Specifically, $X^*_{i,k-1}$ and $t^*_{i,k-1}$ in the above expression are swapped with $X_{i,j-1}$ and $t_{i,j-1}$, respectively. Note that the above formulae are a multivariate generalization of Eq. (6) (i.e., the mean and variance-covariance matrix of samples from a multivariate Brownian bridge).

We can now combine $X^*$ and $X$ to form a single continuous stochastic character map. Our implementation arranges these values into a collection *em* vectors consisting of the values sampled at both critical and interpolant time points for each phenotype along each edge. We form each of these vectors by first splitting the columns $X^*$ and $X$ into individual vectors, then inserting the phenotypic values at critical time points into the corresponding vector of values at interpolant points, arranged such that the entries occur in chronological order.

#### *Extending the algorithm: mapping under "evolving rates" models*

We developed a method for generating continuous stochastic character maps based on posterior samples of parameters under models fitted via the *evorates* package (Martin et al., 2023). Models implemented in the *evorates* package work by assuming the rate parameter of a Brownian Motion model of trait evolution itself "evolves" according to a constant-rate geometric Brownian Motion process (i.e., the natural log of the evolutionary rates evolve according to a constant-rate Brownian Motion process). Thus, the model infers a "rate evolution process" controlled by a trend ($\mu_{\sigma^2}$) parameter determining whether rates tend to decrease or increase over time, as well as a rate variance ($\sigma^2_{\sigma^2}$) parameter controlling how quickly random variation in rates accumulates over time. Additionally, the model estimates both the rate at the root of the phylogeny ($\sigma^2_0$) and average rates of evolution along each branch of the phylogeny, which we term "branchwise rates" or $\overline{\sigma^2_i}$, where $i$ denotes the index of a particular edge. To generate continuous stochastic character maps under such models, we implemented a two-step procedure whereby the inferred rate evolution process and branchwise rates are first used to generate maps of evolutionary rates at a user-specified resolution, which are in turn used to generate maps of trait (or factor) values of the same resolution by treating the rate maps as high-resolution regime maps (i.e., a regime for each time point spanning the preceding time interval). We summarize this process graphically in Fig. S22.

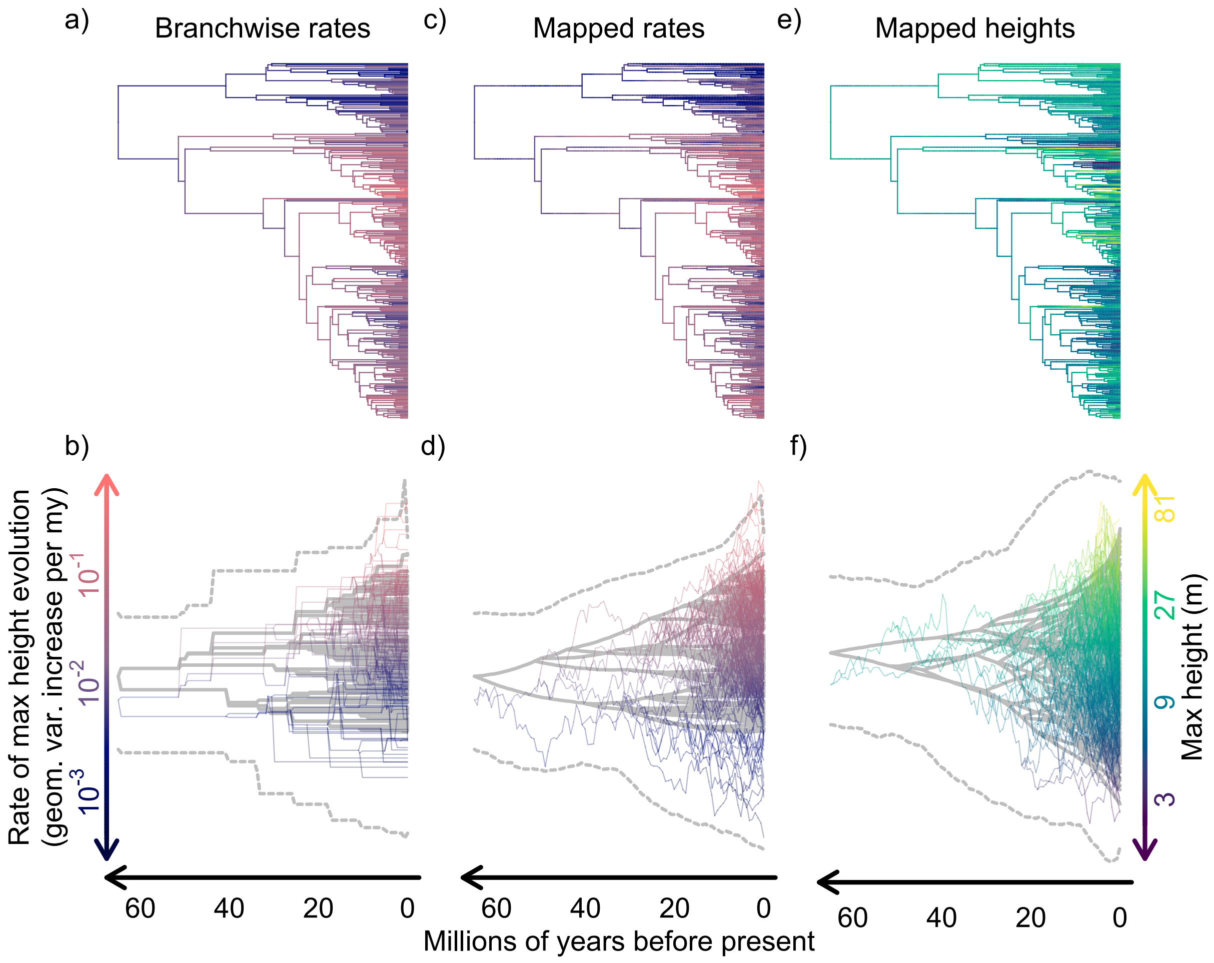

Figure S22. A brief graphical summary of our procedure for generating continuous stochastic character maps under "evolving rates" or evorates models, using our fitted model of maximum height evolution across eucalypts as an example. The phenograms in panels b, d, and f depict the distributions of: inferred average evolutionary rates (i.e., branchwise rates), which form the main input for our procedure (panel b); 2) the mapped rate values sampled under the inferred rate evolution process conditional on branchwise rates, which are generated by the first step of our procedure (panel d); and 3) mapped height values sampled conditional on both the observed maximum height data at the tips and mapped rates, which are generated by the second step of our procedure (panel f). Each phenogram in panels b, d, and f consists of the overall mean rates/heights for each lineage in solid gray lines, as well as a single posterior sample/associated continuous stochastic character map in thinner, transparent lines, which are colored according to their position along the vertical axis (different color gradients are used to visually distinguish rates from heights). The phylograms in panels a, c, and e provide an alternate view of the posterior sample/associated continuous stochastic character map, with rate/height values represented solely by color gradients rather than positions along vertical axes. The gray dashed lines in each phenogram in panels b, d, and f depict the overall range of rate/heights, and were derived from calculating 95% confidence intervals along each lineage, then taking the minimum and maximum interval bounds at 100 equally-spaced time points spanning the height of the phylogeny.

Because the time-average of a geometric Brownian Motion process has no closed-form probability distribution (Lepage et al., 2007), there is no simple and/or exact method for sampling rate values at arbitrary time points across a phylogeny conditional on the inferred rate evolution process and branchwise rates. However, we can take advantage of the approximation of geometric Brownian Motion time-averages used by the *evorates* package (see section "Approximating Geometric Brownian Motion Time-Averages" in the Online Appendix for Martin et al., 2023; see also Dufresne, 2004; Welch and Waxman, 2008). Specifically, we assume each branchwise rate is the sum of a trend and noise component, with the noise component assumed to follow the distribution of geometric, rather than arithmetic, time-averages of an untrended (i.e., $\mu_{\sigma^2} = 0$) geometric Brownian Motion process. Under this assumption, we can derive straight-forward normal distributions describing how the natural log of rates at the nodes of a phylogeny are distributed. To start out, let $\ln \sigma^2_{t_1}$ and $\ln \sigma^2_{t_2}$ represent the natural log of the starting and ending values of a geometric Brownian Motion process over an interval of length $l$ with trend $\mu_{\sigma^2}$, rate variance $\sigma^2_{\sigma^2}$, and time-average $\overline{\sigma^2}$. Then the distributions of $\ln \sigma^2_{t_1}$ and $\ln \sigma^2_{t_2}$ under the aforementioned approximation are given by:

$$\ln \sigma^2_{t_1} \sim \mathrm{N}\left(\ln \overline{\sigma^2} - \beta,\ \frac{\sigma^2_{\sigma^2} l}{3}\right) \tag{20}$$

$$\ln \sigma^2_{t_2} \sim \mathrm{N}\left(\ln \overline{\sigma^2} - \beta + \mu_{\sigma^2} l,\ \frac{\sigma^2_{\sigma^2} l}{2}\right) \tag{21}$$

$$\beta = \begin{cases} 0 & \text{if } \mu_{\sigma^2} = 0 \\ \ln |\mathrm{e}^{\mu_{\sigma^2} l} - 1| - \ln |\mu_{\sigma^2}| - \ln l & \text{if } \mu_{\sigma^2} \neq 0 \end{cases} \tag{22}$$

Where $\mathrm{N}(\mu,\ \sigma^2)$ denotes a normal distribution with mean $\mu$ and variance $\sigma^2$. We can generalize these results to define normal distributions for the natural log of rates at each node in a phylogeny given branchwise rates along each edge based on basic algebra of normal random variables. Specifically, Let $\sigma^2_i$ represent the rate at the node immediately descending from edge $i$ (not to be confused with $\overline{\sigma^2_i}$, the branchwise rate along edge $i$), then:

$$\ln \sigma_i^2 \sim \mathrm{N}\left(\alpha_i^{-1}\left(\frac{2(\ln \overline{\sigma_i^2} - \beta_i)}{l_i} + \sum_{d \in \mathrm{des}(i)} \frac{3(\ln \overline{\sigma_d^2} - \beta_d)}{L_d}\right) + \mu_{\sigma^2} t_i,\ \alpha_i^{-1}\sigma_{\sigma^2}^2\right) \tag{23}$$

$$\alpha_i = \frac{2}{l_i} + \sum_{d \in \mathrm{des}(i)} \frac{3}{l_d} \tag{24}$$

$$\beta_i = \begin{cases} 0 & \text{if } \mu_{\sigma^2} = 0 \\ \ln\left|\mathrm{e}^{\mu_{\sigma^2} t_i} - \mathrm{e}^{\mu_{\sigma^2} t_{\mathrm{anc}(i)}}\right| - \ln|\mu_{\sigma^2}| - \ln l_i & \text{if } \mu_{\sigma^2} \neq 0 \end{cases} \tag{25}$$

Where $l_i$ now denotes the length of edge $i$, $t_i$ the height (i.e., distance from root) of the node immediately descending from edge $i$, $\mathrm{des}(i)$ the indices for all edges immediately descending from edge $i$, and $\mathrm{anc}(i)$ the index for the edge immediately ancestral to edge $i$. Based on these formulae, we implemented a forward traversal algorithm that generates continuous stochastic character maps of rates by jointly sampling rate values at the time points along each edge conditional on each edge's branchwise rate and sampled rate value at its ancestral node. Under our approximation, the joint distribution of the natural log of rate values along a given edge follows a straight-forward multivariate normal distribution conditioned to have a particular weighted sum given by the corresponding branchwise rate. Let $\vec{\sigma_i^2}$ and $\vec{t_i}$ denote $p_i$-length vectors of the rate values and corresponding time points along edge $i$ (i.e., the $p_i$th entries correspond to the node immediately descending from edge $i$; $\sigma_i^2 = \vec{\sigma^2}_{i,p_i}$ and $t_i = \vec{t}_{i,p_i}$). Ignoring the condition that $\vec{\sigma_i^2}$ must have a particular weighted sum, the distribution of $\vec{\sigma_i^2}$ is given by:

$$\ln \vec{\sigma_i^2} \sim \mathrm{MVN}\left(\frac{(\mathbb{E}[\ln \sigma_i^2] - \ln \sigma_{\mathrm{anc}(i)}^2)(\vec{t_i} - t_{\mathrm{anc}(i)})}{l_i} + \ln \sigma_{\mathrm{anc}(i)}^2,\ \Sigma_i\right) \tag{26}$$

$$\Sigma_{i,j,k} = \frac{\sigma_{\sigma^2}^2 (t_i - \max\{\vec{t}_{i,j},\ \vec{t}_{i,k}\})(\min\{\vec{t}_{i,j},\ \vec{t}_{i,k}\} - t_{\mathrm{anc}(i)}) + \mathrm{Var}(\ln \sigma_i^2)\sqrt{\vec{t}_{i,j}\vec{t}_{i,k}}}{l_i} \tag{27}$$

Where $\mathrm{MVN}(\vec{\mu},\ \Sigma)$ denotes a multivariate normal distribution with mean vector $\vec{\mu}$ and variance-covariance matrix $\Sigma$, $\mathbb{E}[\ln \sigma_i^2]$ and $\mathrm{Var}(\ln \sigma_i^2)$ are given by Eq. (23), and $\Sigma_{i,j,k}$ is the entry in the $j$th row and $k$th column of matrix $\Sigma_i$. Notably, for edges descending from the root node, $\ln \sigma^2_{\text{anc}(i)}$ corresponds to the inferred root rate parameter, $\ln \sigma^2_0$, and $t_{\text{anc}(i)}$ to 0. We can calculate an approximate branchwise rate for a given sample of $\vec{\sigma_i^2}$ as the weighted sum:

$$\widehat{\overline{\sigma_i^2}} = \frac{1}{l_i} \sum_{j=1}^{p_i} \vec{\sigma_i^2}(\vec{t_{i,j}} - \vec{t_{i,j-1}}) \tag{28}$$

Where $\vec{t_{i,0}}$ is replaced with $t_{\text{anc}(i)}$ in the above expression (or with 0 if $i$ is an edge descending from the root node). Here, we assume each sampled rate at a given time point is constant along its preceding time interval, which simplifies calculations and negligibly differs from more accurate interpolations of rates along an edge given a sufficiently dense set of time points (notably, we use the same approximation to calculate likelihoods of continuous factor-dependent Brownian Motion models).

We use Markov chain Monte Carlo to sample from Eq. (26) under an informative prior conditioning the sampled branchwise rate, $\widehat{\overline{\sigma_i^2}}$, to approximately equal the inferred branchwise rate, $\overline{\sigma_i^2}$. Specifically, we place a normal prior on $\ln \widehat{\overline{\sigma_i^2}}$ with mean $\ln \overline{\sigma_i^2}$ and a user-specified variance or "tolerance", which defaults to 0.001. Lower tolerance values will produce more accurate rate samples at the cost of increased computation time (and vice versa for higher values). To simplify this procedure, we use an "uncentered" parameterization and do not sample $\vec{\sigma_i^2}$ directly. Instead, we sample $p_i$ independent standard normal random variables, denoted $z$ here, which are transformed to follow the distribution given by Eq. (26) via $\Sigma_i^{\frac{1}{2}} z + \mathbb{E}[\ln \vec{\sigma_i^2}]$, where $\Sigma_i^{\frac{1}{2}}$ denotes the lower triangular Cholesky decomposition of $\Sigma_i$ (Betancourt and Girolami, 2019). We propose new samples of $z$ by simply adding $p_i$ normal random variables with mean 0 and variance 0.01 to the previously sampled values of $z$. For each edge, Markov chain Monte Carlo sampling of rate values is run for a user-specified maximum number of iterations (which defaults to 100,000), but terminates early if $(\ln \widehat{\overline{\sigma_i^2}} - \ln \overline{\sigma_i^2})^2$ is less than the user-specified tolerance value (i.e., the sampled branchwise rate is within a standard deviation of the normal prior's mean). So far, we have found that this procedure essentially never reaches the maximum number of iterations under our default settings, ensuring a close match between inferred branchwise rates and the sampled branchwise rates in the resulting continuous stochastic character maps of rates.

Finally, after generating rate maps, we can assume that the sampled rates at each time point are constant along the preceding interval, forming a high-resolution map of regimes corresponding to different rates for each time interval (i.e., all time points are considered critical time points here). Then, we use the generalized continuous stochastic character mapping algorithm described in the prior subsection to sample trait/factor values at all time points conditional on the mapped rates and observed trait/factor data.

### Validation Study

#### *Derivation of joint and conditional distributions*

To derive the expected multivariate normal distribution of all phenotypic values across the phylogeny conditional on observed measurements under an assumed Brownian Motion model, we first calculated the joint distribution of both phenotypic values and observed measurements, then used well-established formulae to compute the corresponding conditional distribution of phenotypic values given exact values for the observed measurements (Petersen and Pedersen, 2012; see also Jhwueng, 2021). In our case, let the mean vector and variance-covariance matrix of the joint multivariate normal distribution respectively be:

$$M = \begin{bmatrix} M_p \\ M_o \end{bmatrix},\ \Sigma = \begin{bmatrix} \Sigma_{pp} & \Sigma_{po} \\ \Sigma_{op} & \Sigma_{oo} \end{bmatrix} \tag{29}$$

Where the $p$ subscript denotes the entries of $M$ and rows/columns of $\Sigma$ corresponding to all time points (including nodes/tips) across the phylogeny, while subscript $o$ denotes entries/rows/columns corresponding to observed measurements. The mean vector and variance-covariance matrix of the conditional distribution at all time points given $M_o = Y$ are then respectively given by (Petersen and Pedersen, 2012):

$$\overline{M_p} = M_p + \Sigma_{po}\Sigma_{oo}^{-1}Y,\ \overline{\Sigma_{pp}} = \Sigma_{pp} - \Sigma_{po}\Sigma_{oo}^{-1}\Sigma_{op} \tag{30}$$

To compute the joint distribution of phenotypic values at all time points across the phylogeny and observed measurements (i.e., $M$ and $\Sigma$ in Eq. (29)), we start by expanding the conventional phylogenetic variance-covariance matrix describing the phenotypic distribution at all nodes/tips of a given phylogeny under a Brownian Motion model, appending additional rows/columns corresponding to the time points along each edge and any observed measurements. The (co)variances for each time point can be derived by viewing time points as additional internal nodes. Similarly, observed measurements can be seen as arising from additional edges of length 1 appended to their associated tips, with "evolutionary rates" along these additional edges given by node error rather than the evolutionary rate parameter (e.g., Felsenstein, 2008). This may be further extended to a multivariate context via the usual Kronecker product formulations with a given evolutionary rate matrix (e.g., Caetano and Harmon, 2019), taking care to respect the "shift" from evolutionary rate matrices to variance-covariance matrices describing node error along the appended edges corresponding to observed measurements. Assuming a flat prior over all possible phenotypic values at the root (as our algorithm does by default), the associated mean vector simply consists of 0s plus the evolutionary trend parameter multiplied by the distance between each time point and the root (note that any evolutionary trends "shift" to 0 along appended edges corresponding to observed measurements). If we denote the resulting variance-covariance matrix $\Sigma^*$, the final $\Sigma$ is then technically given by $\lim_{a\to\infty} a + \Sigma^*$ (note here that $a$ is added to *each* individual entry in $\Sigma^*$). In practice, it suffices to simply take $a$ to be an arbitrarily large number—we set $a = 10^4$ for our validation study. Higher values of $a$ tended to degrade the numerical stability of R's linear algebra functions, which we used to compute $\overline{M_p}$ and $\overline{\Sigma_{pp}}$ from $M$ and $\Sigma$.

#### *Simulation conditions*

For each simulation in the validation study, we simulated phenotypic measurements under a Brownian Motion process on either a random ultrametric or non-ultrametric tree (using *phytools* or *ape*, respectively; Revell, 2012; Paradis and Schliep, 2019), scaled to unit height with the number of tips randomly sampled from log-uniform distribution bounded between 10 and 100. Evolutionary rates were randomly sampled from a log-uniform distribution bounded between 0.1 and 10, while intraspecifc/measurement error variances (if included in the simulation condition) were sampled from a narrower log-uniform distribution bounded between 0.03 and 3. If the simulation included evolutionary trends, these were sampled from a uniform distribution bounded between -5 and 5. If the simulation included more than one phenotype, we independently sampled rates, node error variances, and/or trends for each phenotype. To simulate correlations among phenotypes, we sampled correlation matrices—one for evolutionary rates and another for node error if applicable—from a flat Lewandowski-Kurowicka-Joe distribution (i.e., $\eta = 1$; (Lewandowski et al., 2009)) using the *rethinking* R package (McElreath, 2025). If the simulation included multiple observations per tip, we independently sampled the number of observations per tip from a Poisson distribution with mean/$\lambda = 3$, yielding ∼0–7 observations per tip. To simulate missing data, we randomly removed either 0, 5, or 25% of measurements from the final trait dataset. In half of cases, we generated continuous stochastic character maps under Brownian Motion models with the same rate, node error, and trend parameters used to simulate the data. For the other half of cases, we instead generated maps under a Brownian Motion model with random parameters (sampling parameters as described above). Ultimately, we defined 360 distinct simulation scenarios (Table S20), simulating 10 replicates per scenario. For each replicate, we generated 100 continuous stochastic character maps with a random resolution sampled from a log-uniform distribution bounded between 20 and 500.

Table S20. *All 360 simulation conditions used to validate the correctness of our continuous stochastic character mapping algorithm. For each condition, we additionally report the proportion of individual simulations (10 total per condition) which signficantly differed (i.e., p-value* $< 0.05$*) from their expected distribution based on one-sample Kolmogorov-Smirnov tests. Single and mult. refer to conditions whereby one or ∼3 measurements of each phenotype are available per tip, respectively, while w/ error indicates conditions where we simulated intraspecific variation/measurement error in observed phenotypic measurements. Conditions with identical assumed parameters used the same parameters to both simulate data and generate continuous stochastic character maps/compute expected distributions, while those with random assumed parameters generated continuous stochastic character maps/computed expected distributions based on randomly sampled parameters.*

| ultrametric phylogenies? | number of phenotypes | phenotypic correlations? | evolutionary trends? | measurement type | proportion missing data | assumed parameters | proportion $p < 0.05$ |
|---|---|---|---|---|---|---|---|
| × | 1 | × | × | single exact | 0% | identical | 10% |
| ✓ | 1 | × | × | single exact | 0% | identical | 0% |
| × | 2 | × | × | single exact | 0% | identical | 20% |
| ✓ | 2 | × | × | single exact | 0% | identical | 10% |
| × | 3 | × | × | single exact | 0% | identical | 10% |
| ✓ | 3 | × | × | single exact | 0% | identical | 10% |
| × | 2 | ✓ | × | single exact | 0% | identical | 0% |
| ✓ | 2 | ✓ | × | single exact | 0% | identical | 20% |
| × | 3 | ✓ | × | single exact | 0% | identical | 0% |
| ✓ | 3 | ✓ | × | single exact | 0% | identical | 0% |
| × | 1 | × | ✓ | single exact | 0% | identical | 10% |
| ✓ | 1 | × | ✓ | single exact | 0% | identical | 10% |
| × | 2 | × | ✓ | single exact | 0% | identical | 20% |
| ✓ | 2 | × | ✓ | single exact | 0% | identical | 0% |
| × | 3 | × | ✓ | single exact | 0% | identical | 20% |
| ✓ | 3 | × | ✓ | single exact | 0% | identical | 0% |
| × | 2 | ✓ | ✓ | single exact | 0% | identical | 10% |
| ✓ | 2 | ✓ | ✓ | single exact | 0% | identical | 0% |
| × | 3 | ✓ | ✓ | single exact | 0% | identical | 10% |
| ✓ | 3 | ✓ | ✓ | single exact | 0% | identical | 0% |
| × | 1 | × | × | single w/ error | 0% | identical | 10% |
| ✓ | 1 | × | × | single w/ error | 0% | identical | 10% |
| × | 2 | × | × | single w/ error | 0% | identical | 20% |
| ✓ | 2 | × | × | single w/ error | 0% | identical | 10% |
| × | 3 | × | × | single w/ error | 0% | identical | 0% |
| ✓ | 3 | × | × | single w/ error | 0% | identical | 20% |
| × | 2 | ✓ | × | single w/ error | 0% | identical | 0% |
| ✓ | 2 | ✓ | × | single w/ error | 0% | identical | 0% |
| × | 3 | ✓ | × | single w/ error | 0% | identical | 0% |
| ✓ | 3 | ✓ | × | single w/ error | 0% | identical | 0% |
| × | 1 | × | ✓ | single w/ error | 0% | identical | 0% |
| ✓ | 1 | × | ✓ | single w/ error | 0% | identical | 10% |
| × | 2 | × | ✓ | single w/ error | 0% | identical | 0% |
| ✓ | 2 | × | ✓ | single w/ error | 0% | identical | 10% |
| × | 3 | × | ✓ | single w/ error | 0% | identical | 0% |
| ✓ | 3 | × | ✓ | single w/ error | 0% | identical | 0% |
| × | 2 | ✓ | ✓ | single w/ error | 0% | identical | 0% |
| ✓ | 2 | ✓ | ✓ | single w/ error | 0% | identical | 0% |
| × | 3 | ✓ | ✓ | single w/ error | 0% | identical | 0% |
| ✓ | 3 | ✓ | ✓ | single w/ error | 0% | identical | 0% |
| × | 1 | × | × | mult. w/ error | 0% | identical | 0% |
| ✓ | 1 | × | × | mult. w/ error | 0% | identical | 0% |
| × | 2 | × | × | mult. w/ error | 0% | identical | 0% |
| ✓ | 2 | × | × | mult. w/ error | 0% | identical | 0% |
| × | 3 | × | × | mult. w/ error | 0% | identical | 10% |
| ✓ | 3 | × | × | mult. w/ error | 0% | identical | 0% |
| × | 2 | ✓ | × | mult. w/ error | 0% | identical | 0% |
| ✓ | 2 | ✓ | × | mult. w/ error | 0% | identical | 0% |
| × | 3 | ✓ | × | mult. w/ error | 0% | identical | 10% |
| ✓ | 3 | ✓ | × | mult. w/ error | 0% | identical | 0% |
| × | 1 | × | ✓ | mult. w/ error | 0% | identical | 0% |
| ✓ | 1 | × | ✓ | mult. w/ error | 0% | identical | 10% |

| ultrametric phylogenies? | number of phenotypes | phenotypic correlations? | evolutionary trends? | measurement type | proportion missing data | assumed parameters | proportion $p < 0.05$ |
|---|---|---|---|---|---|---|---|
| × | 2 | × | ✓ | mult. w/ error | 0% | identical | 0% |
| ✓ | 2 | × | ✓ | mult. w/ error | 0% | identical | 0% |
| × | 3 | × | ✓ | mult. w/ error | 0% | identical | 20% |
| ✓ | 3 | × | ✓ | mult. w/ error | 0% | identical | 20% |
| × | 2 | ✓ | ✓ | mult. w/ error | 0% | identical | 0% |
| ✓ | 2 | ✓ | ✓ | mult. w/ error | 0% | identical | 10% |
| × | 3 | ✓ | ✓ | mult. w/ error | 0% | identical | 10% |
| ✓ | 3 | ✓ | ✓ | mult. w/ error | 0% | identical | 10% |
| × | 1 | × | × | single exact | 5% | identical | 0% |
| ✓ | 1 | × | × | single exact | 5% | identical | 0% |
| × | 2 | × | × | single exact | 5% | identical | 10% |
| ✓ | 2 | × | × | single exact | 5% | identical | 0% |
| × | 3 | × | × | single exact | 5% | identical | 0% |
| ✓ | 3 | × | × | single exact | 5% | identical | 0% |
| × | 2 | ✓ | × | single exact | 5% | identical | 10% |
| ✓ | 2 | ✓ | × | single exact | 5% | identical | 0% |
| × | 3 | ✓ | × | single exact | 5% | identical | 0% |
| ✓ | 3 | ✓ | × | single exact | 5% | identical | 10% |
| × | 1 | × | ✓ | single exact | 5% | identical | 0% |
| ✓ | 1 | × | ✓ | single exact | 5% | identical | 0% |
| × | 2 | × | ✓ | single exact | 5% | identical | 10% |
| ✓ | 2 | × | ✓ | single exact | 5% | identical | 10% |
| × | 3 | × | ✓ | single exact | 5% | identical | 10% |
| ✓ | 3 | × | ✓ | single exact | 5% | identical | 0% |
| × | 2 | ✓ | ✓ | single exact | 5% | identical | 0% |
| ✓ | 2 | ✓ | ✓ | single exact | 5% | identical | 20% |
| × | 3 | ✓ | ✓ | single exact | 5% | identical | 10% |
| ✓ | 3 | ✓ | ✓ | single exact | 5% | identical | 0% |
| × | 1 | × | × | single w/ error | 5% | identical | 0% |
| ✓ | 1 | × | × | single w/ error | 5% | identical | 0% |
| × | 2 | × | × | single w/ error | 5% | identical | 10% |
| ✓ | 2 | × | × | single w/ error | 5% | identical | 10% |
| × | 3 | × | × | single w/ error | 5% | identical | 10% |
| ✓ | 3 | × | × | single w/ error | 5% | identical | 10% |
| × | 2 | ✓ | × | single w/ error | 5% | identical | 0% |
| ✓ | 2 | ✓ | × | single w/ error | 5% | identical | 0% |
| × | 3 | ✓ | × | single w/ error | 5% | identical | 0% |
| ✓ | 3 | ✓ | × | single w/ error | 5% | identical | 0% |
| × | 1 | × | ✓ | single w/ error | 5% | identical | 20% |
| ✓ | 1 | × | ✓ | single w/ error | 5% | identical | 10% |
| × | 2 | × | ✓ | single w/ error | 5% | identical | 10% |
| ✓ | 2 | × | ✓ | single w/ error | 5% | identical | 0% |
| × | 3 | × | ✓ | single w/ error | 5% | identical | 10% |
| ✓ | 3 | × | ✓ | single w/ error | 5% | identical | 0% |
| × | 2 | ✓ | ✓ | single w/ error | 5% | identical | 20% |
| ✓ | 2 | ✓ | ✓ | single w/ error | 5% | identical | 0% |
| × | 3 | ✓ | ✓ | single w/ error | 5% | identical | 10% |
| ✓ | 3 | ✓ | ✓ | single w/ error | 5% | identical | 0% |
| × | 1 | × | × | mult. w/ error | 5% | identical | 0% |
| ✓ | 1 | × | × | mult. w/ error | 5% | identical | 10% |
| × | 2 | × | × | mult. w/ error | 5% | identical | 0% |
| ✓ | 2 | × | × | mult. w/ error | 5% | identical | 0% |
| × | 3 | × | × | mult. w/ error | 5% | identical | 0% |
| ✓ | 3 | × | × | mult. w/ error | 5% | identical | 10% |
| × | 2 | ✓ | × | mult. w/ error | 5% | identical | 0% |
| ✓ | 2 | ✓ | × | mult. w/ error | 5% | identical | 0% |
| × | 3 | ✓ | × | mult. w/ error | 5% | identical | 10% |
| ✓ | 3 | ✓ | × | mult. w/ error | 5% | identical | 0% |
| × | 1 | × | ✓ | mult. w/ error | 5% | identical | 0% |
| ✓ | 1 | × | ✓ | mult. w/ error | 5% | identical | 0% |
| × | 2 | × | ✓ | mult. w/ error | 5% | identical | 0% |
| ✓ | 2 | × | ✓ | mult. w/ error | 5% | identical | 0% |
| × | 3 | × | ✓ | mult. w/ error | 5% | identical | 0% |
| ✓ | 3 | × | ✓ | mult. w/ error | 5% | identical | 0% |
| × | 2 | ✓ | ✓ | mult. w/ error | 5% | identical | 0% |
| ✓ | 2 | ✓ | ✓ | mult. w/ error | 5% | identical | 0% |
| × | 3 | ✓ | ✓ | mult. w/ error | 5% | identical | 10% |
| ✓ | 3 | ✓ | ✓ | mult. w/ error | 5% | identical | 0% |
| × | 1 | × | × | single exact | 25% | identical | 0% |
| ✓ | 1 | × | × | single exact | 25% | identical | 0% |
| × | 2 | × | × | single exact | 25% | identical | 10% |
| ✓ | 2 | × | × | single exact | 25% | identical | 0% |
| × | 3 | × | × | single exact | 25% | identical | 0% |

| ultrametric phylogenies? | number of phenotypes | phenotypic correlations? | evolutionary trends? | measurement type | proportion missing data | assumed parameters | proportion $p < 0.05$ |
|---|---|---|---|---|---|---|---|
| ✓ | 3 | × | × | single exact | 25% | identical | 0% |
| × | 2 | ✓ | × | single exact | 25% | identical | 0% |
| ✓ | 2 | ✓ | × | single exact | 25% | identical | 20% |
| × | 3 | ✓ | × | single exact | 25% | identical | 10% |
| ✓ | 3 | ✓ | × | single exact | 25% | identical | 0% |
| × | 1 | × | ✓ | single exact | 25% | identical | 10% |
| ✓ | 1 | × | ✓ | single exact | 25% | identical | 0% |
| × | 2 | × | ✓ | single exact | 25% | identical | 0% |
| ✓ | 2 | × | ✓ | single exact | 25% | identical | 10% |
| × | 3 | × | ✓ | single exact | 25% | identical | 0% |
| ✓ | 3 | × | ✓ | single exact | 25% | identical | 0% |
| × | 2 | ✓ | ✓ | single exact | 25% | identical | 0% |
| ✓ | 2 | ✓ | ✓ | single exact | 25% | identical | 10% |
| × | 3 | ✓ | ✓ | single exact | 25% | identical | 10% |
| ✓ | 3 | ✓ | ✓ | single exact | 25% | identical | 0% |
| × | 1 | × | × | single w/ error | 25% | identical | 0% |
| ✓ | 1 | × | × | single w/ error | 25% | identical | 0% |
| × | 2 | × | × | single w/ error | 25% | identical | 0% |
| ✓ | 2 | × | × | single w/ error | 25% | identical | 0% |
| × | 3 | × | × | single w/ error | 25% | identical | 0% |
| ✓ | 3 | × | × | single w/ error | 25% | identical | 0% |
| × | 2 | ✓ | × | single w/ error | 25% | identical | 0% |
| ✓ | 2 | ✓ | × | single w/ error | 25% | identical | 20% |
| × | 3 | ✓ | × | single w/ error | 25% | identical | 0% |
| ✓ | 3 | ✓ | × | single w/ error | 25% | identical | 0% |
| × | 1 | × | ✓ | single w/ error | 25% | identical | 0% |
| ✓ | 1 | × | ✓ | single w/ error | 25% | identical | 0% |
| × | 2 | × | ✓ | single w/ error | 25% | identical | 0% |
| ✓ | 2 | × | ✓ | single w/ error | 25% | identical | 10% |
| × | 3 | × | ✓ | single w/ error | 25% | identical | 0% |
| ✓ | 3 | × | ✓ | single w/ error | 25% | identical | 20% |
| × | 2 | ✓ | ✓ | single w/ error | 25% | identical | 0% |
| ✓ | 2 | ✓ | ✓ | single w/ error | 25% | identical | 10% |
| × | 3 | ✓ | ✓ | single w/ error | 25% | identical | 10% |
| ✓ | 3 | ✓ | ✓ | single w/ error | 25% | identical | 0% |
| × | 1 | × | × | mult. w/ error | 25% | identical | 0% |
| ✓ | 1 | × | × | mult. w/ error | 25% | identical | 0% |
| × | 2 | × | × | mult. w/ error | 25% | identical | 30% |
| ✓ | 2 | × | × | mult. w/ error | 25% | identical | 0% |
| × | 3 | × | × | mult. w/ error | 25% | identical | 0% |
| ✓ | 3 | × | × | mult. w/ error | 25% | identical | 0% |
| × | 2 | ✓ | × | mult. w/ error | 25% | identical | 0% |
| ✓ | 2 | ✓ | × | mult. w/ error | 25% | identical | 0% |
| × | 3 | ✓ | × | mult. w/ error | 25% | identical | 10% |
| ✓ | 3 | ✓ | × | mult. w/ error | 25% | identical | 0% |
| × | 1 | × | ✓ | mult. w/ error | 25% | identical | 0% |
| ✓ | 1 | × | ✓ | mult. w/ error | 25% | identical | 10% |
| × | 2 | × | ✓ | mult. w/ error | 25% | identical | 0% |
| ✓ | 2 | × | ✓ | mult. w/ error | 25% | identical | 20% |
| × | 3 | × | ✓ | mult. w/ error | 25% | identical | 10% |
| ✓ | 3 | × | ✓ | mult. w/ error | 25% | identical | 0% |
| × | 2 | ✓ | ✓ | mult. w/ error | 25% | identical | 20% |
| ✓ | 2 | ✓ | ✓ | mult. w/ error | 25% | identical | 0% |
| × | 3 | ✓ | ✓ | mult. w/ error | 25% | identical | 10% |
| ✓ | 3 | ✓ | ✓ | mult. w/ error | 25% | identical | 0% |
| × | 1 | × | × | single exact | 0% | random | 0% |
| ✓ | 1 | × | × | single exact | 0% | random | 10% |
| × | 2 | × | × | single exact | 0% | random | 10% |
| ✓ | 2 | × | × | single exact | 0% | random | 0% |
| × | 3 | × | × | single exact | 0% | random | 0% |
| ✓ | 3 | × | × | single exact | 0% | random | 0% |
| × | 2 | ✓ | × | single exact | 0% | random | 0% |
| ✓ | 2 | ✓ | × | single exact | 0% | random | 10% |
| × | 3 | ✓ | × | single exact | 0% | random | 20% |
| ✓ | 3 | ✓ | × | single exact | 0% | random | 0% |
| × | 1 | × | ✓ | single exact | 0% | random | 0% |
| ✓ | 1 | × | ✓ | single exact | 0% | random | 0% |
| × | 2 | × | ✓ | single exact | 0% | random | 0% |
| ✓ | 2 | × | ✓ | single exact | 0% | random | 0% |
| × | 3 | × | ✓ | single exact | 0% | random | 0% |
| ✓ | 3 | × | ✓ | single exact | 0% | random | 10% |
| × | 2 | ✓ | ✓ | single exact | 0% | random | 0% |
| ✓ | 2 | ✓ | ✓ | single exact | 0% | random | 0% |

| ultrametric phylogenies? | number of phenotypes | phenotypic correlations? | evolutionary trends? | measurement type | proportion missing data | assumed parameters | proportion $p < 0.05$ |
|---|---|---|---|---|---|---|---|
| × | 3 | ✓ | ✓ | single exact | 0% | random | 20% |
| ✓ | 3 | ✓ | ✓ | single exact | 0% | random | 0% |
| × | 1 | × | × | single w/ error | 0% | random | 0% |
| ✓ | 1 | × | × | single w/ error | 0% | random | 10% |
| × | 2 | × | × | single w/ error | 0% | random | 10% |
| ✓ | 2 | × | × | single w/ error | 0% | random | 0% |
| × | 3 | × | × | single w/ error | 0% | random | 10% |
| ✓ | 3 | × | × | single w/ error | 0% | random | 0% |
| × | 2 | ✓ | × | single w/ error | 0% | random | 10% |
| ✓ | 2 | ✓ | × | single w/ error | 0% | random | 10% |
| × | 3 | ✓ | × | single w/ error | 0% | random | 0% |
| ✓ | 3 | ✓ | × | single w/ error | 0% | random | 10% |
| × | 1 | × | ✓ | single w/ error | 0% | random | 0% |
| ✓ | 1 | × | ✓ | single w/ error | 0% | random | 10% |
| × | 2 | × | ✓ | single w/ error | 0% | random | 0% |
| ✓ | 2 | × | ✓ | single w/ error | 0% | random | 0% |
| × | 3 | × | ✓ | single w/ error | 0% | random | 0% |
| ✓ | 3 | × | ✓ | single w/ error | 0% | random | 0% |
| × | 2 | ✓ | ✓ | single w/ error | 0% | random | 20% |
| ✓ | 2 | ✓ | ✓ | single w/ error | 0% | random | 10% |
| × | 3 | ✓ | ✓ | single w/ error | 0% | random | 20% |
| ✓ | 3 | ✓ | ✓ | single w/ error | 0% | random | 0% |
| × | 1 | × | × | mult. w/ error | 0% | random | 0% |
| ✓ | 1 | × | × | mult. w/ error | 0% | random | 10% |
| × | 2 | × | × | mult. w/ error | 0% | random | 0% |
| ✓ | 2 | × | × | mult. w/ error | 0% | random | 20% |
| × | 3 | × | × | mult. w/ error | 0% | random | 10% |
| ✓ | 3 | × | × | mult. w/ error | 0% | random | 0% |
| × | 2 | ✓ | × | mult. w/ error | 0% | random | 0% |
| ✓ | 2 | ✓ | × | mult. w/ error | 0% | random | 10% |
| × | 3 | ✓ | × | mult. w/ error | 0% | random | 10% |
| ✓ | 3 | ✓ | × | mult. w/ error | 0% | random | 10% |
| × | 1 | × | ✓ | mult. w/ error | 0% | random | 0% |
| ✓ | 1 | × | ✓ | mult. w/ error | 0% | random | 0% |
| × | 2 | × | ✓ | mult. w/ error | 0% | random | 10% |
| ✓ | 2 | × | ✓ | mult. w/ error | 0% | random | 20% |
| × | 3 | × | ✓ | mult. w/ error | 0% | random | 0% |
| ✓ | 3 | × | ✓ | mult. w/ error | 0% | random | 10% |
| × | 2 | ✓ | ✓ | mult. w/ error | 0% | random | 0% |
| ✓ | 2 | ✓ | ✓ | mult. w/ error | 0% | random | 20% |
| × | 3 | ✓ | ✓ | mult. w/ error | 0% | random | 0% |
| ✓ | 3 | ✓ | ✓ | mult. w/ error | 0% | random | 10% |
| × | 1 | × | × | single exact | 5% | random | 0% |
| ✓ | 1 | × | × | single exact | 5% | random | 20% |
| × | 2 | × | × | single exact | 5% | random | 0% |
| ✓ | 2 | × | × | single exact | 5% | random | 0% |
| × | 3 | × | × | single exact | 5% | random | 0% |
| ✓ | 3 | × | × | single exact | 5% | random | 10% |
| × | 2 | ✓ | × | single exact | 5% | random | 0% |
| ✓ | 2 | ✓ | × | single exact | 5% | random | 10% |
| × | 3 | ✓ | × | single exact | 5% | random | 0% |
| ✓ | 3 | ✓ | × | single exact | 5% | random | 0% |
| × | 1 | × | ✓ | single exact | 5% | random | 10% |
| ✓ | 1 | × | ✓ | single exact | 5% | random | 10% |
| × | 2 | × | ✓ | single exact | 5% | random | 10% |
| ✓ | 2 | × | ✓ | single exact | 5% | random | 10% |
| × | 3 | × | ✓ | single exact | 5% | random | 0% |
| ✓ | 3 | × | ✓ | single exact | 5% | random | 0% |
| × | 2 | ✓ | ✓ | single exact | 5% | random | 10% |
| ✓ | 2 | ✓ | ✓ | single exact | 5% | random | 0% |
| × | 3 | ✓ | ✓ | single exact | 5% | random | 0% |
| ✓ | 3 | ✓ | ✓ | single exact | 5% | random | 0% |
| × | 1 | × | × | single w/ error | 5% | random | 0% |
| ✓ | 1 | × | × | single w/ error | 5% | random | 10% |
| × | 2 | × | × | single w/ error | 5% | random | 10% |
| ✓ | 2 | × | × | single w/ error | 5% | random | 0% |
| × | 3 | × | × | single w/ error | 5% | random | 10% |
| ✓ | 3 | × | × | single w/ error | 5% | random | 10% |
| × | 2 | ✓ | × | single w/ error | 5% | random | 10% |
| ✓ | 2 | ✓ | × | single w/ error | 5% | random | 10% |
| × | 3 | ✓ | × | single w/ error | 5% | random | 0% |
| ✓ | 3 | ✓ | × | single w/ error | 5% | random | 10% |
| × | 1 | × | ✓ | single w/ error | 5% | random | 20% |

| ultrametric phylogenies? | number of phenotypes | phenotypic correlations? | evolutionary trends? | measurement type | proportion missing data | assumed parameters | proportion $p < 0.05$ |
|---|---|---|---|---|---|---|---|
| ✓ | 1 | × | ✓ | single w/ error | 5% | random | 10% |
| × | 2 | × | ✓ | single w/ error | 5% | random | 10% |
| ✓ | 2 | × | ✓ | single w/ error | 5% | random | 0% |
| × | 3 | × | ✓ | single w/ error | 5% | random | 0% |
| ✓ | 3 | × | ✓ | single w/ error | 5% | random | 0% |
| × | 2 | ✓ | ✓ | single w/ error | 5% | random | 0% |
| ✓ | 2 | ✓ | ✓ | single w/ error | 5% | random | 0% |
| × | 3 | ✓ | ✓ | single w/ error | 5% | random | 0% |
| ✓ | 3 | ✓ | ✓ | single w/ error | 5% | random | 0% |
| × | 1 | × | × | mult. w/ error | 5% | random | 0% |
| ✓ | 1 | × | × | mult. w/ error | 5% | random | 20% |
| × | 2 | × | × | mult. w/ error | 5% | random | 0% |
| ✓ | 2 | × | × | mult. w/ error | 5% | random | 20% |
| × | 3 | × | × | mult. w/ error | 5% | random | 0% |
| ✓ | 3 | × | × | mult. w/ error | 5% | random | 0% |
| × | 2 | ✓ | × | mult. w/ error | 5% | random | 10% |
| ✓ | 2 | ✓ | × | mult. w/ error | 5% | random | 20% |
| × | 3 | ✓ | × | mult. w/ error | 5% | random | 0% |
| ✓ | 3 | ✓ | × | mult. w/ error | 5% | random | 10% |
| × | 1 | × | ✓ | mult. w/ error | 5% | random | 0% |
| ✓ | 1 | × | ✓ | mult. w/ error | 5% | random | 0% |
| × | 2 | × | ✓ | mult. w/ error | 5% | random | 10% |
| ✓ | 2 | × | ✓ | mult. w/ error | 5% | random | 10% |
| × | 3 | × | ✓ | mult. w/ error | 5% | random | 0% |
| ✓ | 3 | × | ✓ | mult. w/ error | 5% | random | 10% |
| × | 2 | ✓ | ✓ | mult. w/ error | 5% | random | 10% |
| ✓ | 2 | ✓ | ✓ | mult. w/ error | 5% | random | 10% |
| × | 3 | ✓ | ✓ | mult. w/ error | 5% | random | 0% |
| ✓ | 3 | ✓ | ✓ | mult. w/ error | 5% | random | 0% |
| × | 1 | × | × | single exact | 25% | random | 10% |
| ✓ | 1 | × | × | single exact | 25% | random | 20% |
| × | 2 | × | × | single exact | 25% | random | 10% |
| ✓ | 2 | × | × | single exact | 25% | random | 0% |
| × | 3 | × | × | single exact | 25% | random | 0% |
| ✓ | 3 | × | × | single exact | 25% | random | 0% |
| × | 2 | ✓ | × | single exact | 25% | random | 0% |
| ✓ | 2 | ✓ | × | single exact | 25% | random | 0% |
| × | 3 | ✓ | × | single exact | 25% | random | 10% |
| ✓ | 3 | ✓ | × | single exact | 25% | random | 20% |
| × | 1 | × | ✓ | single exact | 25% | random | 10% |
| ✓ | 1 | × | ✓ | single exact | 25% | random | 0% |
| × | 2 | × | ✓ | single exact | 25% | random | 10% |
| ✓ | 2 | × | ✓ | single exact | 25% | random | 0% |
| × | 3 | × | ✓ | single exact | 25% | random | 0% |
| ✓ | 3 | × | ✓ | single exact | 25% | random | 10% |
| × | 2 | ✓ | ✓ | single exact | 25% | random | 20% |
| ✓ | 2 | ✓ | ✓ | single exact | 25% | random | 10% |
| × | 3 | ✓ | ✓ | single exact | 25% | random | 0% |
| ✓ | 3 | ✓ | ✓ | single exact | 25% | random | 0% |
| × | 1 | × | × | single w/ error | 25% | random | 0% |
| ✓ | 1 | × | × | single w/ error | 25% | random | 10% |
| × | 2 | × | × | single w/ error | 25% | random | 0% |
| ✓ | 2 | × | × | single w/ error | 25% | random | 0% |
| × | 3 | × | × | single w/ error | 25% | random | 0% |
| ✓ | 3 | × | × | single w/ error | 25% | random | 0% |
| × | 2 | ✓ | × | single w/ error | 25% | random | 0% |
| ✓ | 2 | ✓ | × | single w/ error | 25% | random | 0% |
| × | 3 | ✓ | × | single w/ error | 25% | random | 10% |
| ✓ | 3 | ✓ | × | single w/ error | 25% | random | 0% |
| × | 1 | × | ✓ | single w/ error | 25% | random | 20% |
| ✓ | 1 | × | ✓ | single w/ error | 25% | random | 0% |
| × | 2 | × | ✓ | single w/ error | 25% | random | 10% |
| ✓ | 2 | × | ✓ | single w/ error | 25% | random | 0% |
| × | 3 | × | ✓ | single w/ error | 25% | random | 0% |
| ✓ | 3 | × | ✓ | single w/ error | 25% | random | 0% |
| × | 2 | ✓ | ✓ | single w/ error | 25% | random | 10% |
| ✓ | 2 | ✓ | ✓ | single w/ error | 25% | random | 0% |
| × | 3 | ✓ | ✓ | single w/ error | 25% | random | 0% |
| ✓ | 3 | ✓ | ✓ | single w/ error | 25% | random | 50% |
| × | 1 | × | × | mult. w/ error | 25% | random | 10% |
| ✓ | 1 | × | × | mult. w/ error | 25% | random | 0% |
| × | 2 | × | × | mult. w/ error | 25% | random | 10% |
| ✓ | 2 | × | × | mult. w/ error | 25% | random | 0% |

| ultrametric phylogenies? | number of phenotypes | phenotypic correlations? | evolutionary trends? | measurement type | proportion missing data | assumed parameters | proportion $p < 0.05$ |
|---|---|---|---|---|---|---|---|
| × | 3 | × | × | mult. w/ error | 25% | random | 0% |
| ✓ | 3 | × | × | mult. w/ error | 25% | random | 0% |
| × | 2 | ✓ | × | mult. w/ error | 25% | random | 10% |
| ✓ | 2 | ✓ | × | mult. w/ error | 25% | random | 20% |
| × | 3 | ✓ | × | mult. w/ error | 25% | random | 10% |
| ✓ | 3 | ✓ | × | mult. w/ error | 25% | random | 0% |
| × | 1 | × | ✓ | mult. w/ error | 25% | random | 10% |
| ✓ | 1 | × | ✓ | mult. w/ error | 25% | random | 0% |
| × | 2 | × | ✓ | mult. w/ error | 25% | random | 0% |
| ✓ | 2 | × | ✓ | mult. w/ error | 25% | random | 20% |
| × | 3 | × | ✓ | mult. w/ error | 25% | random | 20% |
| ✓ | 3 | × | ✓ | mult. w/ error | 25% | random | 0% |
| × | 2 | ✓ | ✓ | mult. w/ error | 25% | random | 10% |
| ✓ | 2 | ✓ | ✓ | mult. w/ error | 25% | random | 20% |
| × | 3 | ✓ | ✓ | mult. w/ error | 25% | random | 0% |
| ✓ | 3 | ✓ | ✓ | mult. w/ error | 25% | random | 20% |

#### *Comparing multivariate samples/distributions with Mahalanobis distances*

To actually check whether sampled continuous stochastic character maps follow their expected conditional multivariate normal distributions given by $\overline{M_p}$ and $\overline{\Sigma_{pp}}$ in Eq. (30), we calculated the Mahalanobis distance for each sampled map based on their expected distribution. Theoretically, these distances should follow a simple $\chi^2$ distribution with degrees of freedom equal to the dimensionality of the conditional distribution (Petersen and Pedersen, 2012). To determine whether the observed distributions of Mahalanobis distances significantly differed from their $\chi^2$ expectations, we used R's 1-sample Kolmogorov-Smirnov test, which should yield a uniform distribution of p-values if our algorithm works as intended (Marsaglia et al., 2003).

### Pipeline and Simulation Study

#### *Marginalizing over root value parameters*

Our implementation for fitting factor-dependent Brownian Motion models does not estimate additional parameters for the trait value at the root of a phylogeny, unlike many other phylogenetic comparative methods for modeling continuous trait evolution (e.g., Revell, 2012; Pennell et al., 2014; Boucher et al., 2018). Instead, our implementation calculates conditional likelihoods for each continuous stochastic character map while marginalizing over all possible root values assuming either a flat or nuisance prior (the latter is equivalent to the "Fitzjohn root prior" used in discrete trait evolution models, see FitzJohn et al., 2009). We take this approach because different continuous stochastic character maps often imply conflicting root values under a given model (see Figure 4 in Boyko et al., 2023 for an analogous phenomenon). Notably, we used a flat prior on root values for all analyses presented in the main paper, as preliminary simulation experiments and empirical analyses suggested that using nuisance priors instead only had a minor impact on parameter estimates and associated likelihoods. Nonetheless, we retained the option to use nuisance priors on root values in our implementation in case users wish to experiment with this setting in the future. We hope to more systematically study the impact of alternative root value priors in the future.

To actually calculate likelihoods while marginalizing over root values, note that the likelihood of observed data under any (potentially multivariate) Brownian Motion model conditional on a vector of root values $x$ is given by $r\Phi(x; \hat{x}, V)$, where $r$ is a proportionality constant (called the "remainder" in Hassler et al., 2022), $\hat{x}$ and $V$ denote the expected root values and associated uncertainty (in the form of a variance-covariance matrix), and $\Phi(x; \mu, \Sigma)$ represents the probability density function of a multivariate normal distribution with mean $\mu$ and variance-covariance matrix $\Sigma$ evaluated at $x$. Because the integral of any probability density function is 1 by definition, the integral of this expression is $r$—thus, the overall likelihood for a given continuous stochastic character map is $r$ under a flat prior. Under a nuisance prior, the likelihood is equal to the proportionality constant $r$ multiplied by the integral of the squared multivariate normal probability density function ($r/\sqrt{|V|(4\pi)^k}$, where $k$ denotes the number of traits/dimensions of the multivariate normal distribution). Thus, assuming a nuisance prior will deflate likelihoods conditional on any given continuous stochastic character map when root values are highly uncertain (i.e., $|V|$ is large) and vice versa.

#### *Simulating noisy factor-rate relationships*

To simulate noise around factor-rate relationships, we multiplied rates of trait $(Y)$ evolution at each time point with a random variable sampled from a gamma distribution with shape and rate $\frac{dt}{\nu}$, where $dt$ is the length of the time interval preceding a time point. These multipliers represent the average value of a white noise gamma process with a mean of 1 and unit variance $\nu$ over a time interval of length $dt$. This model of rate change has been used in uncorrelated relaxed clock models in previous work (Lepage et al., 2007; Lartillot et al., 2016) and has the advantage of defining a stochastic rate change process in continuous time. More common uncorrelated relaxed clock models, which assign independent rate scalars to entire branches, do not yield a coherent process of rate change that can be simulated over fine-grained time points (Lepage et al., 2007). Additionally, this rate change process is also identical to the expected rate fluctuations under a variance-gamma process (a type of Lévy or "pulsed" trait evolution process explored in some prior work; see Landis et al., 2013; Landis and Schraiber, 2017). Importantly, this procedure ensures random noise around rates tends to "average out" over sufficiently long periods of time, such that rates converge to what would be expected under the simulated factor-rate relationship given enough data. Thus, this uncorrelated rate noise—in contrast to autocorrelated noise—only weakens simulated factor-rate relationships rather than completely altering them. For our simulations, we set $\nu$ to 0.05, corresponding to rates ranging between roughly 10%-300% their expected value over a time interval of 0.1, 40%-200% for an interval of 0.3, and 60%-150% for an interval of 1 (keep in mind we rescaled all phylogenies to have a height of 1 for our simulation study).

#### *Averaging conditional likelihoods in the face of outliers*

In general, we found likelihood surfaces under continuous factor-dependent Brownian Motion models to be quite complex, exhibiting multiple optima and/or relatively flat "ridges" that present challenges for numerical optimization. Such issues seem partially due to necessarily finite sample of continuous stochastic character maps underlying these models, as it is common for multiple parts of parameter space to yield relatively high overall likelihoods based on extremely high, “outlier” conditional likelihoods for just one to a few maps. This phenomenon—which we dubbed “optimizing for a single map”—tends to result in inflated likelihoods and unrealistic parameter estimates (e.g., extremely strong and narrow threshold/sweetspot relationships). We experimented with various techniques for weighting and/or removing outlier conditional likelihoods to prevent such behavior, ultimately finding that Winsorization—that is, rounding outliers to particular quantiles of the conditional likelihood distribution across all maps—seemed to combat the issue most effectively. Our current implementation for fitting factor-dependent Brownian Motion models uses two-sided, percentile-based Winsorization, such that 5% Winsorization corresponds to rounding extremely low and high conditional likelihoods to the empirical 2.5 and 97.5% quantiles, respectively, of the conditional likelihood distribution (Liao et al., 2017).

To select the optimal number of continuous stochastic character maps (i.e., “map sample size”) and level of Winsorization to use in our simulation study, we performed a preliminary version of our simulation study with only 30 replicates per condition instead of 120. For each simulated dataset, we fit all models described in the main text based on either 50, 100, or 200 continuous stochastic character maps while Winsorizing either the top 0, 2, or 4 outlier conditional likelihoods. We generally analyzed these results by calculating differences in overall likelihoods among identical models fit to the same simulated datasets but with different map sample sizes/Winsorization levels. Note that we were not yet sure at this point if it was more convenient to Winsorize based on percentiles or counts of conditional likelihoods. Our final decision to use percentile-based Winsorization was informed by the results of these preliminary simulation experiments, which revealed that the effect of Winsorization on likelihoods tended to be more consistent based on percentiles rather than counts. For example, compared to their non-Winsorized counterparts, models based on 100 maps with 2% Winsorization (i.e., 2 conditional likelihoods) decreased likelihoods by an amount similar to those based on 200 maps with 2% Winsorization (i.e., 4 conditonal likelihoods). By comparison, models based on 200 maps with 1% Winsorization (i.e., 2 conditional likelihoods) yielded smaller decreases in likelihoods.

Generally speaking, larger map sample sizes tended to increase overall likelihoods of fitted models, while Winsorization tended to decrease likelihoods. These results are unsurprising, as they simply suggest that basing models on additional continuous stochastic character maps increases the chances of finding a "good" map which is particularly likely to explain the observed data. However, the apparent magnitude of these trends varied quite dramatically between observed and dummy-factor dependent models. For observed factor-dependent models, any level of Winsorization decreased log likelihoods by about 0.1–0.2 on average. Beyond this, however, the exact map sample size/Winsorization level used did not matter much, amounting to average log likelihood changes of $\sim$0.05 at most. By comparison, map sample size and especially Winsorization level had dramatic effects on likelihoods of dummy factor-dependent models. While map sample size still had a fairly modest effect on likelihoods for dummy factor-dependent models (doubling map sample size was associated with average log likelihood increases of about 0.5), higher Winsorization levels had fairly profound effects, commonly decreasing log likelihoods by several units. Notably, however, this effect got weaker at higher and higher Winsorization levels (e.g., the log likelihood decrease between models with 4 and 0% Winsorization averaged around 3, whereas the decrease between models with 8 and 4% Winsorization averaged around 1). We attribute the divergent patterns between observed and dummy factor-dependent models to the fact that the space of all possible evolutionary histories on a phylogeny is inevitably smaller if conditioned on a set of observed measurements (i.e., as in the observed factor histories). Consequently, maps of observed factor histories will generally be more "redundant" with one another compared to maps of dummy factor histories, and adding additional maps and/or reducing the influence of particular maps will therefore lead to more pronounced effects on likelihoods for dummy factor-dependent models compared to observed factor-dependent models.

While larger map sample sizes may benefit inference under our pipeline by more adequately representing the possible evolutionary histories of a given factor, our preliminary simulation experiment nonetheless suggests such effects are relatively minor. Thus, we decided to base all models in our final simulation study on samples of 100 continuous stochastic character maps. In addition to rendering the simulation study more computationally manageable, this better reflects typical sample sizes used by macroevoltuionary researchers in analyses based on discrete stochastic character maps. As for Winsorization, we ultimately decided to fit dummy factor-dependent models with either 0 or 2% Winsorization and observed factor-dependent models with either 0 or 5%. We used lower Winsorization levels for dummy factor-dependent models based on the increased sensitivity of their overall likelihoods to Winsorization per our preliminary simulation experiments.

#### *Optimization settings*

We fit all models to simulated data by running NLOPT's (Johnson, 2021; Ypma et al., 2022) SBPLX (Rowan, 1990) algorithm 10 times from initial parameter values sampled from a uniform distribution spanning from -5 to 5, taking inferred parameter values from whichever optimization run found the highest likelihood. To save on the time needed to run the simulation study, we limited each optimization run to a maximum of 10,000 iterations. To prevent optimization algorithms from spending too much exploring these trivial regions of parameter space (e.g., arbitrarily small/large location/$\theta$ and/or width/$\omega$ parameters leading to effectively constant rate models), we imposed boundaries on parameters by defining a "factor interval" spanning from $\min X - (\max X - \min X)/2$ to $\max X + (\max X - \min X)/2$, where $X$ represents whatever factor a given parameter function depends on (either the observed factor $X_o$ or simulated dummy factor $D$). We constrained the location of inferred shifts and peaks/dips to lie within the factor interval and the corresponding widths (given by $e^{\omega}$) to be between $1/100^{\text{th}}$ and 5 times the range of the factor interval. We also limited the rate deviation ($\delta$) parameter to be between -10 and 10, as rate deviations with an absolute value of 10 or greater cause minimum and maximum rates to be numerically indistinguishable from 0 and $2e^{\alpha}$ (i.e., the maximum allowable rate for a given mid rate or $\alpha$ value), respectively.

As described in the previous section, we fit all models based on samples of 100 continuous stochastic character maps with 0/2% Winsorization and 0/5% Winsorization for dummy and observed factor-dependent models, respectively. Ultimately, we decided to report results in the main text based on dummy and observed factor-dependent models with 0% and 5% Winsorization, respectively. We made this decision for two main reasons. First, we found that—relative to non-Winsorized models—even 2% Winsorization reduced the likelihoods of dummy factor-dependent models enough to substantially increase the overall error rates of our pipeline (see following section for how we define error rates in this study). This results from maps of dummy factors generally exhibiting divergent histories that often support conflicting inferences. Further, the goal of dummy factor-dependent models is to find just a few continuous stochastic character maps associated with high conditional likelihoods—exactly what Winsorization is meant to prevent in this context. Second, while Winsorization of observed factor-dependent models barely affected overall model selection results (e.g., 0% and 5% Winsorization resulted in an average error rates of 6% and 5%, respectively), we nonetheless found that Winsorization slightly improved parameter estimates under observed factor-dependent models. Without Winsorization, these models were prone to inferring extreme rate deviation and width parameters which were unrealistic given our simulation conditions, often bumping up against the parameter boundaries described in the previous paragraph. Such patterns suggest non-Winsorized observed factor-dependent models were more likely to optimize threshold/sweetspot relationships based on just one to a few stochastic character maps (i.e., "optimizing for a single map"; see previous section) and that Winsorization generally improves the robustness of parameter inferences under observed factor-dependent models.

#### *Model selection and model-averaged predictions of factor-rate relationships*

We assessed the behavior of our proposed pipeline for detecting associations between continuously-varying factors and rates of continuous trait evolution through both model selection and parameter inference-based approaches to interpreting model results.

For model selection, we used small sample size corrected Akaike Information Criteria (AICc) because it is convenient and standard practice among modern phylogenetic comparative studies. Preliminary results suggested that, even when considering dummy factor ($D$)-dependent models, simulated hidden factor ($X_h$)-rate relationships and/or noisy rates still occasionally yielded spurious support for particular observed factor ($X_o$)-dependent models (usually threshold and especially sweetspot relationship-based models). On the other hand, simulated $X_o$-rate relationships typically resulted in more broadly increased support across all $X_o$-dependent models (albeit to differing degrees based on the specific $X_o$-rate relationship used to simulate data). Thus, we obtained the best model selection results by summarizing patterns of support across multiple models using sums of AICc weights rather than directly focusing on pairwise differences in AICc. To select a single best-fitting model for each simulated dataset, we chose whichever $X_o$-dependent model exhibited the lowest AICc *only* if the sum of AICc weights across all $X_o$-dependent models exceeded a given cutoff (our results suggest 0.7–0.9 works well in practice), and selecting whichever null (i.e., constant rate or $D$-dependent) model exhibited the lowest AICc otherwise. To summarize our model selection results, we assessed how error (i.e., "false positives", the percent of times an $X_o$-dependent model is selected as the best-fitting model for simulated data with constant/$X_h$-dependent rates) and power (i.e., "true positives", the percent of times an $X_o$-dependent model is selected as the best-fitting model for simulated data with $X_o$-dependent rates) vary across different simulation conditions. Additionally, among the simulations correctly yielding support for $X_o$-rate relationships, we calculated "differentiation rates" as the percent of times for which the best-fitting model assumed the same $X_o$-rate relationship used to simulate data.

For the parameter inference-based approach, we used AICc weights to calculate model-averaged predictions of $X_o$-rate relationships for all simulations with either constant or $X_o$-dependent rates, ignoring simulations with $X_h$-dependent rates because they lack a straightforward expected $X_o$-rate relationship with which to compare predictions. Importantly, we ignored AICc weights for $D$-dependent models when calculating model-averaged predictions because such models make no direct inferences regarding the relationship between rates and $X_o$. We measured the overall quality of model-averaged rate predictions across different $X_o$ values under each simulation condition by calculating the percent of predicted rates greater than simulated rates (bias), the fold difference between the 2.5% and 97.5% quantiles of predicted rates across all replicates (precision), and the median absolute difference between predicted and simulated rates on a log scale (accuracy; multiplied by -1 such that higher values correspond to greater accuracy).

### Empirical Example

#### *Phylogenetic inference*

To develop a phylogenetic framework for eucalypts, we used a recently-published dataset of 101 nuclear loci alignments ranging from 225 to 6555 (average 1281) base pairs long (Crisp et al., 2024). We used IQ-Tree 2 (Minh et al., 2020) to select the substitution models and infer maximum likelihood phylogenies for each loci, then used wASTRAL (Zhang and Mirarab, 2022) and weighted TREE-QMC (Han and Molloy, 2025) to infer species trees based on the resulting gene trees. The species tree topologies inferred under both methods were highly similar, though the TREE-QMC topology exhibits a few minor differences that better align with taxonomic consensus (e.g., monophyly of species

*Eucalyptus behriana* and section Latoangulatae). Thus, we based downstream analyses on the TREE-QMC topology. We dated the phylogeny using branch-length based methods (e.g., Smith and O'Meara, 2012; Mai and Mirarab, 2021; Mai et al., 2024) in conjunction with recently-developed tools for estimating branch lengths while explicitly accounting for phylogenetic discordance (e.g., Tabatabaee et al., 2023; Arasti et al., 2024). This allowed us to more fully utilize data across all loci compared to more conventional dating approaches for phylogenetic data exhibiting high levels of discordance, as is the case here. We used CASTLES (Tabatabaee et al., 2023) along with topology-constrained metric matching (TCMM; Arasti et al., 2024) to estimate branch lengths for our species tree and wLogDate (Mai and Mirarab, 2021) to date the resulting phylogeny.

We initially inferred branch lengths for our species tree via CASTLES based on a filtered set of 45 out of 101 gene trees which exhibited both relatively high branch support (at least 70% of branches with approximate Bayes test/aBayes support greater than the minimum 1/3; see Anisimova et al., 2011) and relatively even sampling among *Corymbia* s.l. (i.e., (sub)genera *Angophora*, *Corymbia*, and *Blakella*; see Crisp et al., 2024 and Nicolle et al., 2025) and *Eucalyptus* (% of missing *Corymbia* s.l. samples at least 2/3 the % of missing *Eucalyptus* samples). We used this latter criterion because *Corymbia* s.l. samples were missing from gene trees more frequently than *Eucalyptus* samples, resulting in distorted branch lengths among *Corymbia* s.l. based on preliminary analyses. We then used TCMM to estimate branch lengths based on each of the 101 gene trees—regularizing estimates with the species tree branch lengths inferred via CASTLES under a regularization parameter of 0.01—and averaged the resulting distributions of estimates for each branch on the species tree after removing outliers. To detect and remove outliers, we discarded any estimates below $10^{-6}$ (a fairly consistent and conspicuous break point across all branch length distributions estimated via TCMM), then discarded any estimates above $Q_3 + 3\text{IQR}$, where $Q_3$ and IQR denote the 3rd quartile and interquartile range, respectively, of the distribution for each branch in the species tree (Arasti et al., 2024).

For absolute time calibrations using wLogDate (Mai and Mirarab, 2021), we assigned an age of 65 million years to the most recent common ancestor (MRCA) of all eucalpyts and an age of 50 million years to the MRCA of the *Eucalyptus* subgenera Eucalyptus and Symphyomyrtus. The latter calibration is based on ∼52 million year old Patagonian *Eucalyptus* fossils which exhibit Symphyomyrtus-like morphology (Gandolfo et al., 2011; Zamaloa et al., 2020), while the former reflects a mixture of results from previous phylogenetic analyses (Thornhill et al., 2012, 2015, 2019; Berger et al., 2016; Vasconcelos et al., 2017; Balbinott et al., 2022) as well as the general lack of reliable Myrtaceae fossils predating the very end of the Cretaceous period (Thornhill and Macphail, 2012). The final dated phylogeny largely agreed with previous results (Thornhill et al., 2019)—notable discrepancies include older divergences among the major sections of the *Eucalyptus* subgenus Symphyomyrtus, a slightly younger crown age for the *Eucalyptus* subgenus Eucalyptus, and a younger crown age for the (sub)genus *Angophora*.

Our final phylogeny included more than one tip for 61 out of the 318 unique eucalypt species it included. Because we could only collect species-level phenotypic data, we had to select one tip to represent each of these species for our final comparative analyses. Fortunately, duplicate tips formed exclusive monophyletic clades for 41 of these 61 species, meaning that selecting any given tip for these 41 species would result in an identical dated phylogeny.

We trimmed the phylogeny to one tip per species as follows. Monophyletic duplicates (41 out of 61 cases) were randomly trimmed. We grouped the remaining 20 non-monophyletic duplicates into non-overlapping clades. We removed all but a single tip for each species within a given clade—systematically trying all possible combinations of retained tips—and calculated quartet scores for the resulting pruned phylogenies using wASTRAL (Zhang and Mirarab, 2022). For each clade, we then selected the combination of retained tips resulting in the highest quartet score to make this tip “de-duplication” process as objective as possible. Unfortunately, the procedure yielded tied quartet scores for 9 out of 14 clades. We were able to resolve these ties in 4 of the remaining 9 clades based on taxonomic consensus among eucalypt researchers (Brooker et al., 2015; Nicolle, 2024). For the remaining 5 clades, we removed the most recently-diverged tip for a given species—both for consistency and because recent divergences tend to more strongly influence evolutionary rate estimates than older divergences (e.g., see Felsenstein, 2008).

#### *Additional details on trait data preparation/processing*

To detect and fix potential errors in our regular expression-based extractions of measurement data from EUCLID, we manually checked descriptions for any species associated with missing measurements, maximum measurements less than minimum ones, and/or abnormally high Mahalanobis distances (i.e., greater than the 90th percentile based on the corresponding $\chi^2$ distribution, Petersen and Pedersen, 2012). For the purpose of outlier detection, we imputed missing trait values and calculated Mahalanobis distances based on the empirical means and (co)variances among traits. Most traits exhibited substantial within-species variation/plasticity, and EUCLID—like most other botanical publications—tends to either provide ranges of measurements for a given trait (e.g., "petioles (0.8)1–1.5(1.9) cm"; the measurements in parentheses represent outliers) or only report maximum measurements (e.g., "Grows up to 8 meters in height"). To include as much data as possible in our comparative analyses while avoiding misleading comparisons between average trait values in some species and maximum values in others, we summarized trait values for each species based on maximum measurements (excluding outliers) rather than averages.

Adult leaf measurements were coded as missing for species known to retain juvenile/intermediate leaves in their mature crowns, despite crown leaf measurements often being reported in lieu of adult leaf measurements for these species (the only exceptions were species descriptions explicitly indicating that crown leaf measurements came from adult leaves only). Some species had stem-like structures (i.e., petioles, peduncles, and/or pedicels) described as either sessile or, equivalently, exhibiting a maximum length of 0 mm. Because measurements of these traits were consistently rounded to the nearest millimeter in EUCLID, we rounded up maximum lengths of 0 mm to 0.5 mm, allowing us to log-transform these measurements for comparative analysis. While such rounding could conceivably impact rate inferences, we ultimately found that our final comparative dataset did not include any species with sessile leaves or umbels (i.e., all species had petioles and peduncles that grew up to 1 mm or greater) and included relatively few species with sessile flower buds/fruit (10 and 14 out of 310 measurements for bud and fruit pedicel lengths, respectively).

Based on specific epithets, most eucalypt species represented in our phylogenetic data could be matched up to a single unique species in our trait data (250/318 species). Among the remaining species, 55 out of 68 simply matched to multiple subspecific taxa in our trait data. In these cases, we took the maximum trait measurement for each trait across all subspecies assigned to a given tip. Unmatched species were dropped from our phylogeny unless EUCLID considered them synonymous with another species in our trait data (provided the synonym was not already present in our phylogenetic data).

#### *Modeling maximum height evolution*

Height disparity varies markedly across eucalypt subclades. The *Eucalyptus* subgenus Eucalyptus and section Maidenaria (subgenus Symphyomyrtus) each encompass both the shortest and tallest known eucalypt species (roughly 1 and 100 m tall, respectively) despite originating just around 10–20 million years ago, while a much older clade, the (sub)genus *Blakella*, only contains species $\sim$10 to 50 m tall. Such patterns suggest rates of body size evolution are heterogeneous across eucalypts, and a homogeneous Brownian Motion process is unlikely to accurately model maximum height evolution across the clade.

We used the *evorates* package (Martin et al., 2023) to fit four models to the eucalypt maximum height data, including ones that allow rates of height evolution to vary across clades (see "Extending the algorithm: mapping under 'evolving rates' models" subsection in section "Technical Details of Continuous Stochastic Character Mapping Algorithm", page 50). Specifically, we fit: 1) a "trend" model whereby rates of height evolution exponentially decrease or increase over time (equivalent to an "early/late burst" model), 2) a "rate variance" model whereby different subclades gradually diverge in rates of height evolution, 3) a "full" model combing both the trend and rate variance models, and 4) a "null" model with constant rates (equivalent to a homogeneous Brownian Motion model). We fit all models using four independent Hamiltonian Monte Carlo chains each consisting of 2,000 iterations, discarding the first 1,000 as warmup for a total of 4,000 posterior samples. All chains adequately converged ($\hat{R} < 1.01$) and achieved sufficient effective sample sizes (effective sample sizes > 100 per chain) (Stan Development Team, 2019). Ultimately, we found high support for rate heterogeneity among clades under both the rate variance and full models (Savage-Dickey ratio < 0.01) but equivocal evidence for an overall trend in rates through time (posterior probability of increasing trend ~0.85 under both trend and full models). For these reasons, we based our final continuous stochastic character maps and associated empirical analyses on the rate variance model, but we briefly describe some notable differences in inferred evolutionary dynamics among these models in the following paragraph.

We inferred similar average rates of height evolution across all models, with geometric variance in height (i.e., variance in log height) increasing by about 0.02 per million years. However, the rate variance and full models infer evolutionary rates spanning from 0.001 to 0.3 across eucalypt lineages (for comparison, the trend model only inferred rates ranging from ~0.005 at the root of the phylogeny to 0.03 at the tips). Additionally, the more consistent rates inferred under null and trend models could not fully account for observed height disparity across related eucalypt species, resulting in both higher inferred levels of within-species variation/measurement error and extreme shrinkage of estimated heights towards clade-wide means (Fig. S23). Specifically, the null and trend models both inferred "tip error" (*sensu* Landis and Schraiber, 2017) geometric standard deviations of 50–60%, roughly suggesting "typical" height measurements for any given eucalypt species should vary around 6-fold overall. On the other hand, rate variance and full models inferred geometric standard deviations of about 30–40%, corresponding to expected height ranges a little over 3-fold. For reference, species height ranges reported in the literature typically exhibit 1.5 to 4-fold variation (e.g., Brooker et al., 2015; Falster et al., 2021). In general, the relatively small portion of eucalypt species reported to exhibit more extreme height variation around 6-fold or higher are known to adopt diminutive growth habits in exposed environments with nutrient-poor soils (e.g., *Corymbia deserticola*, *Eucalyptus calcicola*, *E. cordata*; Brooker et al., 2015).

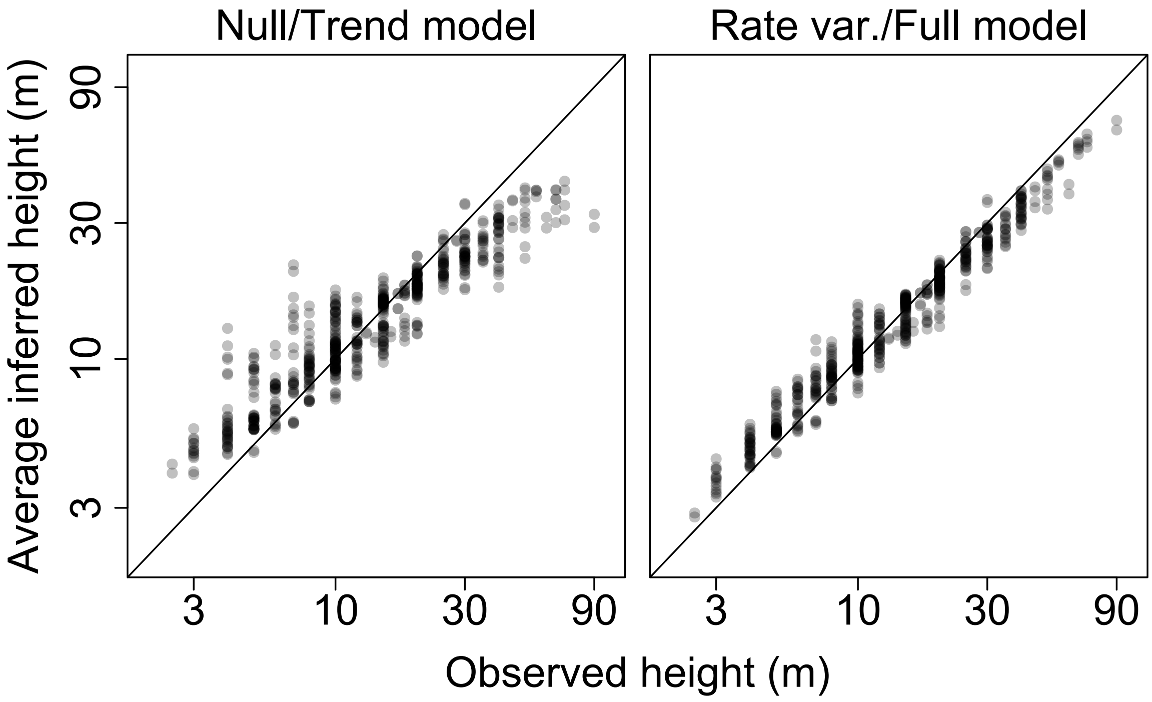

Figure S23. Plots depicting the relationship between observed maximum height measurements across eucalypts and average maximum heights inferred under different models fit via the *evorates* package (Martin et al., 2023). Note that the null and trend models—which did not allow eucalpyt clades to diverge in rates height evolution—yielded high estimates of intraspecific variation/measurement error, resulting in shrinkage of particularly low/high height measurements towards their cladewide means. By comparison, rate variance and full models—which did allow for divergence in rates of height evolution—yielded inferred heights that better agreed with observed measurements.

Ultimately, we generated 4,000 continuous stochastic character maps of eucalypt maximum heights and associated rates under the rate variance model (i.e., one map for each posterior sample), all with resolutions of 100 (see Fig. S22 in "Extending the algorithm: mapping under 'evolving rates' models" subsection in section "Technical Details of Continuous Stochastic Character Mapping Algorithm", page 50). To render downstream analyses more computationally manageable, we thinned the resulting sample of maps to every tenth map, the maximum thinning rate whereby effective sample sizes of mapped heights/rates at each time point remained above 100 (Geyer, 2011; Stan Development Team, 2019). To form the final set of height and dummy factor continuous stochastic character maps for our empirical analyses, we then generated 400 corresponding maps of dummy factor histories by simulating trait evolution with root values and mapped rates identical to those of the height maps.

*Optimization settings*

We used the same optimization algorithm settings, parameter boundaries, and Winsorization levels as the simulation study (see "Optimization settings" subsection in section "Further Details on Pipeline and Simulation Study", page 68). The one setting we altered was the number of optimization algorithm runs: to improve reliability of maximum likelihood estimates, we reran the optimization algorithm for each model fit 20 rather than 10 times. Additionally, to render numerical optimization more stable, we centered and rescaled our mapped height and dummy factor values to a mean of 0 and standard deviation of 1 across all maps. We also modified rate parameter functions to implicitly rescale the phylogeny to a height of 1 (i.e., dividing calculated rates by 65). Note that these parameters were back-transformed to proper units in all figures and tables.

Falster D., Gallagher R., Wenk E.H., Wright I.J., Indiarto D., Andrew S.C., Baxter C., Lawson J., Allen S., Fuchs A., Monro A., Kar F., Adams M.A., Ahrens C.W., Alfonzetti M., Angevin T., Apgaua D.M.G., Arndt S., Atkin O.K., Atkinson J., Auld T., Baker A., von Balthazar M., Bean A., Blackman C.J., Bloomfield K., Bowman D.M.J.S., Bragg J., Brodribb T.J., Buckton G., Burrows G., Caldwell E., Camac J., Carpenter R., Catford J.A., Cawthray G.R., Cernusak L.A., Chandler G., Chapman A.R., Cheal D., Cheesman A.W., Chen S.C., Choat B., Clinton B., Clode P.L., Coleman H., Cornwell W.K., Cosgrove M., Crisp M., Cross E., Crous K.Y., Cunningham S., Curran T., Curtis E., Daws M.I., DeGabriel J.L., Denton M.D., Dong N., Du P., Duan H., Duncan D.H., Duncan R.P., Duretto M., Dwyer J.M., Edwards C., Esperon-Rodriguez M., Evans J.R., Everingham S.E., Farrell C., Firn J., Fonseca C.R., French B.J., Frood D., Funk J.L., Geange S.R., Ghannoum O., Gleason S.M., Gosper C.R., Gray E., Groom P.K., Grootemaat S., Gross C., Guerin G., Guja L., Hahs A.K., Harrison M.T., Hayes P.E., Henery M., Hochuli D., Howell J., Huang G., Hughes L., Huisman J., Ilic J., Jagdish A., Jin D., Jordan G., Jurado E., Kanowski J., Kasel S., Kellermann J., Kenny B., Kohout M., Kooyman R.M., Kotowska M.M., Lai H.R., Laliberté E., Lambers H., Lamont B.B., Lanfear R., van Langevelde F., Laughlin D.C., Laugier-Kitchener B.A., Laurance S., Lehmann C.E.R., Leigh A., Leishman M.R., Lenz T., Lepschi B., Lewis J.D., Lim F.,

Liu U., Lord J., Lusk C.H., Macinnis-Ng C., McPherson H., Magallón S., Manea A., López-Martinez A., Mayfield M., McCarthy J.K., Meers T., van der Merwe M., Metcalfe D.J., Milberg P., Mokany K., Moles A.T., Moore B.D., Moore N., Morgan J.W., Morris W., Muir A., Munroe S., Nicholson Á., Nicolle D., Nicotra A.B., Niinemets Ü., North T., O'Reilly-Nugent A., O'Sullivan O.S., Oberle B., Onoda Y., Ooi M.K.J., Osborne C.P., Paczkowska G., Pekin B., Guilherme Pereira C., Pickering C., Pickup M., Pollock L.J., Poot P., Powell J.R., Power S.A., Prentice I.C., Prior L., Prober S.M., Read J., Reynolds V., Richards A.E., Richardson B., Roderick M.L., Rosell J.A., Rossetto M., Rye B., Rymer P.D., Sams M.A., Sanson G., Sauquet H., Schmidt S., Schönenberger J., Schulze E.D., Sendall K., Sinclair S., Smith B., Smith R., Soper F., Sparrow B., Standish R.J., Staples T.L., Stephens R., Szota C., Taseski G., Tasker E., Thomas F., Tissue D.T., Tjoelker M.G., Tng D.Y.P., de Tombeur F., Tomlinson K., Turner N.C., Veneklaas E.J., Venn S., Vesk P., Vlasveld C., Vorontsova M.S., Warren C.A., Warwick N., Weerasinghe L.K., Wells J., Westoby M., White M., Williams N.S.G., Wills J., Wilson P.G., Yates C., Zanne A.E., Zemunik G., and Ziemińska K. 2021. AusTraits, a curated plant trait database for the Australian flora. Sci Data 8:254.

Felsenstein J. 2008. Comparative methods with sampling error and within-species variation: Contrasts revisited and revised. Am Nat 171:713–725.

FitzJohn R.G., Maddison W.P., and Otto S.P. 2009. Estimating trait-dependent speciation and extinction rates from incompletely resolved phylogenies. Syst Biol 58:595–611.

Gandolfo M.A., Hermsen E.J., Zamaloa M.C., Nixon K.C., González C.C., Wilf P., Cúneo N.R., and Johnson K.R. 2011. Oldest known eucalyptus macrofossils are from south america. PLoS One 6:e21084.

Geyer C. 2011. Introduction to Markov chain Monte Carlo. *in* Handbook of Markov Chain Monte Carlo (S. Brooks, A. Gelman, G. L. Jones, and M. Xiao-Li, eds.). Chapman and Hall/CRC, Boca Raton, FL.

Goolsby E.W. 2017. Rapid maximum likelihood ancestral state reconstruction of continuous characters: A rerooting-free algorithm. Ecol Evol 7:2791–2797.

Han Y. and Molloy E.K. 2025. Improved robustness to gene tree incompleteness, estimation errors, and systematic homology errors with weighted TREE-QMC. Syst. Biol. Page syaf009.

Hansen T.F. and Bartoszek K. 2012. Interpreting the evolutionary regression: the interplay between observational and biological errors in phylogenetic comparative studies. Syst. Biol. 61:413–425.

Hansen T.F. and Martins E.P. 1996. Translating between microevolutionary process and macroevolutionary patterns: The correlation structure of interspecific data. Evolution 50:1404–1417.

Hassler G., Tolkoff M.R., Allen W.L., Ho L.S.T., Lemey P., and Suchard M.A. 2022. Inferring phenotypic trait evolution on large trees with many incomplete measurements. J Am Stat Assoc 117:678–692.

Hiscott G., Fox C., Parry M., and Bryant D. 2016. Efficient recycled algorithms for quantitative trait models on phylogenies. Genome Biol. Evol. 8:1338–1350.

Jhwueng D.C. 2021. Two Gaussian bridge processes for mapping continuous trait evolution along phylogenetic trees. Mathematics 9:1998.

Johnson S.G. 2021. The NLopt nonlinear-optimization package. Version 2.7.1.

Karatzas I. and Shreve S.E. 2004. Brownian motion and stochastic calculus. Graduate texts in mathematics 2 ed. Springer, New York, NY.

Landis M.J. and Schraiber J.G. 2017. Pulsed evolution shaped modern vertebrate body sizes. Proc Natl Acad Sci USA 114:13224–13229.

Landis M.J., Schraiber J.G., and Liang M. 2013. Phylogenetic analysis using Lévy processes: finding jumps in the evolution of continuous traits. Syst Biol 62:193–204.

Lartillot N., Phillips M.J., and Ronquist F. 2016. A mixed relaxed clock model. Philos Trans R Soc B 371:20150132.

Lepage T., Bryant D., Philippe H., and Lartillot N. 2007. A general comparison of relaxed molecular clock models. Mol Biol Evol 24:2669–2680.

Lewandowski D., Kurowicka D., and Joe H. 2009. Generating random correlation matrices based on vines and extended onion method. J Multivar Anal 100:1989–2001.

Liao H., Li Y., and Brooks G.P. 2017. Outlier impact and accommodation on power. J Mod Appl Stat Methods 16:261–278.

Mai U., Charvel E., and Mirarab S. 2024. Expectation-maximization enables phylogenetic dating under a categorical rate model. Syst. Biol. 73:823–838.

Mai U. and Mirarab S. 2021. Log transformation improves dating of phylogenies. Mol. Biol. Evol. 38:1151–1167.

Marsaglia G., Tsang W.W., and Wang J. 2003. Evaluating kolmogorov's distribution. J. Stat. Softw. 8:1–4.

Martin B.S., Bradburd G.S., Harmon L.J., and Weber M.G. 2023. Modeling the evolution of rates of continuous trait evolution. Syst Biol 72:590–605.

McElreath R. 2025. Statistical rethinking book package.

Minh B.Q., Schmidt H.A., Chernomor O., Schrempf D., Woodhams M.D., von Haeseler A., and Lanfear R. 2020. IQ-TREE 2: New models and efficient methods for phylogenetic inference in the genomic era. Mol Biol Evol 37:1530–1534.

Nicolle D. 2024. Classification of the eucalypts, genus *Eucalyptus*. version 7.1. `https://www.dn.com.au/Classification-Of-The-Eucalypts.pdf` accessed: 2025-8-10.

Nicolle D., Ritter M.K., Jones R.C., Phillips G.P., French M.E., Cumming R., and Bell S.A.J. 2025. The genus problem–*Eucalyptus* as a model system for minimising taxonomic disruption. Taxon 74:495–506.

Paradis E. and Schliep K. 2019. ape 5.0: An environment for modern phylogenetics and evolutionary analyses in R. Bioinformatics 35:526–528.

Pennell M.W., Eastman J.M., Slater G.J., Brown J.W., Uyeda J.C., FitzJohn R.G., Alfaro M.E., and Harmon L.J. 2014. geiger v2.0: An expanded suite of methods for fitting macroevolutionary models to phylogenetic trees. Bioinformatics 30:2216–2218.

Petersen K.B. and Pedersen M.S. 2012. The matrix cookbook. Version 20121115.

Revell L.J. 2012. phytools: an R package for phylogenetic comparative biology (and other things). Methods Ecol Evol 3:217–223.

Rowan T.H. 1990. Functional stability analysis of numerical algorithms. Ph.D. thesis Department of Computer Science, University of Texas at Austin, TX.

Smith S.A. and O'Meara B.C. 2012. treePL: divergence time estimation using penalized likelihood for large phylogenies. Bioinformatics 28:2689–2690.

Stan Development Team . 2019. Stan Modeling Language Users Guide and Reference Manual. Version 2.21.0.

Tabatabaee Y., Zhang C., Warnow T., and Mirarab S. 2023. Phylogenomic branch length estimation using quartets. Bioinformatics 39:i185–i193.

Thornhill A.H., Crisp M.D., Külheim C., Lam K.E., Nelson L.A., Yeates D.K., and Miller J.T. 2019. A dated molecular perspective of eucalypt taxonomy, evolution and diversification. Aust Syst Bot 32:29–48.

Thornhill A.H., Ho S.Y.W., Külheim C., and Crisp M.D. 2015. Interpreting the modern distribution of myrtaceae using a dated molecular phylogeny. Mol. Phylogenet. Evol. 93:29–43.

Thornhill A.H. and Macphail M. 2012. Fossil myrtaceous pollen as evidence for the evolutionary history of myrtaceae: A review of fossil myrtaceidites species. Rev. Palaeobot. Palynol. 176-177:1–23.

Thornhill A.H., Popple L.W., Carter R.J., Ho S.Y.W., and Crisp M.D. 2012. Are pollen fossils useful for calibrating relaxed molecular clock dating of phylogenies? a comparative study using myrtaceae. Mol. Phylogenet. Evol. 63:15–27.

Vasconcelos T.N.C., Proença C.E.B., Ahmad B., Aguilar D.S., Aguilar R., Amorim B.S., Campbell K., Costa I.R., De-Carvalho P.S., Faria J.E.Q., Giaretta A., Kooij P.W., Lima D.F., Mazine F.F., Peguero B., Prenner G., Santos M.F., Soewarto J., Wingler A., and Lucas E.J. 2017. Myrteae phylogeny, calibration, biogeography and diversification patterns: Increased understanding in the most species rich tribe of myrtaceae. Mol. Phylogenet. Evol. 109:113–137.

Welch J.J. and Waxman D. 2008. Calculating independent contrasts for the comparative study of substitution rates. J Theor Biol 251:667–678.

Yang Z. 2006. Computational Molecular Evolution. Oxford University Press, New York, NY.

Ypma J., Johnson S.G., Stamm A., Borchers H.W., Eddelbuettel D., Ripley B., Hornik K., Chiquet J., Adler A., Dai X., and Ooms J. 2022. nlotpr: R Interface to NLOPT. R package version 2.0.3.

Zamaloa M.C., Gandolfo M.A., and Nixon K.C. 2020. 52 million years old eucalyptus flower sheds more than pollen grains. Am. J. Bot. 107:1763–1771.

Zhang C. and Mirarab S. 2022. Weighting by gene tree uncertainty improves accuracy of quartet-based species trees. Mol. Biol. Evol. 39:msac215.
